# Meta-analysis and experimental evidence reveal no impact of *Nosema ceranae* infection on honeybee carbohydrate consumption

**DOI:** 10.1101/2025.02.11.637035

**Authors:** Monika Ostap-Chec, Weronika Antoł, Daniel Bajorek, Ewelina Berbeć, Dawid Moroń, Marcin Rapacz, Krzysztof Miler

## Abstract

Honeybees (*Apis mellifera*) are indispensable pollinators, essential for ecosystem stability and agricultural productivity. However, they face numerous challenges, including pathogens that threaten their survival and ecosystem services. Among these pathogens, *Nosema ceranae*, a microsporidian parasite, causes significant damage to the intestinal tract and induces energetic imbalances in an organism, posing a major threat to both individual bees and entire colonies. In response to infections, bees often engage in behavioural defenses, such as selecting foods with higher antibiotic properties. We hypothesized that bees infected with *N. ceranae* might compensate behaviourally by increasing their carbohydrate consumption. To test this hypothesis, we conducted a meta-analysis of existing studies comparing sugar consumption in healthy and infected bees, complemented by an experimental study. In our experiment, we measured sugar intake and quantified trehalose levels in the hemolymph, a key indicator of energy reserves. Both the meta-analysis and experimental results consistently showed no significant differences in sugar consumption between healthy and infected bees. Similarly, trehalose levels in the hemolymph remained comparable between the two groups. Our findings suggest that the infection caused by *N. ceranae* does not elicit compensatory feeding behavior in honeybees. Moreover, the meta-analysis revealed significant gaps in current research, particularly a lack of studies focusing on forager bees, which face the highest energetic demands among colony members. Our findings call for future studies on the energetic effects of nosemosis and studies conducted under natural or semi-natural conditions.

## Introduction

Parasites are among the most ubiquitous organisms on Earth, exerting both direct and indirect negative effects on their hosts [1]. Throughout coevolution, hosts have developed a range of strategies to defend themselves [2]. One of the first lines of defense against parasites lies in infection avoidance behavior. This defense is not always possible as, for example, intestinal parasites often enter the host accidentally, via ingestion of food or water. In response to parasitic infections, hosts may show changes in their appetite, dietary preferences, and foraging behavior [3, 4]. These changes may stem from adaptive responses of the host, host manipulation by parasites, or even be a by-product of infection [5]. Self-medication and compensatory feeding are among the most prevalent adaptive behavioral responses of hosts to gut infections. Self-medication involves the active selection of substances with therapeutic properties to combat infection [6, 7], while compensatory feeding focuses on replenishing resources lost due to infection or its treatment [8, 9]. These strategies can be difficult to distinguish, but the key distinction lies in the fact that self-medication involves diets that are harmful to healthy individuals [e.g. 10, 11], whereas compensatory feeding does not incur harmful effects for uninfected individuals [e.g. 12]. Thus, dietary compensation in infected individuals often involves the intake of familiar foods in different proportions or increased quantities [13, 14].

Honeybees (*Apis mellifera*) are critical pollinators that support both natural ecosystems and agricultural productivity [15]. However, these insects face numerous threats, among which parasitic infections stand out as a significant challenge to their health and survival [16]. Nosemosis, caused by *Nosema ceranae* and *N. apis*, is one of the most extensively studied gut infections in honeybees due to its severe impact on entire colony health. Given the critical role of honeybees in pollination, understanding and mitigating the effects of *Nosema* infection seems vital [17–19]. Transmission of *Nosema* spores is often accidental and difficult to avoid as it can occur via food exchange between individuals, collection of contaminated resources, or even sexually, between drones and queens [20–22]. Among the two agents responsible for nosemosis, *N. ceranae* has received greater attention due to its broader host range, year-round infection cycles with limited seasonal variation, and higher biotic potential across varying temperatures [23, 24].

*N. ceranae* completes its life cycle within the epithelial cells of the honeybee midgut, a critical area for nutrient absorption [25, 26]. Once spores reach the midgut, they multiply rapidly, exploiting host resources and triggering cell destruction, ultimately resulting in gut lesions, impaired absorption, and other effects (reviewed in [27, 28]). Although infected bees may not show overt symptoms, they often suffer from digestive disorders, lethargy, and shortened lifespans [19, 23, 29–32]. The physiological damage and chronic energetic stress from *N. ceranae* infection contribute to immunosuppression in honeybees [33–37]. Transcriptomic analyses have shown the upregulation of energy metabolism genes in the midgut cells of infected bees [38, 39], reflecting the parasite’s demand for host resources. Indeed, *Nosema* parasites, lacking mitochondria, rely on their hosts for essential energy sources, such as adenosine triphosphate (ATP), for their growth and reproduction [40]. The metabolic stress imposed by *N. ceranae* infection is further evidenced by decreased levels of hemolymph trehalose, a key sugar used for energy storage [41–43]. These energetic strains not only impair digestion but also affect thermoregulation and increase susceptibility to starvation [38, 44, 45]. Infected bees show prolonged foraging times and decreased flight frequency [46–49], likely due to the energetic burden of infection [43–45, 50]. Given that carbohydrates are the primary fuel for honeybee flight [43, 51], these effects can significantly reduce foraging success, as evidenced by reduced food stores in colonies heavily infected with *N. ceranae* [52].

Given the severe energetic strain and digestive impairment caused by *N. ceranae* infections, one might expect honeybees to exhibit adaptive changes in their feeding behavior to compensate for energy deficits. While honeybees are known to adjust their diets in response to various stressors, such as selecting nectar with higher antibiotic properties [53] or gathering propolis to reduce parasite loads [54], the impact of *N. ceranae* on carbohydrate intake remains inconclusive. Considering the unresolved questions regarding the effects of *N. ceranae* on honeybee feeding behavior, this study aimed to systematically investigate whether infection prompts an increase in carbohydrate consumption as a compensatory response. We conducted a systematic review and meta-analysis of existing literature to assess food intake patterns in healthy versus infected bees. Additionally, we conducted a laboratory experiment to measure food intake and trehalose levels in both healthy and infected bees. We hypothesised that infection with *N. ceranae* depletes the energy reserves of honeybees, and in response, bees may compensate for this loss by increasing their carbohydrate intake.

## Methods

### 1. Consumption experiment

#### 1.1. Experimental procedure

We conducted the experiment using four unrelated, queenright honeybee colonies, each in good overall condition and previously treated with oxalic acid against *Varroa destructor* in early spring. The bees were also examined for *Nosema* infection (see below).

We removed a single brood frame with capped cells, free of adult bees, from each colony and placed it in an incubator (KB53, Binder, Germany) set at 32°C overnight. The following day, newly emerged, one-day-old bees were collected and individually placed on Petri dishes. To increase feeding motivation, the bees were kept without food for approximately one hour.

The bees were then divided into two treatment groups: infected and control. Bees in the infected group were individually fed a 10 µl drop of 1M sucrose solution containing 100,000 *Nosema ceranae* spores, while bees in the control group received a 10 µl drop of 1M sucrose solution without spores. The bees were monitored for up to three hours, and those that fully consumed their solution were promptly transferred to the appropriate wooden cage prepared in advance. Bees that did not consume their solution were excluded from the experiment.

For each colony, we established 10 cages: five replicates containing infected bees and five containing control bees, with each cage housing 40 bees. This resulted in 200 bees per treatment group per colony, amounting to a total of 20 cages (1000 bees) per treatment group across all colonies. The cages were provided with *ad libitum* water and gravity feeders with 40% sucrose solution, then placed in an incubator (KB400, Binder, Germany) maintained at 32 °C. After an acclimation period of 24 hours, the initial number of living bees in each cage was recorded.

Each morning, the water and food in each cage were renewed and weighed both before and after renewal to calculate daily food consumption over the preceding 24 hours. Mortality was monitored daily, with all dead individuals removed. The experiment continued for 14 days. On the final day, three bees from each cage were frozen for later analysis to confirm infection status (control vs. infected). Additionally, hemolymph was collected from five bees per cage using the antennae method [55]. Each hemolymph sample was collected into a 10 μl end-to-end microcapillary and frozen at –20 °C for later analysis of trehalose content.

#### 1.2. Preparation of spores for infection and their genetic identification

To confirm the identity of the *N. ceranae* spores, we used PCR following the protocol by Berbeć et al. [56]. In brief, 50 µl of the spore suspension was incubated in TNES buffer (100 mM Tris-HCl pH 8.0, 5 mM EDTA, 0.3 % SDS, 200 mM NaCl) with 8 µl of proteinase K (10 mg/ml) for 2 hours at 56°C with shaking. After incubation, the DNA was extracted by centrifugation, followed by precipitation with an equal volume of 100% isopropanol. The DNA pellet was then washed twice with 70% ethanol, dried, and resuspended in 50 µl of nuclease-free water.

PCR amplification was conducted using species-specific primers targeting the rRNA genes of *Nosema* species, with a PCR Mix Plus kit containing PCR anti-inhibitors (A&A Biotechnology). The amplification conditions were as follows: an initial denaturation at 94°C for 3 minutes, followed by 35 cycles of 94°C for 30 seconds, 52°C for 30 seconds, and 72°C for 30 seconds. The following primer pairs were used: for *N. ceranae*: forward: 5’- CGGATAAAAGAGTCCGTTACC, reverse: 5’- TGAGCAGGGTTCTAGGGAT [57], and for *N. apis*: forward: 5’- CCATTGCCGGATAAGAGAGT, reverse: 5’- CCACCAAAAACTCCCAAGAG [58]. The PCR products were analyzed using gel electrophoresis to confirm the exclusive presence of *N. ceranae*.

The spore suspension used for experimental inoculation was freshly prepared on the same day the bees were fed. We sourced the spores from our stock population of infected honeybees, which were maintained in controlled incubator conditions to sustain the infection for spore harvesting. To prepare the suspension, we homogenized the digestive tracts of several infected individuals using a micropestle in distilled water. The mixture was centrifuged (Frontier 5306, Ohaus, Switzerland) at 6,000 G for 5 minutes, repeating this process three times. After each centrifugation, the supernatant was replaced with fresh distilled water.

The final supernatant was replaced with a 1M sucrose solution, and the concentration of spores was determined using a Bürker hemocytometer under a Leica DMLB light microscope equipped with phase contrast (PCM) and a digital camera. The final infection solution was adjusted to achieve a concentration of 100,000 spores per 10 µl by diluting with 1M sucrose solution.

#### 1.3. Verification of Nosema infection status

To confirm that bees exposed to *N. ceranae* spores were indeed infected and that bees in the control group remained uninfected, we assessed spore levels using qPCR quantification. We sampled three bees from each cage within both treatment groups across all colonies (as described above), resulting in a total of 15 infected and 15 control bees per colony (120 individuals overall).

Additionally, to ensure that the colonies used for the experiment were initially free from *N. ceranae* infection, we tested their infection status. For this, we collected two bees from the same brood frames used to gather one-day-old bees, froze them, and subsequently analyzed their spore presence using qPCR to confirm they were free of *Nosema* spores.

#### 1.4. DNA extraction and sample preparation for qPCR

For DNA extraction, the abdomens of individual bees were cut using sterile instruments and placed into 2 ml cryovials, each containing 800 µl dH_2_O. The samples were homogenized using a Bead Ruptor ELITE homogenizer (Omni International) with a combination of “small” (0.5 mm) and “big” (2.8 mm) ceramid beads (Omni International).

From each homogenate, a 200 µl aliquot was taken for DNA extraction following a solution-based protocol with Nuclei Lysis Solution and Protein Precipitation Solution (Promega). The resulting DNA pellets were resuspended in 100 µl of 1× TE buffer for further analysis.

#### 1.5. Quantification of N. ceranae load by qPCR

To quantify the parasite load, we amplified a 65 bp fragment of the *N. ceranae Hsp70* gene using primers designed by Cilia et al. [59]. Since the *Hsp70* gene exists as a single copy per spore, the measured copy number can be directly translated into the spore load per individual [59, 60].

The original DNA extracts were diluted 10× for qPCR which was performed using a CFX96 Touch Real-Time PCR Detection System (Bio-Rad) in a 20 µl reaction mix containing: 5 µl of DNA template, 10 µl of SsoAdvanced^TM^ Universal SYBR^®^ Green Supermix (Bio-Rad), target primers at a final concentration of 0.2 µM (see Table 1 for primer sequences), and ddH_2_O up to a total volume of 20 µl. The cycling conditions were as follows: an initial denaturation at 98 °C for 3 minutes, followed by 40 cycles of 98 °C for 15 seconds and 60 °C for 30 seconds.

**Table 1.**
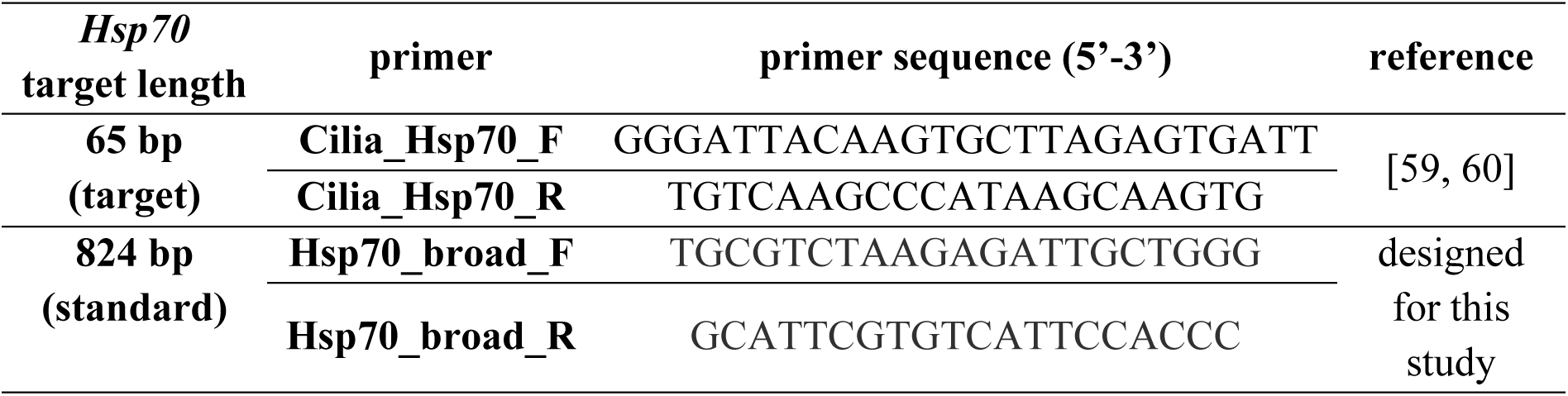
Primers used in qPCR for *Hsp70* amplification and absolute quantification.

For absolute quantification, a purified PCR product of a broader 824 bp *Hsp70* fragment was used as a standard. The fragment was amplified using newly designed primers (see Table 1) in a 25 µl reaction mix containing: 2 µl of DNA extract from an infected bee, 12.5 µl of DreamTaq^TM^ Hot Start PCR Master Mix, 0.4 µM of each primer, and ddH_2_O to a final volume of 25 µl. The cycling conditions were: initial denaturation at 95 °C for 3 minutes, followed by 40 cycles of 95 °C for 30 seconds, 58 °C for 30 seconds, and 72 °C for 60 seconds, with a final elongation at 72 °C for 5 minutes.

To increase yield, products from multiple reactions were pooled and subjected to agarose gel electrophoresis. The target band was then excised and purified using the ZymoClean Gel Recovery Kit. The concentration of the purified product was measured using a Qubit Broad Range Assay. The copy number was calculated based on the concentration and fragment length according to Qiagen guidelines [61] for absolute quantification.

A standard curve for each plate was prepared using the purified product, with a dilution series covering a dynamic range of 8 logs (from 10^−1^ to 10^6^ copies). Standards were run in triplicate and samples were in duplicate on each qPCR plate. The method demonstrated a sensitivity of 6.20 × 10^−1^ copies/µl (equivalent to 2 480 copies/bee or 3.39 log copies/bee), which was the lowest concentration in the dilution series with high reproducibility and strong linearity (R^2^ ≥ 0.996). PCR efficiencies for the standard curve between 90 and 110% were accepted. To confirm specificity, a melting curve analysis was performed at the end of each run, covering the temperature range of 65-95 °C in 0.5 °C increments, with a dwell time of 5 seconds per step.

#### 1.6. Calculation of spore load

To determine the spore load, the mean starting quantity of the template for each duplicate based on the standard curve was calculated. The results were expressed as the number of *Hsp70* copies/µl, which corresponds to the number of spores (following Cilia et al. [59]). Given that: DNA extract was diluted 10× for the reaction, the total DNA extract volume was 100 µl, and that the DNA extraction was performed on 1/4 of the original homogenate volume, the initial spore count was multiplied by 4000. Finally, the values were log_10_-transformed to obtain the spore load as the log number of spores per bee.

#### 1.7. Hemolymph trehalose analysis

Trehalose levels were measured using a trehalose assay kit (K-TREH, Neogen, USA), following the manufacturer’s protocol. Each hemolymph sample (1 μl) was diluted 200-fold with distilled water. Samples with less than 1 µl were discarded, as volumes below this threshold were considered not precise enough for quantitative analysis.

For the assay, 20 µl of each diluted sample was analyzed in duplicate. The analysis was performed in batches (consecutive plates). The increase in absorbance, resulting from enzymatic reaction products, was measured at 340 nm using the Multiskan FC microplate reader (Thermo Scientific, USA). Trehalose concentrations were calculated using a calibration curve ranging from 0.00625 to 0.4 g/l and adjusted by a factor of 200 to reflect the original concentration in the undiluted samples.

Due to the detection limit of the assay kit (1 g/l after correcting for dilution), any values below this threshold were considered 0. In total, we analyzed 84 samples from the control group and 65 from the infected group.

#### 1.8. Statistical analysis

All statistical analyses were performed using R [62]. Daily sucrose solution consumption was calculated based on the differences in the feeder weight. The per capita consumption (mg/bee/day) was estimated by dividing the total consumption by the number of live bees on a given day, yielding the average amount of food consumed per individual per day. To analyze this consumption data, we employed a generalized additive model (GAM, [63] using the *mgcv* package [64, 65]). The model, with a Gaussian distribution and an identity link function, was used to examine the relationship between sugar consumption and time (days) for both the control and infected groups. The colony and cage (replicate) nested within the colony were included as random effects.

For mortality analysis, we used Cox mixed-effects regression with the *survival* and *lme4* packages [65, 66]. The model included group (control vs. infected) as a fixed effect, with colony and cage (replicate) nested within the colony as random effects.

Trehalose levels were analyzed using a linear mixed-effect model fitted with the *lmer* function from the *lme4* package [66]. The model included group (control vs. infected) as a fixed factor, while colony, cage (replicate), and plate (data analysis batch) were treated as random effects. Data were transformed by applying square root to achieve a normal distribution of residuals. The significance of the fixed effect was tested using the *anova* function (stats package) [62].

### 2. Systematic review and meta-analysis

We conducted a systematic review and meta-analysis following the Preferred Reporting Items for Systematic Reviews and Meta-Analyses (PRISMA) guidelines [68]. The review protocol was not pre-registered.

#### 2.1. Eligibility criteria

We established the following eligibility criteria for studies to be included in the systematic review and meta-analysis: (i) studies using worker honeybees (*A. mellifera*), (ii) laboratory experiments comparing workers exposed to *N. ceranae* spores with workers not exposed to *Nosema*, and (iii) studies reporting food consumption measurements for both groups over time.

#### 2.2. Data sources and search

An electronic search was conducted on January 16, 2024, using the Web of Science and Scopus databases. The search phrase used was “nosema and apis” in the topic, along with a forward search (i.e. documents citing one or more works from the list) refined by the same phrase. References were de-duplicated using Mendeley and then Zotero. We filtered out articles with “review” or “meta-analysis” in the title and further searched for all studies containing “ceranae” in the title, abstract, or keywords. Titles and abstracts of these articles were reviewed for potential inclusion. Two independent investigators assessed the eligibility of the articles, with studies marked as potentially eligible by either investigator being included for full-text review.

#### 2.3. Extraction of study details and study exclusion

From the eligible studies, we extracted the following data: (i) mean and standard deviation (SD) of food consumption for both groups, (ii) number of replicates per group, (iii) duration of consumption measurement (days), (iv) concentration of sucrose solution used, (v) age of the inoculated workers, (vi) method of inoculation (individual or collective), and (vii) inoculation dose (spores per individual). Data extraction was performed by one investigator.

In cases where data were reported in a different format, appropriate conversions were applied to standardize measurements (mg/bee/day). As such, in two cases, SD was calculated from the reported standard error (SE) using the formula SD = SE × √_sample size_. In one case, SD was obtained from the range of values (range divided by 4). In five cases, the mean and SD were calculated based on the median, quartiles, and maximum/minimum values using the method by Wan et al. [69]. In seven cases, the data were reported as volume (μL), so we converted it to mass (mg) using the following formula: consumed mg = sucrose solution concentration used in the study × 1.39 g/ml × consumed volume reported × 1000 mg/g. If not available in the text, the data were extracted from figures using Web Plot Digitizer [70] for maximum accuracy.

Studies were excluded if: (i) there was no comparison between infected and control groups (e.g. consumption measured only for the infected), (ii) food consumption was measured but not reported in a usable form (e.g. data not shown), (iii) complete data extraction was not possible due to missing information (e.g. unknown worker age, infection dose, mode of infection, sucrose solution concentration).

In two studies where the number of replicates varied between groups and was not specified, we assumed the lowest reported replicate count for both groups.

#### 2.4. Risk of bias

The risk of bias was assessed by one investigator based on: (i) similarity of baseline characteristics between control and infected groups (low bias if similar, high bias if not, and unclear if differentiating factors were of unknown effect), and (ii) whether contamination was measured and reported (low bias if measured and reported, high bias if not, and unclear if not mentioned).

#### 2.5. Meta-analysis

The meta-analysis was conducted in R [62] using the *metafor* [71] and *orchaRd* packages [72]. We calculated standardized mean differences (SMDs) in consumption between infected and control groups for each study. A multivariate random-effects model was fitted using the *rma.mv* function, with random effects at the study level. Results were visualized using an orchard plot, and heterogeneity was assessed with Q and I^2^ statistics. An influence analysis was conducted to identify outliers based on Cook’s distance, and the analysis was repeated with influential studies removed. A meta-regression was conducted to examine potential sources of heterogeneity, using the following moderators: age at inoculation, inoculation dose, duration of study, and mode of inoculation, as fixed effects.

#### 2.6. Publication bias

Publication bias was assessed visually using a funnel plot by examining the degree of effect asymmetry (plotting standard error against SMDs, see Supplementary Material 1) and statistically using Egger’s test.

#### 2.7. Quality of evidence

The strength of the evidence was evaluated by one investigator using the Grading of Recommendations Assessment, Development, and Evaluation (GRADE) guidelines [73]. The quality assessment considered study design, risk of bias, publication bias, inconsistencies, indirectness, imprecision, and the effect size in the included studies.

## Results

### 1. Consumption experiment

The analysis revealed that infection with *N. ceranae* spores had no significant effect on sugar consumption in honeybees (estimate ± SE, control: 45.53 ± 1.54 mg/bee/day, infected: 44.68 ± 1.54 mg/bee/day; t = -0.953, p = 0.341). However, sugar consumption patterns over time displayed significant non-linearity in both groups (control: F = 31.84, p < 0.001, edf = 6.44; infected: F = 28.90, p < 0.001, edf = 6.53). The high edf’s (> 2) indicate strong non-linear trends (Fig. 1) characterized by a steep increase in consumption during the initial days, followed by a more stable intake for the remaining time. The model accounted for 34% of the variance in sugar consumption.

**Fig. 1.**
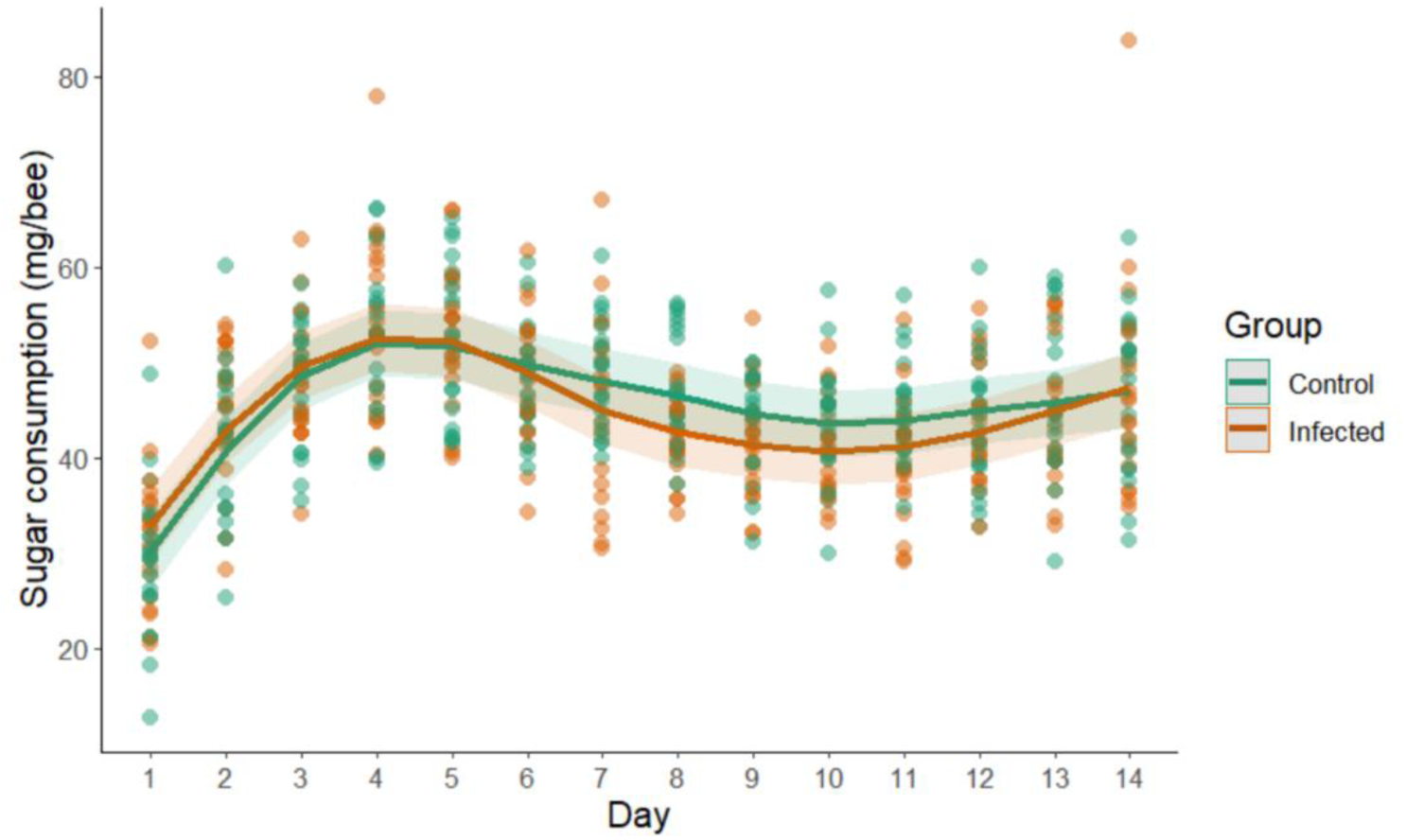
Sucrose consumption (mg/bee) in control and infected bees over 14 days. There are no significant differences between groups. Lines represent generalized additive model predictions and shaded areas show 95% CI.

Mortality rates differed significantly between the two groups. The hazard ratio, representing the risk of death, was 2.87 times higher (SE: 0.12) for infected bees compared to control individuals (z = 8.84, p < 0.001, Fig. 2).

**Fig. 2.**
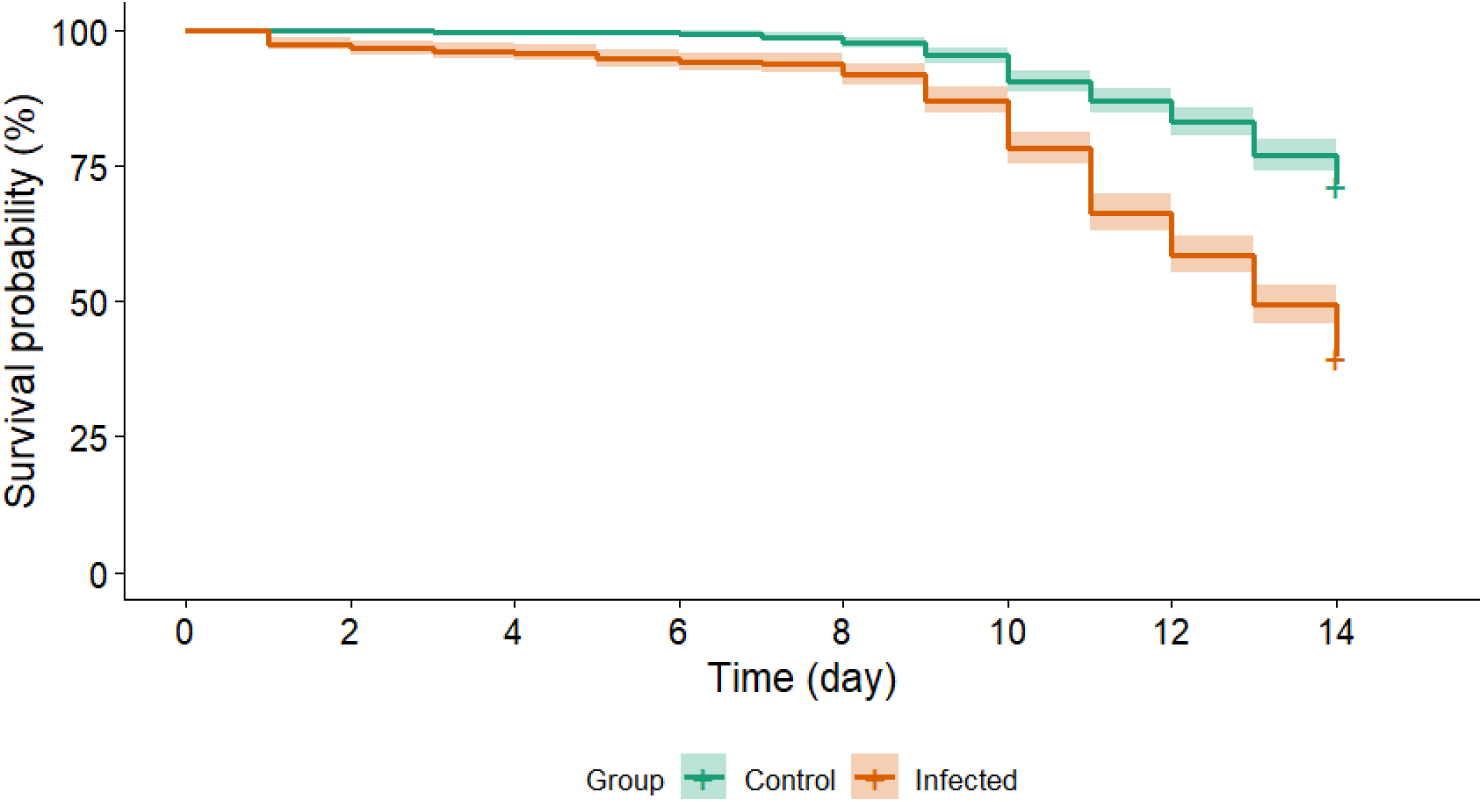
Survival plot for control (healthy) and *Nosema*-infected bees over the 14 days. Each group consisted of 20 cages, with an initial number of individuals of 37-40 bees per cage. Lines represent Cox regression predictions and shaded areas show 95% CI.

Spore loads differed significantly between the groups. In the infected group (N = 60), the mean log spore count per bee (± SD) was 6.58 ± 1.15, whereas the control group (N = 60) showed a much lower spore level of 2.59 ± 1.12, comparable to background spore counts in bees sampled directly from the frames before the experiment (N = 12, 2.05 ± 1.27). These background levels were consistent with previous studies [74–76]. To account for the possible cross-contamination with spores during DNA extraction, extraction blanks (N = 4) were analyzed, showing minimal spore levels (1.83 ± 1.27 log spores in the whole homogenate volume).

At the end of the 14-day experiment, no significant differences were found in hemolymph trehalose levels between the *Nosema*-infected and control bees (F = 0.5568, p = 0.48, Fig. 3). The mean trehalose level in the control group was 27.86 ± 0.62 g/l (estimate ± SE, back-transformed from the square root) and in the infected group – 24.54 ± 0.63 g/l.

**Fig. 3.**
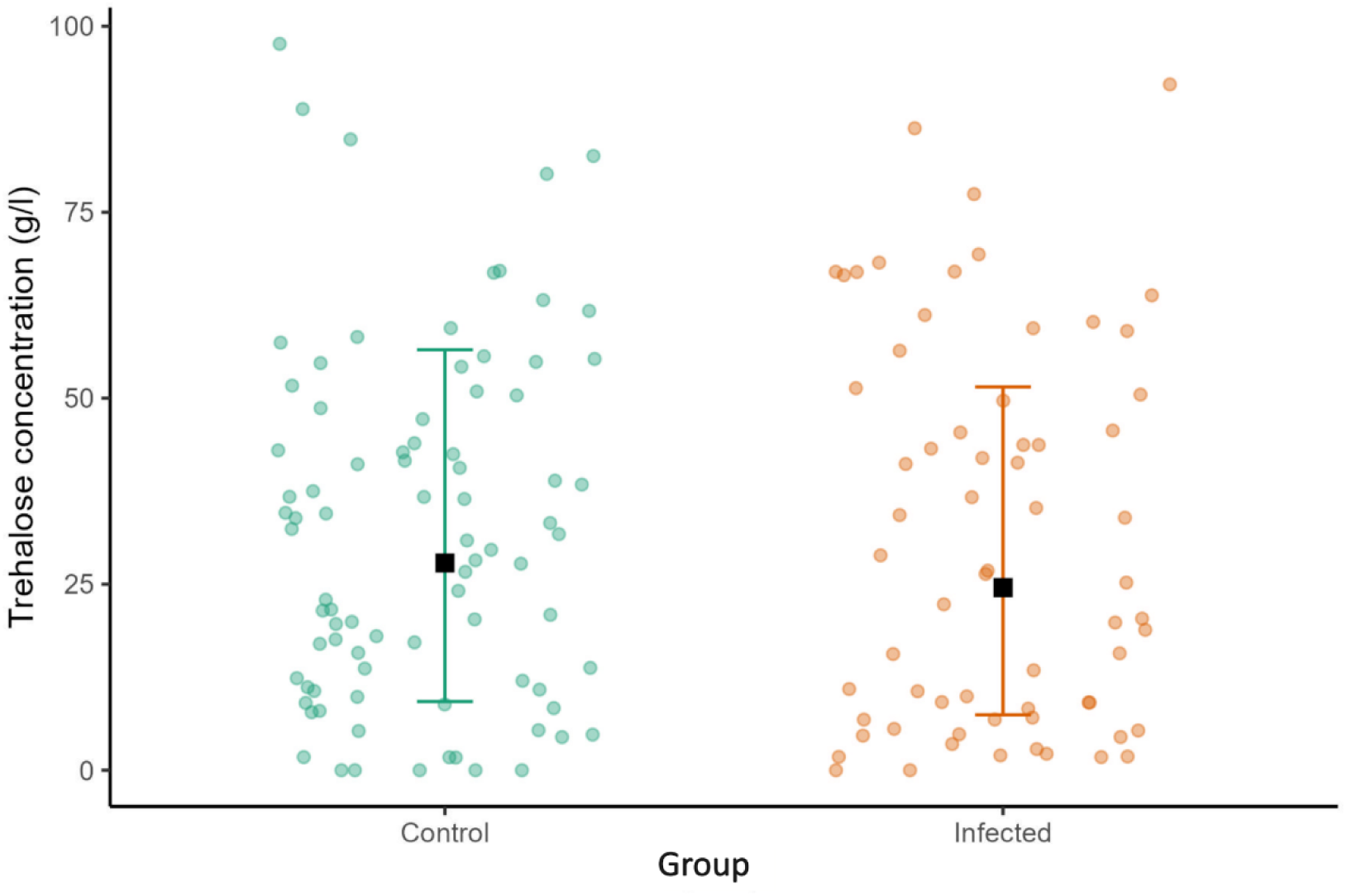
Trehalose levels in control and *Nosema*-infected groups of bees at the end of the 14-day experiment. Black squares indicate model estimates and whiskers show confidence intervals, both back-transformed from the square root. Individual datapoints are shown as dots.

### 2. Systematic review and meta-analysis

#### 2.1. Study characteristics

Our search yielded 8259 documents (2478 from Web of Science and 5781 from Scopus). After excluding two retracted articles and de-duplicating, 4481 documents remained. Filtering out reviews and meta-analyses removed an additional 114 articles and among those remaining, 819 documents contained “ceranae” in the title, abstract, or keywords. From these, we retrieved 207 potentially eligible articles for full-text review (listed in Supplementary Material 2), ultimately identifying 15 eligible studies from 14 documents. The current experiment was added as the 16th study (Fig. 4).

**Fig. 4.**
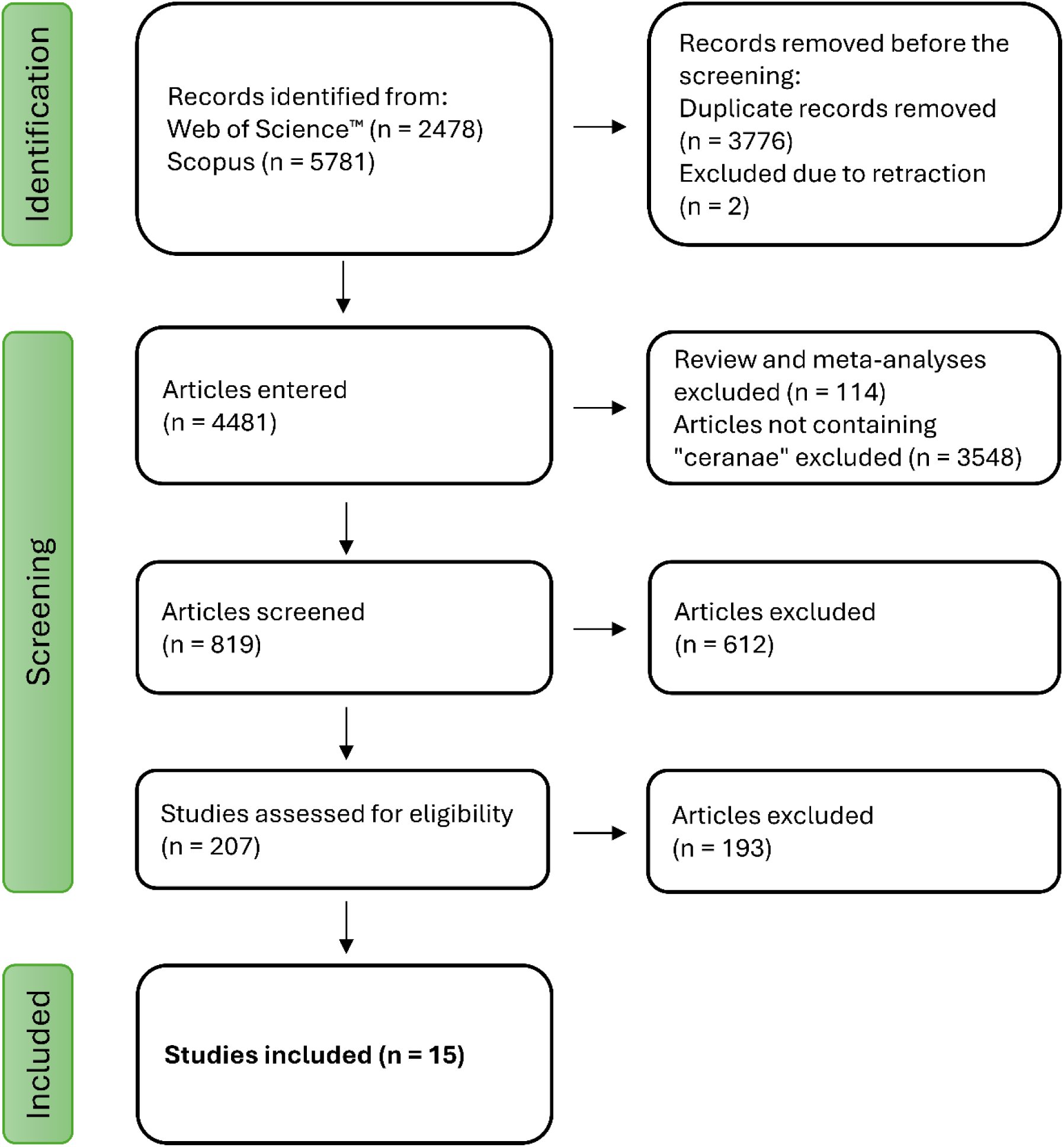
PRISMA flow diagram of the study selection process.

Among the included studies (Table 2), only two used more than 10 replicates, with most relying on three to four replicates (56% of studies). The average duration of sucrose consumption measurement was two weeks, with 81% of studies using a 50% sucrose solution as food. Most studies inoculated one-day-old workers, while only two used bees of at least 10 days of age. Individual inoculation was more common (69% of studies), typically using a dose of 100,000 spores (56 % of studies).

**Table 2.** Overview of studies included in the systematic review and meta-analysis.

| Reference | Study | Number of replicates (cages per group) | Duration of analysis (days) | Sucrose solution concentration (% w/v) | Age of workers when inoculated | Method of inoculation | Inoculation dose of spores |
| --- | --- | --- | --- | --- | --- | --- | --- |
| Almasri et al. 2021 [77] | 1 | 7 | 23 | 60 | 1 | Individual | 100,000 |
| Al Naggar & Baer 2019 [78] | 2 | 4 | 10 | 35 | 1 | Individual | 10,000 |
| Balbuena et al. 2023 [79] | 3 | 3 | 15 | 50 | 1 | Collective | 100,000 |
| Berbec et al. 2022 [56] | 4 | 4 | 7 | 50 | 2 | Collective | 100,000 |
|  | 5 | 4 | 7 | 50 | 10 | Collective | 100,000 |
| De la Mora et al. 2023 [80] | 6 | 6 | 18 | 50 | 1 | Individual | 100,000 |
| Ferguson et al. 2018 [81] | 7 | 8 | 19 | 50 | 1 | Individual | 10,000 |
| Garrido et al. 2023 [76] | 8 | 3 | 10 | 50 | 1 | Collective | 50,000 |
| Paris et al. 2017 [82] | 9 | 3 | 22 | 50 | 5 | Individual | 100,000 |
| Peghaire et al. 2020 [83] | 10 | 3 | 16 | 50 | 2 | Collective | 10,000 |
| Straub et al. 2020 [84] | 11 | 12 | 14 | 50 | 1 | Collective | 10,000 |
| Tritschler et al. 2017 [85] | 12 | 6 | 14 | 50 | 3 | Collective | 100,000 |
| Uruena et al. 2023 [86] | 13 | 3 | 14 | 50 | 6 | Individual | 100,000 |
| Vidau et al. 2011 [87] | 14 | 9 | 10 | 50 | 5 | Individual | 125,000 |
| Williams et al. 2014 [88] | 15 | 4 | 7 | 50 | 1 | Individual | 35,000 |
| **Current study** | 16 | 20 | 14 | 40 | 1 | Individual | 100,000 |

#### 2.2. Risk of bias

The overall risk of bias was low. Baseline characteristics of bees were similar between groups in most studies, and only two studies failed to report contamination appropriately.

#### 2.3. Meta-analysis

The random-effects meta-analysis showed that infection with *N. ceranae* had no significant effect on sucrose consumption (SMD = 0.0765 [95% CI: -0.3073, 0.4602], z = 0.3905, p = 0.6962) (Fig. 5). There was high heterogeneity (Q = 332.40, df = 15, p < 0.0001, I^2^ = 94.91%).

**Fig. 5.**
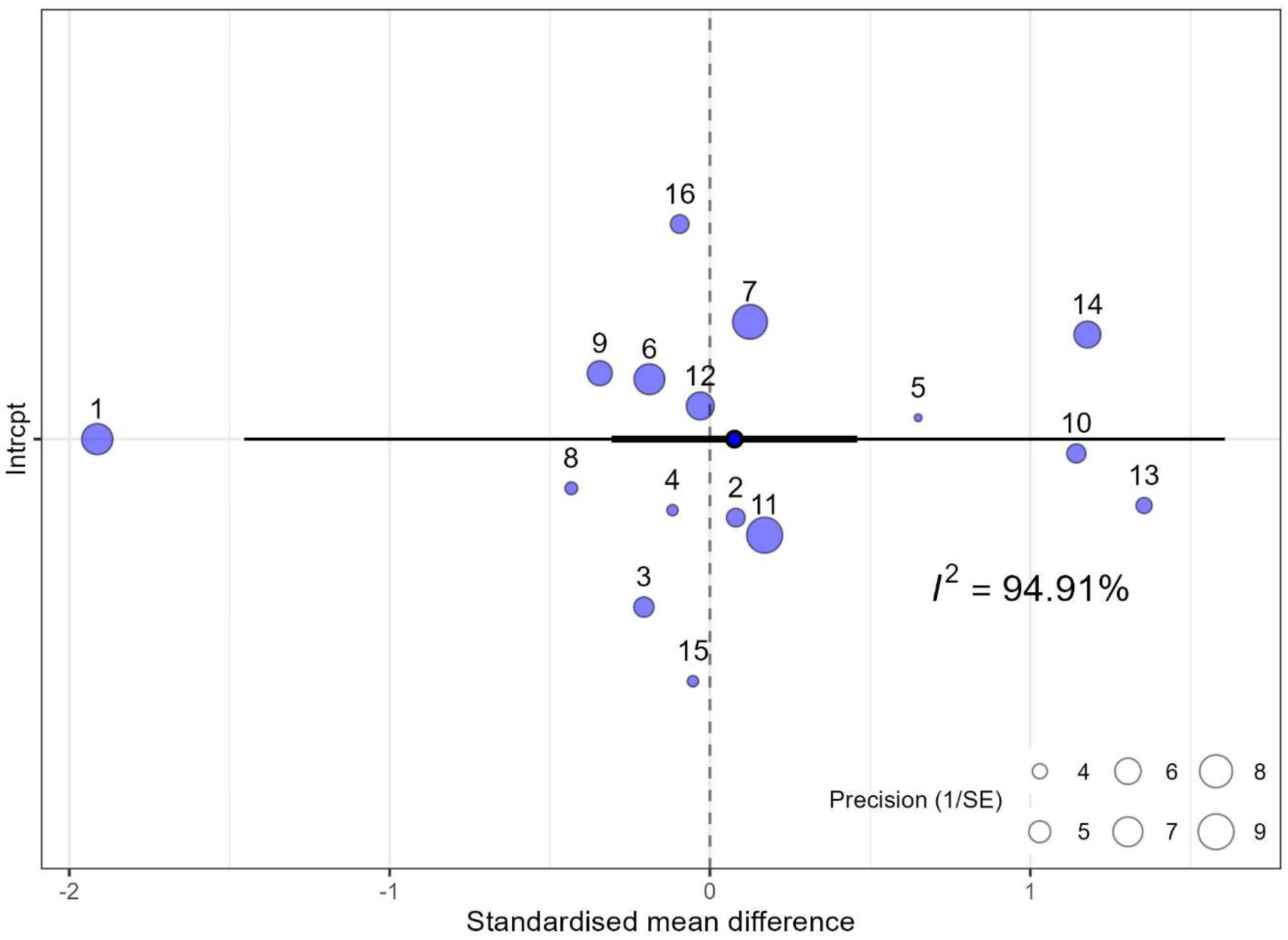
Orchard plot of standardized mean differences in sucrose consumption. Dots represent individual effect sizes, scaled by their precision, and numbers refer to the study number in the meta-analysis. The model estimate is shown as a dot outlined in black, with thin black lines indicating the 95% confidence intervals (CI) and thick black lines indicating the 95% prediction intervals (PI). The I² statistic estimates heterogeneity across studies.

Influence analysis identified four outliers (studies number 1, 10, 13, and 14). Removing these outliers reduced heterogeneity but it remained substantial (Q = 19.25, df = 11, p = 0.0568; I^2^ = 40.40%). After outlier removal, there was still no significant effect of infection on consumption rates (SMD = 0.031 [95% CI: -0.165, 0.104], z = 0.447, p = 0.655).

#### 2.4. Exploration of data heterogeneity

Subgroup meta-analysis comparing individual versus collective *Nosema* inoculation methods revealed no significant differences (Q = 0.16, df = 1, p = 0.692), with similar levels of heterogeneity (individual: 96.9%, collective: 79.4%). Among the moderators assessed (inoculation dose, age at inoculation, and duration of study), only inoculation age had a significant effect (estimate ± SE = 0.174 ± 0.077, p = 0.024). This suggests that studies using older bees reported higher effect sizes. In contrast, dose and duration had no significant impact (p > 0.05).

#### 2.5. Publication bias

Visual inspection of the funnel plot and Egger’s test indicated no publication bias (t = 0.845, p = 0.413).

#### 2.6. Quality of evidence

All included studies were randomized trials. Given the low risk of bias, absence of publication bias, and lack of inconsistencies or indirectness, the overall quality of evidence was rated as high. However, there was some concern regarding imprecision due to variability in consumption measures, which had to be standardized, potentially introducing errors. The summarized quality of evidence assessment is presented in Table 3.

**Table 3.** Summarized quality of evidence assessment.

| Number of studies | Overall risk of bias | Inconsistencies | Indirectness | Imprecision | Publication bias | Main effect | Dose response | Quality |
| --- | --- | --- | --- | --- | --- | --- | --- | --- |
| 16 | Low | Not serious | Not serious | Serious | Not likely | No effect | No effect | High |

## Discussion

Both our experimental findings and the meta-analysis suggest that infection with *N. ceranae* does not impact sugar consumption in honeybees. Among works marked as potentially eligible in our systematic review, several were excluded because no data was shown, even though they measured sucrose consumption. Notably, however, some of those studies reported no differences in consumption between the control and infected groups of workers [89–92]. Four other studies, two excluded for the usage of unspecified sucrose concentration [93, 94], one for unspecified dose and mode of *Nosema* inoculation [95], and another for the usage of mixed *Nosema* species for inoculation [96] all demonstrated no effects of infection on consumption. Although formally excluded from our analysis, these studies strongly support the conclusion that nosemosis does not affect sucrose consumption. This result is somewhat surprising since the system of *Nosema* and honeybees seems a potential candidate for pronounced dietary compensation due to its strong relevance for the energy budget of the host [28].

A recent meta-analysis by Mrugała et al. [97] addresses the issue of how parasitic infections influence host feeding behavior. The analyses, carried out across all taxonomic groups, demonstrate that, on average, infected hosts consume about 25% less food compared to uninfected ones, with considerable variability across taxa. Notably, the study highlights that parasitic infections can increase the variability of host consumption rates by an average of 25%, suggesting that the effects are highly context-dependent and influenced by factors such as host type or age [97]. In more general terms, the impact of parasitism on feeding behavior may depend on the specific host-parasite relationship and ecological context. Our meta-analysis revealed high heterogeneity between studies (I² = 94.9%). Neither the dose of inoculation nor the number of assessment days significantly influenced effect sizes. Additionally, no differences were observed between the individual and collective inoculation methods. Among the included moderators, only the age at inoculation had a meaningful effect. Specifically, studies that utilized older bees showed that infected individuals reacted to nosemosis the strongest and by consuming more sugar [56, 86]. This suggests that older bees may exhibit dietary compensation. Notably, none of the abovementioned studies that were formally excluded from the meta-analysis and demonstrated no differences between healthy and infected individuals utilized older bees. This indirectly supports the idea of a stronger effect of nosemosis in older individuals. However, more studies that contrast bees of different ages are needed. Interestingly, this issue seems to be confounded by the fact that it is not the age of bees per se that seems to be important, but the time of their inoculation. Specifically, in studies with relatively long duration of the analysis, in which the age of bees at the end of the experiment was high, but with inoculation at the very beginning of their life, infected bees did not consume more sugar than healthy ones [77, 81]. In any case, there is a notable lack of research focusing on forager-age bees.

It is well-documented that bees adjust their behavior to cope with illness by selecting food with specific benefits during infection, like nectar with higher antibiotic potential [53], gathering propolis to counter parasite levels [54], or choosing higher-quality pollen when infected with *Nosema* [81]. Our findings did not support the hypothesis that bees infected with *N. ceranae* exhibit increased sugar consumption. It might be that *Nosema* infection does not induce sufficient energetic stress to warrant an increase in sugar intake. This conclusion is supported by the lack of significant changes in trehalose levels observed in our study. This finding aligns with those of Li et al. [98], who reported that *Nosema* infection does not alter glycogen levels in bees, a primary glucose storage molecule. Kurze et al. [99] found that in *Nosema*-tolerant bees, the infection did not affect fructose, glucose, or trehalose levels, while in the sensitive strains, trehalose levels were altered (i.e. decreased). In contrast, studies by Mayack and Naug [43] and Aliferis et al. [50] reported significant changes in carbohydrate and amino acid levels, including reduced levels of fructose, L-proline, and cryoprotectants such as sorbitol and glycerol, suggesting higher energetic stress in *Nosema*-infected bees. However, these studies differ considerably from ours in key aspects. While our study, along with those by Li et al. [98] and Kurze et al. [99], was conducted in controlled laboratory conditions, Mayack et al. [43] and Aliferis et al. [50] investigated wild-caught foragers. The higher energy expenditure associated with flight and foraging activities in these bees likely introduces a different metabolic response compared to laboratory-reared bees. This further supports the idea of more pronounced energetic effects of *Nosema* infection in older or otherwise more sensitive bees.

Importantly, the energetic stress in *Nosema*-infected honeybees may involve more than just changes in sugar metabolism. Studies have shown that as *Nosema* infection progresses, protein synthesis is reduced, indicating a disruption of protein metabolism [50, 98, 100, 101]. Additionally, infected bees show increased respirometric activity and lipid loss, suggesting that lipids may be used to meet the heightened energy demands caused by the infection [98]. Notably, Gilbert et al. [102] report severe depletion of lipids in the honeybee fat body 14 days after *N. ceranae* infection. In summer bees, this depletion mirrors the lipid loss associated with aging. However, seasonal variations in lipid metabolism are significant: fall bees, with larger lipid reserves, experience a 50% reduction in their lipid stores when infected with *N. ceranae*. Furthermore, recent studies on microRNA (miRNA) expression and metabolomic analyses further highlight the extensive metabolic disruptions caused by *N. ceranae* infection [103, 104]. These findings highlight that it is important to incorporate various measures of responses to infection, ideally accounting for various factors such as bee age, nutritional status, seasonal timing and environmental conditions.

In conclusion, our experiment and meta-analysis provide consistent and reliable evidence that *N. ceranae* infection does not appear to significantly increase sugar consumption in honeybees. Moreover, the experiment revealed no significant difference in trehalose levels between infected and healthy bees, suggesting that their energetic status may be similar. However, it is important to note that both our study and the included meta-analysis have limited representation of older bees, particularly foragers. This highlights a significant gap in research. Furthermore, it is crucial to complement existing studies with field research conducted in natural or semi-natural conditions to gain a more comprehensive understanding of the metabolic responses to *Nosema* infection in honeybees.

## Acknowledgments

We thank Magdalena Migalska for her valuable advice during the development and optimization of the qPCR protocol.

