## Supplementary Material 1 for "Meta-analysis and experimental evidence reveal no impact of *Nosema ceranae* infection on honeybee carbohydrate consumption"

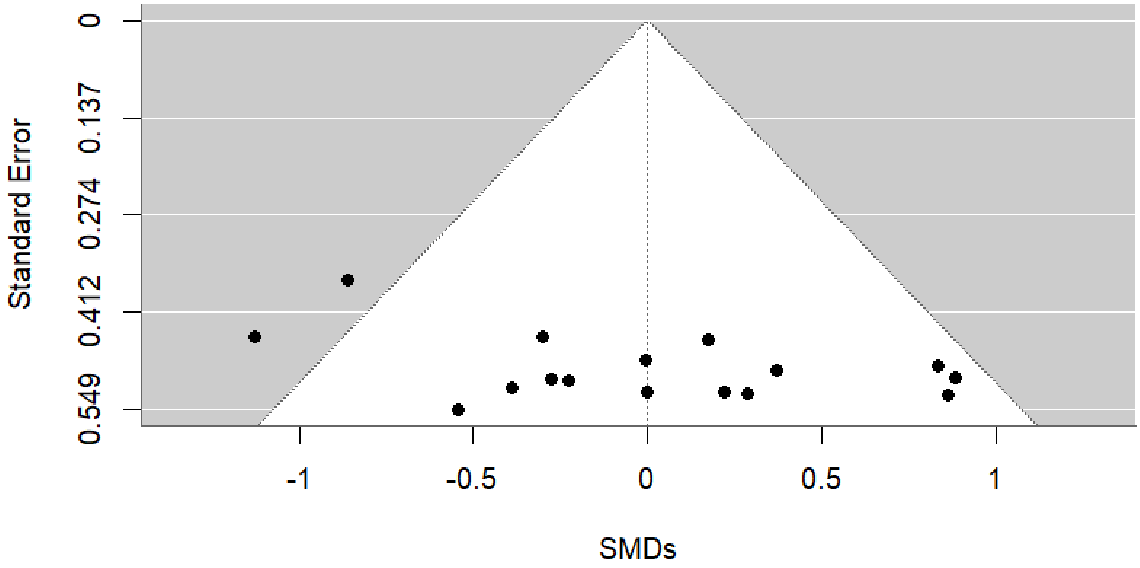


**Supplementary Figure 1**. Funnel plot (standard error plotted against SMDs) for the studies included in the meta-analysis. Each dot represents one study.
