## Supplementary Material 2 for "Meta-analysis and experimental evidence reveal no impact of *Nosema ceranae* infection on honeybee carbohydrate consumption"

23. Blot, N., Veillat, L., Rouzé, R., & Delatte, H. (2019). Glyphosate, but not its metabolite AMPA, alters the honeybee gut microbiota. PLoS ONE, 14(4). https://doi.org/10.1371/journal.pone.0215466
24. Borges, D., Guzman-Novoa, E., & Goodwin, P. H. (2020). Control of the microsporidian parasite Nosema ceranae in honey bees (Apis mellifera) using nutraceutical and immuno-stimulatory compounds. PLoS ONE, 15(1). https://doi.org/10.1371/journal.pone.0227484
25. Borges, D., Guzman-Novoa, E., & Goodwin, P. H. (2021). Effects of prebiotics and probiotics on honey bees (Apis mellifera) infected with the microsporidian parasite nosema ceranae. Microorganisms, 9(3), 1–16. https://doi.org/10.3390/microorganisms9030481
26. Braglia, C., Alberoni, D., Porrini, M. P., Garrido, M. P., Baffoni, L., & Di Gioia, D. (2021). Screening of dietary ingredients against the honey bee parasite nosema ceranae. Pathogens, 10(9). https://doi.org/10.3390/pathogens10091117
27. Bravo, J., Carbonell, V., Sepúlveda, B., Delporte, C., Valdovinos, C. E., Martín-Hernández, R., & Higes, M. (2017). Antifungal activity of the essential oil obtained from Cryptocarya alba against infection in honey bees by Nosema ceranae. Journal of Invertebrate Pathology, 149, 141–147. https://doi.org/10.1016/j.jip.2017.08.012
28. Buczek, K., Deryło, K., Kutyła, M., Rybicka-Jasińska, K., Gryko, D., Borsuk, G., Rodzik, B., & Trytek, M. (2020). Impact of protoporphyrin lysine derivatives on the ability of nosema ceranae spores to infect honeybees. Insects, 11(8), 1–12. https://doi.org/10.3390/insects11080504
29. Burnham, A. J., De Jong, E., Jones, J. A., & Lehman, H. K. (2020). North American Propolis Extracts From Upstate New York Decrease Nosema ceranae (Microsporidia) Spore Levels in Honey Bees (Apis mellifera). Frontiers in Microbiology, 11. https://doi.org/10.3389/fmicb.2020.01719
30. Campbell, J., Kessler, B., Mayack, C., & Naug, D. (2010). Behavioural fever in infected honeybees: Parasitic manipulation or coincidental benefit? Parasitology, 137(10), 1487–1491. https://doi.org/10.1017/S0031182010000235
31. Castelli, L., Balbuena, S., Branchiccela, B., Zunino, P., Liberti, J., Engel, P., & Antúnez, K. (2021). Impact of chronic exposure to sublethal doses of glyphosate on honey bee immunity, gut microbiota and infection by pathogens. Microorganisms, 9(4). https://doi.org/10.3390/microorganisms9040845
32. Castelli, L., Branchiccela, B., Garrido, M., Invernizzi, C., Porrini, M., Romero, H., Santos, E., Zunino, P., & Antúnez, K. (2020). Impact of Nutritional Stress on Honeybee Gut Microbiota, Immunity, and Nosema ceranae Infection. Microbial Ecology, 80(4), 908–919. https://doi.org/10.1007/s00248-020-01538-1
33. Chaimanee, V., & Chantawannakul, P. (2015). Infectivity of Nosema ceranae isolates from different hosts and immune response in honey bees Apis mellifera and Apis cerana. Journal of Apicultural Research, 54(3), 200–206. https://doi.org/10.1080/00218839.2016.1144975
34. Chaimanee, V., Chantawannakul, P., Chen, Y., Evans, J. D., & Pettis, J. S. (2012). Differential expression of immune genes of adult honey bee (Apis mellifera) after inoculated by Nosema ceranae. Journal of Insect Physiology, 58(8), 1090–1095. https://doi.org/10.1016/j.jinsphys.2012.04.016
35. Chaimanee, V., Kasem, A., Nuanjohn, T., Boonmee, T., Siangsuepchart, A., Malaithong, W., Sinpoo, C., Disayathanoowat, T., & Pettis, J. S. (2021). Natural

extracts as potential control agents for Nosema ceranae infection in honeybees, Apis mellifera. Journal of Invertebrate Pathology, 186. https://doi.org/10.1016/j.jip.2021.107688
36. Chaimanee, V., Pettis, J. S., Chen, Y., Evans, J. D., Khongphinitbunjong, K., & Chantawannakul, P. (2013). Susceptibility of four different honey bee species to Nosema ceranae. Veterinary Parasitology, 193(1–3), 260–265. https://doi.org/10.1016/j.vetpar.2012.12.004
37. Chantaphanwattana, T., Houdelet, C., Sinpoo, C., Voisin, S. N., Bocquet, M., Disayathanoowat, T., Chantawannakul, P., & Bulet, P. (2023). Proteomics and Immune Response Differences in Apis mellifera and Apis cerana Inoculated with Three Nosema ceranae Isolates. Journal of Proteome Research, 22(6), 2030–2043. https://doi.org/10.1021/acs.jproteome.3c00095
38. Charbonneau, L. R., Hillier, N. K., Rogers, R. E. L., Williams, G. R., & Shutler, D. (2016). Effects of Nosema apis, N. ceranae, and coinfections on honey bee (Apis mellifera) learning and memory. Scientific Reports, 6. https://doi.org/10.1038/srep22626
39. Chen, H., Du, Y., Xiong, C., Zheng, Y., Chen, D., & Guo, R. (2019). A comprehensive transcriptome data of normal and Nosema ceranae-stressed midguts of Apis mellifera ligustica workers. Data in Brief, 26. https://doi.org/10.1016/j.dib.2019.104349
40. Chen, H., Fan, X., Zhang, W., Ye, Y., Cai, Z., Zhang, K., Zhang, K., Fu, Z., Chen, D., & Guo, R. (2022). Deciphering the CircRNA-Regulated Response of Western Honey Bee (Apis mellifera) Workers to Microsporidian Invasion. Biology, 11(9). https://doi.org/10.3390/biology11091285
41. Chen, X., Wang, S., Xu, Y., Gong, H., Wu, Y., Chen, Y., Hu, F., & Zheng, H. (2021). Protective potential of Chinese herbal extracts against microsporidian Nosema ceranae, an emergent pathogen of western honey bees, Apis mellifera L. Journal of Asia-Pacific Entomology, 24(1), 502–512. https://doi.org/10.1016/j.aspen.2019.08.006
42. Cho, R. M., Kogan, H. V., Elikan, A. B., & Snow, J. W. (2022). Paromomycin Reduces Vairimorpha (Nosema) ceranae Infection in Honey Bees but Perturbs Microbiome Levels and Midgut Cell Function. Microorganisms, 10(6). https://doi.org/10.3390/microorganisms10061107
43. Choppin, M., & Lach, L. (2022). A novel bee host cannot detect a microbial parasite, in contrast to its original host. Insectes Sociaux, 69(2–3), 289–292. https://doi.org/10.1007/s00040-022-00860-w
44. Costa, C., Lodesani, M., & Maistrello, L. (2010). Effect of thymol and resveratrol administered with candy or syrup on the development of Nosema ceranae and on the longevity of honeybees (Apis mellifera L.) in laboratory conditions. Apidologie, 41(2), 141–150. https://doi.org/10.1051/apido/2009070
45. Damiani, N., Fernández, N. J., Porrini, M. P., Gende, L. B., Álvarez, E., Buffa, F., Brasesco, C., Maggi, M. D., Marcangeli, J. A., & Eguaras, M. J. (2014). Laurel leaf extracts for honeybee pest and disease management: Antimicrobial, microsporicidal, and acaricidal activity. Parasitology Research, 113(2), 701–709. https://doi.org/10.1007/s00436-013-3698-3
46. De la Mora, A., Morfin, N., Tapia-Rivera, J. C., Macías-Macías, J. O., Tapia-González, J. M., Contreras-Escareño, F., Petukhova, T., & Guzman-Novoa, E. (2023). The Fungus Nosema ceranae and a Sublethal Dose of the Neonicotinoid Insecticide

Thiamethoxam Differentially Affected the Health and Immunity of Africanized Honey Bees. Microorganisms, 11(5). https://doi.org/10.3390/microorganisms11051258
47. de Mattos, I. M., Soares, A. E. E., & Tarpy, D. R. (2018). Mitigating effects of pollen during paraquat exposure on gene expression and pathogen prevalence in Apis mellifera L. Ecotoxicology, 27(1), 32–44. https://doi.org/10.1007/s10646-017-1868-2
48. de Oliveira, A. H., Souza, A. M. D. C., de Resende, M. T. C. S., Carneiro, L. S., de Oliveira, J. F., Serra, R. S., & Serrão, J. E. (2023). The peritrophic matrix delays Nosema ceranae infection in the honey bee Apis mellifera midgut. Physiological Entomology, 48(2–3), 61–67. https://doi.org/10.1111/phen.12402
49. De Piano, F. G., Maggi, M., Pellegrini, M. C., Cugnata, N. M., Szawarski, N., Buffa, F., Negri, P., Fuselli, S. R., Audisio, C. M., & Ruffinengo, S. R. (2017). Effects of lactobacillus johnsonii AJ5 metabolites on nutrition, nosema ceranae development and performance of Apis Mellifera L. Journal of Apicultural Science, 61(1), 93–104. https://doi.org/10.1515/JAS-2017-0007
50. Di Pasquale, G., Salignon, M., Le Conte, Y., Belzunces, L. P., Decourtye, A., Kretzschmar, A., Suchail, S., Brunet, J.-L., & Alaux, C. (2013). Influence of Pollen Nutrition on Honey Bee Health: Do Pollen Quality and Diversity Matter? PLoS ONE, 8(8). https://doi.org/10.1371/journal.pone.0072016
51. Doublet, V., Labarussias, M., de Miranda, J. R., Moritz, R. F. A., & Paxton, R. J. (2015). Bees under stress: Sublethal doses of a neonicotinoid pesticide and pathogens interact to elevate honey bee mortality across the life cycle. Environmental Microbiology, 17(4), 969–983. https://doi.org/10.1111/1462-2920.12426
52. Doublet, V., Natsopoulou, M. E., Zschiesche, L., & Paxton, R. J. (2015). Within-host competition among the honey bees pathogens Nosema ceranae and Deformed wing virus is asymmetric and to the disadvantage of the virus. Journal of Invertebrate Pathology, 124, 31–34. https://doi.org/10.1016/j.jip.2014.10.007
53. Doublet, V., Paxtona, R. J., McDonnell, C. M., Dubois, E., Nidelet, S., Moritz, R. F. A., Alaux, C., & Le Conte, Y. (2016). Brain transcriptomes of honey bees (Apis mellifera) experimentally infected by two pathogens: Black queen cell virus and Nosema ceranae. GENOMICS DATA, 10, 79–82. https://doi.org/10.1016/j.gdata.2016.09.010
54. Duguet, J., Zuñiga, F., & Martínez, J. (2022). Antifungal activity of "HO21-F", a formulation based on Olea europaea plant extract, in honey bees infected with Nosema ceranae. Journal of Invertebrate Pathology, 193. https://doi.org/10.1016/j.jip.2022.107801
55. Dussaubat, C., Brunet, J.-L., Higes, M., Colbourne, J. K., Lopez, J., Choi, J.-H., Martín-Hernández, R., Botías, C., Cousin, M., McDonnell, C., Bonnet, M., Belzunces, L. P., Moritz, R. F. A., Le Conte, Y., & Alaux, C. (2012). Gut pathology and responses to the microsporidium Nosema ceranae in the honey bee Apis mellifera. PLoS ONE, 7(5). https://doi.org/10.1371/journal.pone.0037017
56. Dussaubat, C., Sagastume, S., Gómez-Moracho, T., Botías, C., García-Palencia, P., Martín-Hernández, R., Le Conte, Y., & Higes, M. (2013). Comparative study of Nosema ceranae (Microsporidia) isolates from two different geographic origins. Veterinary Microbiology, 162(2–4), 670–678. https://doi.org/10.1016/j.vetmic.2012.09.012

69. Garrido, P. M., Porrini, M. P., Antúnez, K., Branchiccela, B., Martínez-Noël, G. M. A., Zunino, P., Salerno, G., Eguaras, M. J., & Ieno, E. (2016). Sublethal effects of acaricides and Nosema ceranae infection on immune related gene expression in honeybees. Veterinary Research, 47(1). https://doi.org/10.1186/s13567-016-0335-z
70. Gherman, B. I., Denner, A., Bobiş, O., Dezmirean, D. S., Mărghitaş, L. A., Schlüns, H., Moritz, R. F. A., & Erler, S. (2014). Pathogen-associated self-medication behavior in the honeybee Apis mellifera. Behavioral Ecology and Sociobiology, 68(11), 1777–1784. https://doi.org/10.1007/s00265-014-1786-8
71. Giacomini, J. J., Leslie, J., Tarpy, D. R., Palmer-Young, E. C., Irwin, R. E., & Adler, L. S. (2018). Medicinal value of sunflower pollen against bee pathogens. Scientific Reports, 8(1). https://doi.org/10.1038/s41598-018-32681-y
72. Glavinic, U., Blagojevic, J., Ristanic, M., Stevanovic, J., Lakic, N., Mirilovic, M., & Stanimirovic, Z. (2022). Use of Thymol in Nosema ceranae Control and Health Improvement of Infected Honey Bees. Insects, 13(7). https://doi.org/10.3390/insects13070574
73. Glavinic, U., Dzogovic, D., Jelisic, S., Ristanic, M., Zorc, M., Aleksic, N., & Stanimirovic, Z. (2023). OXIDATIVE STATUS OF HONEY BEES INFECTED WITH NOSEMA CERANAE MICROSPORIDIUM AND SUPPLEMENTED WITH AGARICUS BISPORUS MUSHROOM EXTRACT. Veterinarski Glasnik, 77(1), 35–50. https://doi.org/10.2298/VETGL220715013G
74. Glavinic, U., Rajkovic, M., Vunduk, J., Vejnovic, B., Stevanovic, J., Milenkovic, I., & Stanimirovic, Z. (2021). Effects of agaricus bisporus mushroom extract on honey bees infected with nosema ceranae. Insects, 12(10). https://doi.org/10.3390/insects12100915
75. Glavinic, U., Stankovic, B., Draskovic, V., Stevanovic, J., Petrovic, T., Lakic, N., & Stanimirovic, Z. (2017). Dietary amino acid and vitamin complex protects honey bee from immunosuppression caused by Nosema ceranae. PLoS ONE, 12(11). https://doi.org/10.1371/journal.pone.0187726
76. Glavinic, U., Stevanovic, J., Ristanic, M., Rajkovic, M., Davitkov, D., Lakic, N., & Stanimirovic, Z. (2021). Potential of fumagillin and agaricus blazei mushroom extract to reduce nosema ceranae in honey bees. Insects, 12(4). https://doi.org/10.3390/insects12040282
77. Glavinic, U., Tesovnik, T., Stevanovic, J., Zorc, M., Cizelj, I., Stanimirovic, Z., & Narat, M. (2019). Response of adult honey bees treated in larval stage with prochloraz to infection with Nosema ceranae. PeerJ, 2019(2). https://doi.org/10.7717/peerj.6325
78. Goblirsch, M., Huang, Z. Y., & Spivak, M. (2013). Physiological and Behavioral Changes in Honey Bees (Apis mellifera) Induced by Nosema ceranae Infection. PLoS ONE, 8(3). https://doi.org/10.1371/journal.pone.0058165
79. Gregorc, A., Jurišić, S., & Sampson, B. (2020). Hydroxymethylfurfural affects caged honey bees (Apis mellifera carnica). Diversity, 12(1). https://doi.org/10.3390/d12010018
80. Gregorc, A., Silva-Zacarin, E. C. M., Carvalho, S. M., Kramberger, D., Teixeira, E. W., & Malaspina, O. (2016). Effects of Nosema ceranae and thiametoxam in Apis mellifera: A comparative study in Africanized and Carniolan honey bees. Chemosphere, 147, 328–336. https://doi.org/10.1016/j.chemosphere.2015.12.030
81. He, N., Zhang, Y., Le Duan, X., Li, J. H., Huang, W.-F., Evans, J. D., Degrandi-Hoffman, G., Chen, Y. P., & Huang, S. K. (2021). Rna interference-mediated

knockdown of genes encoding spore wall proteins confers protection against nosema ceranae infection in the european honey bee, apis mellifera. Microorganisms, 9(3), 1–17. https://doi.org/10.3390/microorganisms9030505
82. Hendriksma, H. P., Bain, J. A., Nguyen, N., & Nieh, J. C. (2020). Nicotine does not reduce Nosema ceranae infection in honey bees. Insectes Sociaux, 67(2), 249–259. https://doi.org/10.1007/s00040-020-00758-5
83. Higes, M., García-Palencia, P., Botías, C., Meana, A., & Martín-Hernández, R. (2010). The differential development of microsporidia infecting worker honey bee (Apis mellifera) at increasing incubation temperature. Environmental Microbiology Reports, 2(6), 745–748. https://doi.org/10.1111/j.1758-2229.2010.00170.x
84. Higes, M., García-Palencia, P., Martín-Hernández, R., & Meana, A. (2007). Experimental infection of Apis mellifera honeybees with Nosema ceranae (Microsporidia). Journal of Invertebrate Pathology, 94(3), 211–217. https://doi.org/10.1016/j.jip.2006.11.001
85. Higes, M., García-Palencia, P., Urbieta, A., Nanetti, A., & Martín-Hernández, R. (2020). Nosema apis and Nosema ceranae Tissue Tropism in Worker Honey Bees (Apis mellifera). Veterinary Pathology, 57(1), 132–138. https://doi.org/10.1177/0300985819864302
86. Higes, M., Juarranz, A., Dias-Almeida, J., Lucena, S., Botías, C., Meana, A., García-Palencia, P., & Martín-Hernández, R. (2013). Apoptosis in the pathogenesis of Nosema ceranae (Microsporidia: Nosematidae) in honey bees (Apis mellifera). Environmental Microbiology Reports, 5(4), 530–536. https://doi.org/10.1111/1758-2229.12059
87. Higes, M., Nozal, M. J., Alvaro, A., Barrios, L., Meana, A., Martín-Hernández, R., Bernal, J. L., & Bernal, J. (2011). The stability and effectiveness of fumagillin in controlling Nosema ceranae (Microsporidia) infection in honey bees (Apis mellifera) under laboratory and field conditions. Apidologie, 42(3), 364–377. https://doi.org/10.1007/s13592-011-0003-2
88. Higes, M., Rodríguez-García, C., Gómez-Moracho, T., Meana, A., Bartolomé, C., Maside, X., Barrios, L., & Martín-Hernández, R. (2016). Survival of honey bees (Apis mellifera) infected with Crithidia mellificae spheroid forms (Langridge and McGhee: ATCC® 30254TM) in the presence of Nosema ceranae. Spanish Journal of Agricultural Research, 14(3). https://doi.org/10.5424/sjar/2016143-8722
89. Holt, H. L., Aronstein, K. A., & Grozinger, C. M. (2013). Chronic parasitization by Nosema microsporidia causes global expression changes in core nutritional, metabolic and behavioral pathways in honey bee workers (Apis mellifera). BMC Genomics, 14(1). https://doi.org/10.1186/1471-2164-14-799
90. Hosaka, Y., Kato, Y., Hayashi, S., Nakai, M., Barribeau, S. M., & Inoue, M. N. (2021). The effects of Nosema ceranae (Microspora: Nosematidae) isolated from wild Apis cerana japonica (Hymenoptera: Apidae) on Apis mellifera. Applied Entomology and Zoology, 56(3), 311–317. https://doi.org/10.1007/s13355-021-00735-9
91. Houdelet, C., Arafah, K., Bocquet, M., & Bulet, P. (2022). Molecular histoproteomy by MALDI mass spectrometry imaging to uncover markers of the impact of Nosema on Apis mellifera. Proteomics, 22(9). https://doi.org/10.1002/pmic.202100224
92. Houdelet, C., Sinpoo, C., Chantaphanwattana, T., Voisin, S. N., Bocquet, M., Chantawannakul, P., & Bulet, P. (2021). Proteomics of Anatomical Sections of the Gut of Nosema-Infected Western Honeybee (Apis mellifera) Reveals Different Early

Responses to Nosema spp. Isolates. Journal of Proteome Research, 20(1), 804–817. https://doi.org/10.1021/acs.jproteome.0c00658
93. Hu, Y.-T., Wu, T.-C., Yang, E.-C., Wu, P.-C., Lin, P.-T., & Wu, Y.-L. (2017). Regulation of genes related to immune signaling and detoxification in Apis mellifera by an inhibitor of histone deacetylation. Scientific Reports, 7. https://doi.org/10.1038/srep41255
94. Huang, Q., Chen, Y., Wang, R. W., Schwarz, R. S., & Evans, J. D. (2015). Honey bee microRNAs respond to infection by the microsporidian parasite Nosema ceranae. Scientific Reports, 5. https://doi.org/10.1038/srep17494
95. Huang, Q., & Evans, J. D. (2020). Targeting the honey bee gut parasite Nosema ceranae with siRNA positively affects gut bacteria. BMC Microbiology, 20(1). https://doi.org/10.1186/s12866-020-01939-9
96. Huang, Q., Lariviere, P. J., Powell, J. E., & Moran, N. A. (2023). Engineered gut symbiont inhibits microsporidian parasite and improves honey bee survival. Proceedings of the National Academy of Sciences of the United States of America, 120(25). https://doi.org/10.1073/pnas.2220922120
97. Huang, Q., Li, W., Chen, Y., Retschnig-Tanner, G., Yanez, O., Neumann, P., & Evans, J. D. (2019). Dicer regulates Nosema ceranae proliferation in honeybees. Insect Molecular Biology, 28(1), 74–85. https://doi.org/10.1111/imb.12534
98. Huang, W.-F., Solter, L., Aronstein, K., & Huang, Z. (2015). Infectivity and virulence of Nosema ceranae and Nosema apis in commercially available North American honey bees. Journal of Invertebrate Pathology, 124, 107–113. https://doi.org/10.1016/j.jip.2014.10.006
99. Huang, W.-F., Solter, L. F., Yau, P. M., & Imai, B. S. (2013). Nosema ceranae Escapes Fumagillin Control in Honey Bees. PLoS Pathogens, 9(3). https://doi.org/10.1371/journal.ppat.1003185
100. Huntsman, E. M., Cho, R. M., Kogan, H. V., McNamara-Bordewick, N. K., Tomko, R. J., & Snow, J. W. (2021). Proteasome inhibition is an effective treatment strategy for microsporidia infection in honey bees. Biomolecules, 11(11). https://doi.org/10.3390/biom11111600
101. Jack, C. J., Uppala, S. S., Lucas, H. M., & Sagili, R. R. (2016). Effects of pollen dilution on infection of Nosema ceranae in honey bees. Journal of Insect Physiology, 87, 12–19. https://doi.org/10.1016/j.jinsphys.2016.01.004
102. Jousse, C., Dalle, C., Abila, A., Traikia, M., Diogon, M., Lyan, B., El Alaoui, H., Vidau, C., & Delbac, F. (2020). A combined LC-MS and NMR approach to reveal metabolic changes in the hemolymph of honeybees infected by the gut parasite Nosema ceranae. Journal of Invertebrate Pathology, 176. https://doi.org/10.1016/j.jip.2020.107478
103. Jovanovic, N. M., Glavinic, U., Ristanic, M., Vejnovic, B., Ilic, T., Stevanovic, J., & Stanimirovic, Z. (2023). Effects of Plant-Based Supplement on Oxidative Stress of Honey Bees (Apis mellifera) Infected with Nosema ceranae. Animals, 13(22). https://doi.org/10.3390/ani13223543
104. Kim, D.-J., Woo, R.-M., Kim, K.-S., & Woo, S.-D. (2023). Screening of Entomopathogenic Fungal Culture Extracts with Honeybee Nosemosis Inhibitory Activity. Insects, 14(6). https://doi.org/10.3390/insects14060538
105. Kim, I.-H., Kim, D.-J., Gwak, W.-S., & Woo, S.-D. (2020). Increased survival of the honey bee Apis mellifera infected with the microsporidian Nosema ceranae by

effective gene silencing. Archives of Insect Biochemistry and Physiology, 105(4). https://doi.org/10.1002/arch.21734
106. Kim, J. H., Park, J. K., & Lee, J. K. (2016). Evaluation of antimicrosporidian activity of plant extracts on Nosema ceranae. Journal of Apicultural Science, 60(2), 167–178. https://doi.org/10.1515/JAS-2016-0027
107. Kurze, C., Dosselli, R., Grassl, J., Le Conte, Y., Kryger, P., Baer, B., & Moritz, R. F. A. (2016). Differential proteomics reveals novel insights into Nosema–honey bee interactions. Insect Biochemistry and Molecular Biology, 79, 42–49. https://doi.org/10.1016/j.ibmb.2016.10.005
108. Kurze, C., Le Conte, Y., Dussaubat, C., Erler, S., Kryger, P., Lewkowski, O., Müller, T., Widder, M., & Moritz, R. F. A. (2015). Nosema tolerant honeybees (apis mellifera) escape parasitic manipulation of apoptosis. PLoS ONE, 10(10). https://doi.org/10.1371/journal.pone.0140174
109. Kurze, C., Le Conte, Y., Kryger, P., Lewkowski, O., Müller, T., & Moritz, R. F. A. (2018). Infection dynamics of Nosema ceranae in honey bee midgut and host cell apoptosis. Journal of Invertebrate Pathology, 154, 1–4. https://doi.org/10.1016/j.jip.2018.03.008
110. Kurze, C., Mayack, C., Hirche, F., Stangl, G. I., Le Conte, Y., Kryger, P., & Moritz, R. F. A. (2016). Nosema spp. Infections cause no energetic stress in tolerant honeybees. Parasitology Research, 115(6), 2381–2388. https://doi.org/10.1007/s00436-016-4988-3
111. Lang, H., Wang, H., Wang, H., Zhong, Z., Xie, X., Zhang, W., Guo, J., Meng, L., Hu, X., Zhang, X., & Zheng, H. (2023). Engineered symbiotic bacteria interfering Nosema redox system inhibit microsporidia parasitism in honeybees. Nature Communications, 14(1). https://doi.org/10.1038/s41467-023-38498-2
112. Lee, J. K., Kim, J. H., Jo, M., Rangachari, B., & Park, J. K. (2018). Anti-nosemosis activity of Aster Scaber and Artemisia Dubia aqueous extracts. Journal of Apicultural Science, 62(1), 27–38. https://doi.org/10.2478/JAS-2018-0003
113. Li, J. H., Evans, J. D., Li, W. F., Zhao, Y. Z., DeGrandi-Hoffman, G., Huang, S. K., Li, Z. G., Hamilton, M., & Chen, Y. P. (2019). New evidence showing that the destruction of gut bacteria by antibiotic treatment could increase the honey bee's vulnerability to nosema infection. PLoS ONE, 12(11). https://doi.org/10.1371/journal.pone.0187505
114. Li, W., Chen, Y., & Cook, S. C. (2018). Chronic Nosema ceranae infection inflicts comprehensive and persistent immunosuppression and accelerated lipid loss in host Apis mellifera honey bees. International Journal for Parasitology, 48(6), 433–444. https://doi.org/10.1016/j.ijpara.2017.11.004
115. Li, W., Evans, J. D., Huang, Q., Rodríguez-García, C., Liu, J., Hamilton, M., Grozinger, C. M., Webster, T. C., Su, S., & Chen, Y. P. (2016). Silencing the honey bee (Apis mellifera) naked cuticle gene (nkd) improves host immune function and reduces Nosema ceranae infections. Applied and Environmental Microbiology, 82(22), 6779–6787. https://doi.org/10.1128/AEM.02105-16
116. Li, Y.-H., Chang, Z.-T., Yen, M.-R., Huang, Y.-F., Chen, T.-H., Chang, J.-C., Wu, M.-C., Yang, Y.-L., Chen, Y.-W., & Nai, Y.-S. (2022). Transcriptome of Nosema ceranae and Upregulated Microsporidia Genes during Its Infection of Western Honey Bee (Apis mellifera). Insects, 13(8). https://doi.org/10.3390/insects13080716

129. Mayack, C., & Naug, D. (2010). Parasitic infection leads to decline in hemolymph sugar levels in honeybee foragers. Journal of Insect Physiology, 56(11), 1572–1575. https://doi.org/10.1016/j.jinsphys.2010.05.016
130. McDonnell, C. M., Alaux, C., Parrinello, H., Desvignes, J.-P., Crauser, D., Durbesson, E., Beslay, D., & Le Conte, Y. (2013). Ecto- and endoparasite induce similar chemical and brain neurogenomic responses in the honey bee (Apis mellifera). BMC Ecology, 13. https://doi.org/10.1186/1472-6785-13-25
131. McGowan, J., De la Mora, A., Goodwin, P. H., Habash, M., Hamiduzzaman, M. M., Kelly, P. G., & Guzman-Novoa, E. (2016). Viability and infectivity of fresh and cryopreserved Nosema ceranae spores. Journal of Microbiological Methods, 131, 16–22. https://doi.org/10.1016/j.mimet.2016.09.021
132. Milbrath, M. O., van Tran, T., Huang, W.-F., Solter, L. F., Tarpy, D. R., Lawrence, F., & Huang, Z. Y. (2015). Comparative virulence and competition between Nosema apis and Nosema ceranae in honey bees (Apis mellifera). Journal of Invertebrate Pathology, 125, 9–15. https://doi.org/10.1016/j.jip.2014.12.006
133. Milbrath, M. O., Xie, X., & Huang, Z. Y. (2013). Nosema ceranae induced mortality in honey bees (Apis mellifera) depends on infection methods. Journal of Invertebrate Pathology, 114(1), 42–44. https://doi.org/10.1016/j.jip.2013.05.006
134. Mura, A., Pusceddu, M., Theodorou, P., Angioni, A., Floris, I., Paxton, R. J., & Satta, A. (2020). Propolis consumption reduces Nosema ceranae infection of European honey bees (Apis mellifera). Insects, 11(2). https://doi.org/10.3390/insects11020124
135. Murray, Z. L., Keyzers, R. A., Barbieri, R. F., Digby, A. P., & Lester, P. J. (2016). Two pathogens change cuticular hydrocarbon profiles but neither elicit a social behavioural change in infected honey bees, Apis mellifera (Apidae: Hymenoptera). Austral Entomology, 55(2), 147–153. https://doi.org/10.1111/aen.12165
136. Nanetti, A., Rodriguez-García, C., Meana, A., Martín-Hernández, R., & Higes, M. (2015). Effect of oxalic acid on Nosema ceranae infection. Research in Veterinary Science, 102, 167–172. https://doi.org/10.1016/j.rvsc.2015.08.003
137. Nanetti, A., Ugolini, L., Cilia, G., Pagnotta, E., Malaguti, L., Cardaio, I., Matteo, R., & Lazzeri, L. (2021). Seed meals from brassica nigra and eruca sativa control artificial nosema ceranae infections in apis mellifera. Microorganisms, 9(5). https://doi.org/10.3390/microorganisms9050949
138. Naree, S., Benbow, M. E., Suwannapong, G., & Ellis, J. D. (2021). Mitigating Nosema ceranae infection in western honey bee (Apis mellifera) workers using propolis collected from honey bee and stingless bee (Tetrigona apicalis) hives. Journal of Invertebrate Pathology, 185. https://doi.org/10.1016/j.jip.2021.107666
139. Naree, S., Ellis, J. D., Benbow, M. E., & Suwannapong, G. (2021). The use of propolis for preventing and treating Nosema ceranae infection in western honey bee (Apis mellifera Linnaeus, 1787) workers. Journal of Apicultural Research, 60(5), 686–696. https://doi.org/10.1080/00218839.2021.1905374
140. Naree, S., Ellis, J. D., Benbow, M. E., & Suwannapong, G. (2022). Experimental Nosema ceranae infection is associated with microbiome changes in the midguts of four species of Apis (honey bees). Journal of Apicultural Research, 61(3), 435–447. https://doi.org/10.1080/00218839.2021.1987086
141. Natsopoulou, M. E., Doublet, V., & Paxton, R. J. (2016). European isolates of the Microsporidia Nosema apis and Nosema ceranae have similar virulence in

laboratory tests on European worker honey bees. Apidologie, 47(1), 57–65. https://doi.org/10.1007/s13592-015-0375-9
142. Natsopoulou, M. E., McMahon, D. P., Doublet, V., Bryden, J., & Paxton, R. J. (2015). Interspecific competition in honeybee intracellular gut parasites is asymmetric and favours the spread of an emerging infectious disease. Proceedings of the Royal Society B: Biological Sciences, 282(1798). https://doi.org/10.1098/rspb.2014.1896
143. Naug, D., & Gibbs, A. (2009). Behavioral changes mediated by hunger in honeybees infected with Nosema ceranae. Apidologie, 40(6), 595–599. https://doi.org/10.1051/apido/2009039
144. Özgör, E. (2021). The effects of nosema apis and nosema ceranae infection on survival and phenoloxidase gene expression in galleria mellonella (Lepidoptera: Galleriidae) compared to apis mellifera. Insects, 12(10). https://doi.org/10.3390/insects12100953
145. Özgör, E., & Keskin, N. (2017). Determination of spore longevity and viability of Nosema apis and Nosema ceranae according to storage conditions. EUROBIOTECH JOURNAL, 1(3), 217–221. https://doi.org/10.24190/ISSN2564-615X/2017/03.03
146. Özklrlm, A., & Küçüközmen, B. (2021). Application of Herbal Essential Oil Extract Mixture for Honey Bees (apis mellifera L.) against Nosema ceranae and Nosema apis. Journal of Apicultural Science, 65(1), 163–175. https://doi.org/10.2478/jas-2021-0010
147. Özüiçli, M., Aydin, L., Girişgin, A. O., Selova, S., & Sabanci, A. Ü. (2023). Determination of the efficacy of thymol, Artemisia absinthium oil and nanoparticle ozone in the treatment of Nosema ceranae in adult honey bees. Journal of Apicultural Research. https://doi.org/10.1080/00218839.2023.2175964
148. Paldi, N., Glick, E., Oliva, M., Zilberberg, Y., Aubin, L., Pettis, J., Chen, Y., & Evans, J. D. (2010). Effective gene silencing in a microsporidian parasite associated with honeybee (Apis mellifera) colony declines. Applied and Environmental Microbiology, 76(17), 5960–5964. https://doi.org/10.1128/AEM.01067-10
149. Palmer-Young, E. C., Tozkar, C. O., Schwarz, R. S., Chen, Y., Irwin, R. E., Adler, L. S., & Evans, J. D. (2017). Nectar and Pollen Phytochemicals Stimulate Honey Bee (Hymenoptera: Apidae) Immunity to Viral Infection. Journal of Economic Entomology, 110(5), 1959–1972. https://doi.org/10.1093/jee/tox193
150. Panek, J., Paris, L., Roriz, D., Mone, A., Dubuffet, A., Delbac, F., Diogon, M., & El Alaoui, H. (2018). Impact of the microsporidian Nosema ceranae on the gut epithelium renewal of the honeybee, Apis mellifera. Journal of Invertebrate Pathology, 159, 121–128. https://doi.org/10.1016/j.jip.2018.09.007
151. Panjad, P., Yongsawas, R., Sinpoo, C., Pakwan, C., Subta, P., Krongdang, S., In-On, A., Chomdej, S., Chantawannakul, P., & Disayathanoowat, T. (2021). Impact of Nosema disease and american foulbrood on gut bacterial communities of honeybees Apis mellifera. Insects, 12(6). https://doi.org/10.3390/insects12060525
152. Paris, L., El Alaoui, H., Delbac, F., & Diogon, M. (2018). Effects of the gut parasite Nosema ceranae on honey bee physiology and behavior. Current Opinion in Insect Science, 26, 149–154. https://doi.org/10.1016/j.cois.2018.02.017
153. Paris, L., Peghaire, E., Moné, A., Diogon, M., Debroas, D., Delbac, F., & El Alaoui, H. (2020). Honeybee gut microbiota dysbiosis in pesticide/parasite co-

exposures is mainly induced by Nosema ceranae. Journal of Invertebrate Pathology, 172. https://doi.org/10.1016/j.jip.2020.107348
154. Paris, L., Roussel, M., Pereira, B., Delbac, F., & Diogon, M. (2017). Disruption of oxidative balance in the gut of the western honeybee Apis mellifera exposed to the intracellular parasite Nosema ceranae and to the insecticide fipronil. Microbial Biotechnology, 10(6), 1702–1717. https://doi.org/10.1111/1751-7915.12772
155. Parrella, P., Elikan, A. B., Kogan, H. V., Wague, F., Marshalleck, C. A., & Snow, J. W. (2024). Bleomycin reduces Vairimorpha (Nosema) ceranae infection in honey bees with some evident host toxicity. MICROBIOLOGY SPECTRUM. https://doi.org/10.1128/spectrum.03349-23
156. Pașca, C., Matei, I. A., Diaconeasa, Z., Rotaru, A., Erler, S., & Dezmirean, D. S. (2021). Biologically active extracts from different medicinal plants tested as potential additives against bee pathogens. Antibiotics, 10(8). https://doi.org/10.3390/antibiotics10080960
157. Paxton, R. J., Klee, J., Korpela, S., & Fries, I. (2007). Nosema ceranae has infected Apis mellifera in Europe since at least 1998 and may be more virulent than Nosema apis. Apidologie, 38(6), 558–565. https://doi.org/10.1051/apido:2007037
158. Peghaire, E., Moné, A., Delbac, F., Debroas, D., Chaucheyras-Durand, F., & El Alaoui, H. (2020). A Pediococcus strain to rescue honeybees by decreasing Nosema ceranae- and pesticide-induced adverse effects. Pesticide Biochemistry and Physiology, 163, 138–146. https://doi.org/10.1016/j.pestbp.2019.11.006
159. Pent, K., Naudi, S., Raimets, R., Jürison, M., Liiskmann, E., & Karise, R. (2023). Overlapping exposure effects of pathogen and dimethoate on honeybee (Apis mellifera Linnaeus) metabolic rate and longevity. Frontiers in Physiology, 14. https://doi.org/10.3389/fphys.2023.1198070
160. Pérez-Morfi, A., Canto, A., Feldman, R. E., Medina-Medina, L. A., Estrella-Maldonado, H., Rodríguez, R., & Andrade, J. L. (2023). Effect of bee bread on Africanized honey bees infected with spores of Nosema spp. Entomologia Experimentalis et Applicata, 171(5), 374–385. https://doi.org/10.1111/eea.13286
161. Pettis, J. S., Lichtenberg, E. M., Andree, M., Stitzinger, J., Rose, R., & vanEngelsdorp, D. (2013). Crop Pollination Exposes Honey Bees to Pesticides Which Alters Their Susceptibility to the Gut Pathogen Nosema ceranae. PLoS ONE, 8(7). https://doi.org/10.1371/journal.pone.0070182
162. Piiroinen, S., & Goulson, D. (2016). Chronic neonicotinoid pesticide exposure and parasite stress differentially affects learning in honeybees and bumblebees. Proceedings of the Royal Society B: Biological Sciences, 283(1828). https://doi.org/10.1098/rspb.2016.0246
163. Porrini, L. P., Porrini, M. P., Garrido, M. P., Müller, F., Arrascaeta, L., Fernández Iriarte, P. J., & Eguaras, M. J. (2020). Infectivity and virulence of Nosema ceranae (Microsporidia) isolates obtained from various Apis mellifera morphotypes. Entomologia Experimentalis et Applicata, 168(4), 286–294. https://doi.org/10.1111/eea.12902
164. Porrini, M. P., Audisio, M. C., Sabaté, D. C., Ibarguren, C., Medici, S. K., Sarlo, E. G., Garrido, P. M., & Eguaras, M. J. (2010). Effect of bacterial metabolites on microsporidian Nosema ceranae and on its host Apis mellifera. Parasitology Research, 107(2), 381–388. https://doi.org/10.1007/s00436-010-1875-1

177. Rodríguez-García, C., Evans, J. D., Li, W., Branchiccela, B., Li, J. H., Heerman, M. C., Banmeke, O., Zhao, Y., Hamilton, M., Higes, M., Martín-Hernández,

R., & Chen, Y. P. (2018). Nosemosis control in European honey bees, Apis mellifera, by silencing the gene encoding Nosema ceranae polar tube protein 3. Journal of Experimental Biology, 221(19). https://doi.org/10.1242/jeb.184606
178. Rodríguez-García, C., Heerman, M. C., Cook, S. C., Evans, J. D., DeGrandi-Hoffman, G., Banmeke, O., Zhang, Y., Huang, S., Hamilton, M., & Chen, Y. P. (2021). Transferrin-mediated iron sequestration suggests a novel therapeutic strategy for controlling Nosema disease in the honey bee, Apis mellifera. PLoS Pathogens, 17(2). https://doi.org/10.1371/JOURNAL.PPAT.1009270
179. Roussel, M., Villay, A., Delbac, F., Michaud, P., Laroche, C., Roriz, D., El Alaoui, H., & Diogon, M. (2015). Antimicrosporidian activity of sulphated polysaccharides from algae and their potential to control honeybee nosemosis. Carbohydrate Polymers, 133, 213–220. https://doi.org/10.1016/j.carbpol.2015.07.022
180. Schwarz, R. S., & Evans, J. D. (2013). Single and mixed-species trypanosome and microsporidia infections elicit distinct, ephemeral cellular and humoral immune responses in honey bees. Developmental and Comparative Immunology, 40(3–4), 300–310. https://doi.org/10.1016/j.dci.2013.03.010
181. Simone-Finstrom, M., Aronstein, K., Goblirsch, M., Rinkevich, F., & de Guzman, L. (2018). Gamma irradiation inactivates honey bee fungal, microsporidian, and viral pathogens and parasites. Journal of Invertebrate Pathology, 153, 57–64. https://doi.org/10.1016/j.jip.2018.02.011
182. Sinpoo, C., Paxton, R. J., Disayathanoowat, T., Krongdang, S., & Chantawannakul, P. (2018). Impact of Nosema ceranae and Nosema apis on individual worker bees of the two host species (Apis cerana and Apis mellifera) and regulation of host immune response. Journal of Insect Physiology, 105, 1–8. https://doi.org/10.1016/j.jinsphys.2017.12.010
183. Smith, M. L. (2012). The honey bee parasite nosema ceranae: Transmissible via food exchange? PLoS ONE, 7(8). https://doi.org/10.1371/journal.pone.0043319
184. Snow, J. W. (2020). Prolyl-tRNA synthetase inhibition reduces microsporidia infection intensity in honey bees. Apidologie, 51(4), 557–569. https://doi.org/10.1007/s13592-020-00742-9
185. Straub, L., Minnameyer, A., Strobl, V., Kolari, E., Friedli, A., Kalbermatten, I., Merkelbach, A. J. W. M., Victor Yañez, O., & Neumann, P. (2020). From antagonism to synergism: Extreme differences in stressor interactions in one species. Scientific Reports, 10(1). https://doi.org/10.1038/s41598-020-61371-x
186. Sulborska, A., Horecka, B., Cebrat, M., Kowalczyk, M., Skrzypek, T. H., Kazimierczak, W., Trytek, M., & Borsuk, G. (2019). Microsporidia Nosema spp. – Obligate bee parasites are transmitted by air. Scientific Reports, 9(1). https://doi.org/10.1038/s41598-019-50974-8
187. Tadei, R., Menezes-Oliveira, V. B., & Silva-Zacarin, E. C. M. (2020). Silent effect of the fungicide pyraclostrobin on the larval exposure of the non-target organism Africanized Apis mellifera and its interaction with the pathogen Nosema ceranae in adulthood. Environmental Pollution, 267. https://doi.org/10.1016/j.envpol.2020.115622
188. Tesovnik, T., Zorc, M., Ristanić, M., Glavinić, U., Stevanović, J., Narat, M., & Stanimirović, Z. (2020). Exposure of honey bee larvae to thiamethoxam and its interaction with Nosema ceranae infection in adult honey bees. Environmental Pollution, 256. https://doi.org/10.1016/j.envpol.2019.113443

Excluded

208. Abbas, M. N., Kausar, S., Gul, I., Li, J., Yu, H., Dong, M., & Cui, H. (2023). The Potential Biological Roles of Circular RNAs in the Immune Systems of Insects to Pathogen Invasion. Genes, 14(4). https://doi.org/10.3390/genes14040895
209. Abd-El-Samie, E. M., Basuny, N. K., & Seyam, H. (2021). Molecular characterization of viruses found in honeybee (Apis mellifera) colonies infested with Varroa destructor and Nosema cerana in Egypt. Molecular and Cellular Probes, 57. https://doi.org/10.1016/j.mcp.2021.101731
210. Abdi, K., Belguith, K., Hamdi, C., Souissi, Y., Essanaa, J., Dridi, W., Hajji, T., Mosbah, A., Ben Hamida, T., & Cherif, A. (2018). Parasites-iflavirus association and emergence of three master variants of DWV affecting Apis mellifera intermissa in Tunisian apiaries. Bulletin of Insectology, 71(2), 273–282.

211. Abou Kubaa, R., Molinatto, G., Solaiman Khaled, B., Daher-Hjaij, N., Heinoun, K., & Saponari, M. (2018). First detection of black queen cell virus, Varroa destructor macula-like virus, Apis mellifera filamentous virus and Nosema ceranae in Syrian honey bees Apis mellifera syriaca. Bulletin of Insectology, 71(2), 217–224.
212. Aditya, I. R. A., & Purwanto, H. (2023). Molecular detection of the pathogen of Apis mellifera (Hymenoptera: Apidae) in honey in Indonesia. Biodiversitas, 24(5), 2612–2622. https://doi.org/10.13057/biodiv/d240513
213. Aglagane, A., Carra, E., Ravaioli, V., Er-Rguibi, O., Santo, E., Mouden, E. H. E., Aourir, M., & Frasnelli, M. (2023). Molecular examination of nosemosis and foulbrood pathogens in honey bee populations from southeastern Morocco. Apidologie, 54(4). https://doi.org/10.1007/s13592-023-01022-y
214. Agripina, S., Savu, V., Radoi, I., Tapaloaga, D., Tanase, P., & Calin, V. (2017). Evaluation of results in research made in order to obtain a phytotherapeutic product for the prophylaxis and fight against nosema in bees. EUROBIOTECH JOURNAL, 1(1), 36–40. https://doi.org/10.24190/ISSN2564-615X/2017/01.06
215. Aguado-López, D., Bartolomé, C., Lopes, A. R., Henriques, D., Segura, S. K., Maside, X., Pinto, M. A., Higes, M., & Martín-Hernández, R. (2023). Frequent Parasitism of Apis mellifera by Trypanosomatids in Geographically Isolated Areas with Restricted Beekeeping Movements. Microbial Ecology, 86(4), 2655–2665. https://doi.org/10.1007/s00248-023-02266-y
216. Akkaya, H., Bayrakal, G. M., & Dümen, E. (2020). Investigation of propolis in terms of hygienic quality, some pathogenic bacteria and Nosema spp. Turkish Journal of Veterinary and Animal Sciences, 44(4), 838–844. https://doi.org/10.3906/vet-2001-20
217. Alaux, C., Crauser, D., Pioz, M., Saulnier, C., & Le Conte, Y. (2014). Parasitic and immune modulation of flight activity in honey bees tracked with optical counters. Journal of Experimental Biology, 217(19), 3416–3424. https://doi.org/10.1242/jeb.105783
218. Alaux, C., Folschweiller, M., McDonnell, C., Beslay, D., Cousin, M., Dussaubat, C., Brunet, J.-L., & Conte, Y. L. (2011). Pathological effects of the microsporidium Nosema ceranae on honey bee queen physiology (Apis mellifera). Journal of Invertebrate Pathology, 106(3), 380–385. https://doi.org/10.1016/j.jip.2010.12.005
219. Al-Hameed, A. S. A., & Hadi, H. A. A.-A. (2020). Evaluation of the first report of (Nosema ceranae) disease on honey bees in Iraq. Plant Archives, 20, 3027–3030.
220. Alonso-Prados, E., González-Porto, A. V., García-Villarubia, C., López-Pérez, J. A., Valverde, S., Bernal, J., Martín-Hernández, R., & Higes, M. (2022). Effects of Thiamethoxam-Dressed Oilseed Rape Seeds and Nosema ceranae on Colonies of Apis mellifera iberiensis, L. under Field Conditions of Central Spain. Is Hormesis Playing a Role? Insects, 13(4). https://doi.org/10.3390/insects13040371
221. Alonso-Prados, E., González-Porto, A.-V., Bernal, J. L., Bernal, J., Martín-Hernández, R., & Higes, M. (2021). A case report of chronic stress in honey bee colonies induced by pathogens and acaricide residues. Pathogens, 10(8). https://doi.org/10.3390/pathogens10080955
222. Alonso-Prados, E., Muñoz, I., De la Rúa, P., Serrano, J., Fernández-Alba, A. R., García-Valcárcel, A. I., Hernando, M. D., Alonso, Á., Alonso-Prados, J. L., Bartolomé, C., Maside, X., Barrios, L., Martín-Hernández, R., & Higes, M. (2020).

The toxic unit approach as a risk indicator in honey bees surveillance programmes: A case of study in Apis mellifera iberiensis. Science of the Total Environment, 698. https://doi.org/10.1016/j.scitotenv.2019.134208
223. Anido, M., Branchiccela, B., Castelli, L., Harriet, J., Campá, J., Zunino, P., & Antúnez, K. (2015). Prevalence and distribution of honey bee pests and pathogens in Uruguay. Journal of Apicultural Research, 54(5), 532–540. https://doi.org/10.1080/00218839.2016.1175731
224. Ansari, M. J., Al-Ghamdi, A., Nuru, A., Khan, K. A., & Alattal, Y. (2017). Geographical distribution and molecular detection of Nosema ceranae from indigenous honey bees of Saudi Arabia. Saudi Journal of Biological Sciences, 24(5), 983–991. https://doi.org/10.1016/j.sjbs.2017.01.054
225. Antúnez, K., Anido, M., Branchiccela, B., Harriet, J., Campa, J., Invernizzi, C., Santos, E., Higes, M., Martín-Hernández, R., & Zunino, P. (2015). Seasonal Variation of Honeybee Pathogens and its Association with Pollen Diversity in Uruguay. Microbial Ecology, 70(2), 522–533. https://doi.org/10.1007/s00248-015-0594-7
226. Antúnez, K., Invernizzi, C., & Zuninoa, P. (2012). Why massive honeybee colony losses do not occur in Uruguay? In Bees: Biology, Threats and Colonies (pp. 189–208). https://www.scopus.com/inward/record.uri?eid=2-s2.0-84871718050&partnerID=40&md5=f4a1de6d3eacf74273a59d62e08bfaf3
227. Applegate, J. R., & Petritz, O. A. (2020). Common and Emerging Infectious Diseases of Honeybees (Apis mellifera). Veterinary Clinics of North America - Exotic Animal Practice, 23(2), 285–297. https://doi.org/10.1016/j.cvex.2020.01.001
228. Araneda, X., Cumian, M., & Morales, D. (2015). Distribution, epidemiological characteristics and control methods of the pathogen Nosema ceranae Fries in honey bees Apis mellifera L. (Hymenoptera, Apidae). Archivos de Medicina Veterinaria, 47(2), 129–138. https://doi.org/10.4067/S0301-732X2015000200002
229. Arbulo, N., Antúnez, K., Salvarrey, S., Santos, E., Branchiccela, B., Martín-Hernández, R., Higes, M., & Invernizzi, C. (2015). High prevalence and infection levels of Nosema ceranae in bumblebees Bombus atratus and Bombus bellicosus from Uruguay. Journal of Invertebrate Pathology, 130, 165–168. https://doi.org/10.1016/j.jip.2015.07.018
230. Ares, A. M., Nozal, M. J., Bernal, J. L., Martín-Hernández, R., & Bernal, J. (2012). Liquid chromatography coupled to ion trap-tandem mass spectrometry to evaluate juvenile hormone III levels in bee hemolymph from Nosema spp. Infected colonies. Journal of Chromatography B: Analytical Technologies in the Biomedical and Life Sciences, 899, 146–153. https://doi.org/10.1016/j.jchromb.2012.05.016
231. Aroee, F., Azizi, H., Shiran, B., & Pirali Kheirabadi, K. (2017). Molecular identification of Nosema species in provinces of Fars, Chaharmahal and Bakhtiari and Isfahan (Southwestern Iran). Asian Pacific Journal of Tropical Biomedicine, 7(1), 10–13. https://doi.org/10.1016/j.apjtb.2016.11.004
232. Aronstein, K. A., Saldivar, E., & Webster, T. C. (2011). Evaluation of Nosema ceranae spore-specific polyclonal antibodies. Journal of Apicultural Research, 50(2), 145–151. https://doi.org/10.3896/IBRA.1.50.2.06
233. Aronstein, K. A., Webster, T. C., & Saldivar, E. (2013). A serological method for detection of Nosema ceranae. Journal of Applied Microbiology, 114(3), 621–625. https://doi.org/10.1111/jam.12066

246. Blažytė-Čereškienė, L., Skrodenytė-Arbačiauskienė, V., Radžiutė, S., Nedveckytė, I., & Būda, V. (2016). Honey bee infection caused by Nosema spp. In Lithuania. Journal of Apicultural Science, 60(2), 77–88. https://doi.org/10.1515/JAS-2016-0019
247. Blot, N., Clémencet, J., Jourda, C., Lefeuvre, P., Warrit, N., Esnault, O., & Delatte, H. (2023). Geographic population structure of the honeybee microsporidian parasite Vairimorpha (Nosema) ceranae in the South West Indian Ocean. Scientific Reports, 13(1). https://doi.org/10.1038/s41598-023-38905-0
248. Bollan, K. A., Hothersall, J. D., Moffat, C., Durkacz, J., Saranzewa, N., Wright, G. A., Raine, N. E., Highet, F., & Connolly, C. N. (2013). The microsporidian parasites Nosema ceranae and Nosema apis are widespread in honeybee (Apis mellifera) colonies across Scotland. Parasitology Research, 112(2), 751–759. https://doi.org/10.1007/s00436-012-3195-0
249. Bordier, C., Pioz, M., Crauser, D., Le Conte, Y., & Alaux, C. (2017). Should I stay or should I go: Honeybee drifting behaviour as a function of parasitism. Apidologie, 48(3), 286–297. https://doi.org/10.1007/s13592-016-0475-1
250. Bordin, F., Zulian, L., Granato, A., Caldon, M., Colamonico, R., Toson, M., Trevisan, L., Biasion, L., & Mutinelli, F. (2022). Presence of Known and Emerging Honey Bee Pathogens in Apiaries of Veneto Region (Northeast of Italy) during Spring 2020 and 2021. Applied Sciences (Switzerland), 12(4). https://doi.org/10.3390/app12042134
251. Borneck, R., Viry, A., Martín-Hernández, R., & Higes, M. (2010). Honey bee colony losses in the Jura Region, France and related pathogens. Journal of Apicultural Research, 49(4), 334–336. https://doi.org/10.3896/IBRA.1.49.4.06
252. Borsuk, G., Kozłowska, M., Anusiewicz, M., & Paleolog, J. (2018). Nosema ceranae changes semen characteristics and damages sperm DNA in honeybee drones. Invertebrate Survival Journal, 15, 197–202.
253. Botías, C., Anderson, D. L., Meana, A., Garrido-Bailón, E., Martín-Hernández, R., & Higes, M. (2012). Further evidence of an oriental origin for Nosema ceranae (Microsporidia: Nosematidae). Journal of Invertebrate Pathology, 110(1), 108–113. https://doi.org/10.1016/j.jip.2012.02.014
254. Botías, C., Jones, J. C., Pamminger, T., Bartomeus, I., Hughes, W. O. H., & Goulson, D. (2021). Multiple stressors interact to impair the performance of bumblebee Bombus terrestris colonies. Journal of Animal Ecology, 90(2), 415–431. https://doi.org/10.1111/1365-2656.13375
255. Botías, C., Martín-Hernández, R., Barrios, L., Garrido-Bailón, E., Nanetti, A., Meana, A., & Higes, M. (2012). Nosema spp. Parasitization decreases the effectiveness of acaricide strips (Apivar ®) in treating varroosis of honey bee (Apis mellifera iberiensis) colonies. Environmental Microbiology Reports, 4(1), 57–65. https://doi.org/10.1111/j.1758-2229.2011.00299.x
256. Botías, C., Martín-Hernández, R., Barrios, L., Meana, A., & Higes, M. (2013). Nosema spp. Infection and its negative effects on honey bees (Apis mellifera iberiensis) at the colony level. Veterinary Research, 44(1). https://doi.org/10.1186/1297-9716-44-25
257. Botías, C., Martín-Hernández, R., Días, J., García-Palencia, P., Matabuena, M., Juarranz, A., Barrios, L., Meana, A., Nanetti, A., & Higes, M. (2012). The effect of induced queen replacement on Nosema spp. Infection in honey bee (Apis mellifera

iberiensis) colonies. Environmental Microbiology, 14(4), 845–859. https://doi.org/10.1111/j.1462-2920.2011.02647.x
258. Botías, C., Martín-Hernández, R., Garrido-Bailón, E., González-Porto, A., Martínez-Salvador, A., De La Rúa, P., Meana, A., & Higes, M. (2012). The growing prevalence of Nosema ceranae in honey bees in Spain, an emerging problem for the last decade. Research in Veterinary Science, 93(1), 150–155. https://doi.org/10.1016/j.rvsc.2011.08.002
259. Botías, C., Martín-Hernández, R., Meana, A., & Higes, M. (2012). Critical aspects of the Nosema spp. Diagnostic sampling in honey bee (Apis mellifera L.) colonies. Parasitology Research, 110(6), 2557–2561. https://doi.org/10.1007/s00436-011-2760-2
260. Botías, C., Martín-Hernández, R., Meana, A., & Higes, M. (2013). Screening alternative therapies to control Nosemosis type C in honey bee (Apis mellifera iberiensis) colonies. Research in Veterinary Science, 95(3), 1041–1045. https://doi.org/10.1016/j.rvsc.2013.09.012
261. Bourgeois, A. L., Rinderer, T. E., Beaman, L. D., & Danka, R. G. (2010). Genetic detection and quantification of Nosema apis and N. ceranae in the honey bee. Journal of Invertebrate Pathology, 103(1), 53–58. https://doi.org/10.1016/j.jip.2009.10.009
262. Bourgeois, A. L., Rinderer, T. E., Sylvester, H. A., Holloway, B., & Oldroyd, B. P. (2012). Patterns of Apis mellifera infestation by Nosema ceranae support the parasite hypothesis for the evolution of extreme polyandry in eusocial insects. Apidologie, 43(5), 539–548. https://doi.org/10.1007/s13592-012-0121-5
263. Bourgeois, L., Beaman, L., Holloway, B., & Rinderer, T. E. (2012). External and internal detection of Nosema ceranae on honey bees using real-time PCR. Journal of Invertebrate Pathology, 109(3), 323–325. https://doi.org/10.1016/j.jip.2012.01.002
264. Bradford, E. L., Gregory, C. L., Roman Longoria, A., Jones, K. R., Bueren, E. K., Haak, D. C., Fell, R., & Belden, L. K. (2022). A new duplex qPCR assay for the quantification of honey bee (Apis mellifera) parasites Nosema ceranae and Nosema apis tested with low dose experimental exposure. Journal of Apicultural Research. https://doi.org/10.1080/00218839.2022.2083846
265. Bramke, K., Müller, U., McMahon, D. P., & Rolff, J. (2019). Exposure of larvae of the solitary bee osmia bicornis to the honey bee pathogen nosema ceranae affects life history. Insects, 10(11). https://doi.org/10.3390/insects10110380
266. Branchiccela, B., Arredondo, D., Higes, M., Invernizzi, C., Martín-Hernández, R., Tomasco, I., Zunino, P., & Antúnez, K. (2017). Characterization of Nosema ceranae Genetic Variants from Different Geographic Origins. Microbial Ecology, 73(4), 978–987. https://doi.org/10.1007/s00248-016-0880-z
267. Branchiccela, B., Castelli, L., Díaz-Cetti, S., Invernizzi, C., Mendoza, Y., Santos, E., Silva, C., Zunino, P., & Antúnez, K. (2023). Can pollen supplementation mitigate the impact of nutritional stress on honey bee colonies? Journal of Apicultural Research, 62(2), 294–302. https://doi.org/10.1080/00218839.2021.1888537
268. Bravi, M. E., Alvarez, L. J., Lucia, M., Pecoraro, M. R. I., García, M. L. G., & Reynaldi, F. J. (2019). Wild bumble bees (Hymenoptera: Apidae: Bombini) as a potential reservoir for bee pathogens in northeastern Argentina. Journal of Apicultural Research, 58(5), 710–713. https://doi.org/10.1080/00218839.2019.1655183

281. Calderón, R. A., Sanchez, L. A., Yañez, O., & Fallas, N. (2008b). Presence of Nosema ceranae in Africanized honey bee colonies in Costa Rica. Journal of Apicultural Research, 47(4), 328–329. https://doi.org/10.3896/IBRA.1.47.4.18
282. Carletto, J., Blanchard, P., Gauthier, A., Schurr, F., Chauzat, M.-P., & Ribière, M. (2013). Improving molecular discrimination of Nosema apis and Nosema ceranae. Journal of Invertebrate Pathology, 113(1), 52–55. https://doi.org/10.1016/j.jip.2013.01.005
283. Carroll, M. J., Meikle, W. G., McFrederick, Q. S., Rothman, J. A., Brown, N., Weiss, M., Ruetz, Z., & Chang, E. (2018). Pre-almond supplemental forage improves colony survival and alters queen pheromone signaling in overwintering honey bee colonies. Apidologie, 49(6), 827–837. https://doi.org/10.1007/s13592-018-0607-x
284. Carvalho, S., Roat, T., Pereira, A. M., Silva-Zacarin, E., Nocelli, R. C. F., Carvalho, C., & Malaspina, O. (2012). Losses of Brazilian bees: An overview of factors that may affect these pollinators. In P. A. Oomen & H. Thompson (Eds.), HAZARDS OF PESTICIDES TO BEES: 11TH INTERNATIONAL SYMPOSIUM OF THE ICP-PR BEE PROTECTION GROUP (Vol. 437, Issue 11th International Symposium of the ICP-BR-Bee-Protection-Group on Hazards of Pesticides to bees, pp. 159–166). https://doi.org/10.5073/jka.2012.437.043
285. Cavigli, I., Daughenbaugh, K. F., Martin, M., Lerch, M., Banner, K., Garcia, E., Brutscher, L. M., & Flenniken, M. L. (2016). Pathogen prevalence and abundance in honey bee colonies involved in almond pollination. Apidologie, 47(2), 251–266. https://doi.org/10.1007/s13592-015-0395-5
286. Cepero, A., Martín-Hernández, R., Bartolomé, C., Gómez-Moracho, T., Barrios, L., Bernal, J., Teresa Martín, M., Meana, A., & Higes, M. (2015). Passive laboratory surveillance in Spain: Pathogens as risk factors for honey bee colony collapse. Journal of Apicultural Research, 54(5), 525–531. https://doi.org/10.1080/00218839.2016.1162978
287. Cepero, A., Ravoet, J., Gómez-Moracho, T., Bernal, J. L., Del Nozal, M. J., Bartolomé, C., Maside, X., Meana, A., González-Porto, A. V., De Graaf, D. C., Martín-Hernández, R., & Higes, M. (2014). Holistic screening of collapsing honey bee colonies in Spain: A case study. BMC Research Notes, 7(1). https://doi.org/10.1186/1756-0500-7-649
288. Chabar, M., Tefiel, H., Adidoy-Chabar, N., Doumandji-Mitiche, B., & Gaouar, S. B. S. (2016). First spatial distribution of nosemosis (Nosema sp.) infected local bee, apis mellifera intermissa l. In Algeria. Egyptian Journal of Biological Pest Control, 26(2), 357–363.
289. Chagas, D. B., Monteiro, F. L., Barcelos, L. D. S., Frühauf, M. I., Ribeiro, L. C., de Lima, M., Hübner, S. D. O., & Fischer, G. (2020). Black queen cell virus and Nosema ceranae coinfection in Africanized honey bees from southern Brazil. Pesquisa Veterinaria Brasileira, 40(11), 892–897. https://doi.org/10.1590/1678-5150-PVB-6678
290. Chaimanee, V., Chantawannakul, P., Chen, Y., Evans, J. D., & Pettis, J. S. (2014). Effects of host age on susceptibility to infection and immune gene expression in honey bee queens (Apis mellifera) inoculated with Nosema ceranae. Apidologie, 45(4), 451–463. https://doi.org/10.1007/s13592-013-0258-x
291. Chaimanee, V., Chen, Y., Pettis, J. S., Scott Cornman, R., & Chantawannakul, P. (2011). Phylogenetic analysis of Nosema ceranae isolated from European and Asian

honeybees in Northern Thailand. Journal of Invertebrate Pathology, 107(3), 229–233. https://doi.org/10.1016/j.jip.2011.05.012
292. Chaimanee, V., Warrit, N., & Chantawannakul, P. (2010). Infections of Nosema ceranae in four different honeybee species. Journal of Invertebrate Pathology, 105(2), 207–210. https://doi.org/10.1016/j.jip.2010.06.005
293. Chang, Z.-T., Ko, C.-Y., Yen, M.-R., Chen, Y.-W., & Nai, Y.-S. (2020). Screening of differentially expressed microsporidia genes from nosema ceranae infected honey bees by suppression subtractive hybridization. Insects, 11(3). https://doi.org/10.3390/insects11030199
294. Chantawannakul, P. (2018). Bee diversity and current status of beekeeping in Thailand. In Asian Beekeeping in the 21st Century (pp. 269–285). https://doi.org/10.1007/978-981-10-8222-1_12
295. Chantawannakul, P., de Guzman, L. I., Li, J., & Williams, G. R. (2016). Parasites, pathogens, and pests of honeybees in Asia. Apidologie, 47(3), 301–324. https://doi.org/10.1007/s13592-015-0407-5
296. Chantawannakul, P., Williams, G., & Neumann, P. (2018). Asian beekeeping in the 21st century. In Asian Beekeeping in the 21st Century (p. 325). https://doi.org/10.1007/978-981-10-8222-1
297. Charistos, L., Parashos, N., & Hatjina, F. (2015). Long term effects of a food supplement HiveAlive™ on honey bee colony strength and Nosema ceranae spore counts. Journal of Apicultural Research, 54(5), 420–426. https://doi.org/10.1080/00218839.2016.1189231
298. Chauzat, M.-P., Higes, M., Martín-Hernández, R., Meana, A., Cougoule, N., & Faucon, J.-P. (2007). Presence of nosema ceranae in French honey bee colonies. Journal of Apicultural Research, 46(2), 127–128. https://doi.org/10.1080/00218839.2007.11101380
299. Chávez-Hernández, E., Otero-Colina, G., Llanderal-Cázares, C., Maggi-Daniel, M., Rodríguez-Dehaibes, S. R., Soto-Rojas, L., Rocha-Martínez, M. K., & Pérez-de la Rosa, J. D. (2021). No effect of abscisic and p-coumaric acids as food supplements and stimulants of the immunological system of Africanized hybrids of Apis mellifera. Journal of Apicultural Research. https://doi.org/10.1080/00218839.2021.2013423
300. Chemurot, M., De Smet, L., Brunain, M., De Rycke, R., & de Graaf, D. C. (2017). Nosema neumanni n. Sp. (Microsporidia, Nosematidae), a new microsporidian parasite of honeybees, Apis mellifera in Uganda. European Journal of Protistology, 61, 13–19. https://doi.org/10.1016/j.ejop.2017.07.002
301. Chen, D., Chen, H., Du, Y., Zhou, D., Geng, S., Wang, H., Wan, J., Xiong, C., Zheng, Y., & Guo, R. (2019). Genome-wide identification of long non-coding rnas and their regulatory networks involved in apis mellifera ligustica response to nosema ceranae infection. Insects, 10(8). https://doi.org/10.3390/insects10080245
302. Chen, D., Du, Y., Chen, H., Fan, Y., Fan, X., Zhu, Z., Wang, J., Xiong, C., Zheng, Y., Hou, C., Diao, Q., & Guo, R. (2019). Comparative identification of microRNAs in Apis cerana cerana workers' midguts in response to Nosema ceranae Invasion. Insects, 10(9). https://doi.org/10.3390/insects10090258
303. Chen, H., Fan, Y., Jiang, H., Wang, J., Fan, X., Zhu, Z., Long, Q., Cai, Z., Zheng, Y., Fu, Z., Xu, G., Chen, D., & Guo, R. (2021). Improvement of nosema ceranae genome annotation based on nanopore full-length transcriptome data. Scientia

Agricultura Sinica, 54(6), 1288–1300. https://doi.org/10.3864/j.issn.0578-1752.2021.06.018
304. Chen, H.-Z., Fan, X.-X., Fan, Y.-C., Wang, J., Zhu, Z.-W., Jiang, H.-B., Zhang, W.-D., Long, Q., Xiong, C.-L., Zheng, Y.-Z., Fu, Z.-M., Xu, G.-J., Chen, D.-F., & Guo, R. (2021). Analysis of alternative splicing and alternative polyadenylation of Nosema ceranae genes. Mycosystema, 40(1), 161–173. https://doi.org/10.13346/j.mycosystema.200216
305. Chen, Y., Evans, J. D., Smith, I. B., & Pettis, J. S. (2008). Nosema ceranae is a long-present and wide-spread microsporidian infection of the European honey bee (Apis mellifera) in the United States. Journal of Invertebrate Pathology, 97(2), 186–188. https://doi.org/10.1016/j.jip.2007.07.010
306. Chen, Y., Evans, J. D., Zhou, L., Boncristiani, H., Kimura, K., Xiao, T., Litkowski, A. M., & Pettis, J. S. (2009). Asymmetrical coexistence of Nosema ceranae and Nosema apis in honey bees. Journal of Invertebrate Pathology, 101(3), 204–209. https://doi.org/10.1016/j.jip.2009.05.012
307. Chen, Y. P., Evans, J. D., Murphy, C., Gutell, R., Zuker, M., Gundensen-Rindal, D., & Pettis, J. S. (2009). Morphological, molecular, and phylogenetic characterization of Nosema ceranae, a microsporidian parasite isolated from the European honey bee, Apis mellifera. Journal of Eukaryotic Microbiology, 56(2), 142–147. https://doi.org/10.1111/j.1550-7408.2008.00374.x
308. Chen, Y. P., & Huang, Z. Y. (2010). Nosema ceranae, a newly identified pathogen of Apis mellifera in the USA and Asia. Apidologie, 41(3), 364–374. https://doi.org/10.1051/apido/2010021
309. Chen, Y. P., Pettis, J. S., Zhao, Y., Liu, X., Tallon, L. J., Sadzewicz, L. D., Li, R., Zheng, H., Huang, S., Zhang, X., Hamilton, M. C., Pernal, S. F., Melathopoulos, A. P., Yan, X., & Evans, J. D. (2013). Genome sequencing and comparative genomics of honey bee microsporidia, Nosema apis reveal novel insights into host-parasite interactions. BMC Genomics, 14(1). https://doi.org/10.1186/1471-2164-14-451
310. Chen, Y.-W., Chung, W.-P., Wang, C.-H., Solter, L. F., & Huang, W.-F. (2012). Nosema ceranae infection intensity highly correlates with temperature. Journal of Invertebrate Pathology, 111(3), 264–267. https://doi.org/10.1016/j.jip.2012.08.014
311. Chupia, V., Patchanee, P., Krutmuang, P., & Pikulkaew, S. (2016). Development and evaluation of loop-mediated isothermal amplification for rapid detection of Nosema ceranae in honeybee. Asian Pacific Journal of Tropical Disease, 6(12), 952–956. https://doi.org/10.1016/S2222-1808(16)61163-5
312. Chupia, V., Pikulkaew, S., Krutmuang, P., Mekchay, S., & Patchanee, P. (2016). Molecular epidemiology and geographical distribution of Nosema ceranae in honeybees, Northern Thailand. Asian Pacific Journal of Tropical Disease, 6(1), 27–31. https://doi.org/10.1016/S2222-1808(15)60980-X
313. Chupia, V., Pikulkaew, S., Ptaszynska, A. A., Borsuk, G., Majewsk, B. M., Dong, Y., Thungrabeab, M., Mekchay, S., & Krutmuang, P. (2019). Seasonal effect of Nosema ceranae in honeybee from Northern Thailand estimated by Loop-Mediated Isothermal Amplification (LAMP). Asia Life Sciences, 28(1), 175–184.
314. Cilia, G., Cabbri, R., Maiorana, G., Cardaio, I., Dall'Olio, R., & Nanetti, A. (2018). A novel TaqMan® assay for Nosema ceranae quantification in honey bee, based on the protein coding gene Hsp70. European Journal of Protistology, 63, 44–50. https://doi.org/10.1016/j.ejop.2018.01.007

329. Copley, T. R., & Jabaji, S. H. (2012). Honeybee glands as possible infection reservoirs of Nosema ceranae and Nosema apis in naturally infected forager bees. Journal of Applied Microbiology, 112(1), 15–24. https://doi.org/10.1111/j.1365-2672.2011.05192.x
330. Core, A., Runckel, C., Ivers, J., Quock, C., Siapno, T., DeNault, S., Brown, B., DeRisi, J., Smith, C. D., & Hafernik, J. (2012). A new threat to honey bees, the parasitic phorid fly apocephalus borealis. PLoS ONE, 7(1). https://doi.org/10.1371/journal.pone.0029639
331. Cornman, R. S., Chen, Y. P., Schatz, M. C., Street, C., Zhao, Y., Desany, B., Egholm, M., Hutchison, S., Pettis, J. S., Lipkin, W. I., & Evans, J. D. (2009). Genomic analyses of the microsporidian Nosema ceranae, an emergent pathogen of honey bees. PLoS Pathogens, 5(6). https://doi.org/10.1371/journal.ppat.1000466
332. Costa, C., Tanner, G., Lodesani, M., Maistrello, L., & Neumann, P. (2011). Negative correlation between Nosema ceranae spore loads and deformed wing virus infection levels in adult honey bee workers. Journal of Invertebrate Pathology, 108(3), 224–225. https://doi.org/10.1016/j.jip.2011.08.012
333. Cristina Dias, A., Taís Ferreira, J., Weinstein Teixeira, É., & Pedro Lourenço, A. (2023). Honey bee viruses in solitary bees in South America: Simultaneous detection and prevalence. Journal of Apicultural Research. https://doi.org/10.1080/00218839.2023.2190066
334. Csáki, T., Heltai, M., Markolt, F., Kovács, B., Békési, L., Ladányi, M., Péntek-Zakar, E., Meana, A., Botías, C., Martín-Hernández, R., & Higes, M. (2015). Permanent prevalence of Nosema Ceranae in honey bees (Apis Mellifera) in Hungary. Acta Veterinaria Hungarica, 63(3), 358–369. https://doi.org/10.1556/004.2015.034
335. Currie, R. W., Pernal, S. F., & Guzmán-Novoa, E. (2010). Honey bee colony losses in Canada. Journal of Apicultural Research, 49(1), 104–106. https://doi.org/10.3896/IBRA.1.49.1.18
336. Dainat, B., Evans, J. D., Chen, Y. P., Gauthier, L., & Neumann, P. (2012). Predictive markers of honey bee colony collapse. PLoS ONE, 7(2). https://doi.org/10.1371/journal.pone.0032151
337. Dainat, B., Evans, J. D., Chen, Y. P., Gauthier, L., & Neumanna, P. (2012). Dead or alive: Deformed wing virus and varroa destructor reduce the life span of winter honeybees. Applied and Environmental Microbiology, 78(4), 981–987. https://doi.org/10.1128/AEM.06537-11
338. D'Alvise, P., Böhme, F., Codrea, M. C., Seitz, A., Nahnsen, S., Binzer, M., Rosenkranz, P., & Hasselmann, M. (2018). The impact of winter feed type on intestinal microbiota and parasites in honey bees. Apidologie, 49(2), 252–264. https://doi.org/10.1007/s13592-017-0551-1
339. D'Alvise, P., Seeburger, V., Gihring, K., Kieboom, M., & Hasselmann, M. (2019). Seasonal dynamics and co-occurrence patterns of honey bee pathogens revealed by high-throughput RT-qPCR analysis. Ecology and Evolution, 9(18), 10241–10252. https://doi.org/10.1002/ece3.5544
340. Daughenbaugh, K. F., Martin, M., Brutscher, L. M., Cavigli, I., Garcia, E., Lavin, M., & Flenniken, M. L. (2015). Honey bee infecting Lake Sinai viruses. Viruses, 7(6), 3285–3309. https://doi.org/10.3390/v7062772
341. Desai, S. D., & Currie, R. W. (2015). Genetic diversity within honey bee colonies affects pathogen load and relative virus levels in honey bees, Apis mellifera

L. Behavioral Ecology and Sociobiology, 69(9), 1527–1541. https://doi.org/10.1007/s00265-015-1965-2
342. Deutsch, K. R., Graham, J. R., Boncristiani, H. F., Bustamante, T., Mortensen, A. N., Schmehl, D. R., Wedde, A. E., Lopez, D. L., Evans, J. D., & Ellis, J. D. (2023). Widespread distribution of honey bee-associated pathogens in native bees and wasps: Trends in pathogen prevalence and co-occurrence. Journal of Invertebrate Pathology, 200. https://doi.org/10.1016/j.jip.2023.107973
343. Diao, Q., Yang, D., Zhao, H., Deng, S., Wang, X., Hou, C., & Wilfert, L. (2019). Prevalence and population genetics of the emerging honey bee pathogen DWV in Chinese apiculture. Scientific Reports, 9(1). https://doi.org/10.1038/s41598-019-48618-y
344. Dolgikh, V. V., Senderskiy, I. V., Zhuravlyov, V. S., Ignatieva, A. N., Timofeev, S. A., Ismatullaeva, D. A., & Mirzakhodjaev, B. A. (2022). Molecular detection of microsporidia Vairimorpha ceranae and Nosema bombycis growth in the lepidopteran Sf9 cell line. Protistology, 16(1), 21–29. https://doi.org/10.21685/1680-0826-2022-16-1-3
345. Dolgikh, V. V., Timofeev, S. A., Zhuravlyov, V. S., & Senderskiy, I. V. (2020). Construction and heterologous overexpression of two chimeric proteins carrying outer hydrophilic loops of Vairimorpha ceranae and Nosema bombycis ATP/ADP carriers. Journal of Invertebrate Pathology, 171. https://doi.org/10.1016/j.jip.2020.107337
346. Dolgikh, V. V., Zhuravlyov, V. S., Senderskiy, I. V., Ignatieva, A. N., Timofeev, S. A., & Seliverstova, E. V. (2022). Heterologous expression of scFv fragment against Vairimorpha (Nosema) ceranae hexokinase in Sf9 cell culture inhibits microsporidia intracellular growth. Journal of Invertebrate Pathology, 191. https://doi.org/10.1016/j.jip.2022.107755
347. Domatskaya, T. F., Domatsky, A. N., & Zinatullina, Z. Y. (2019). Spread of infestations and infections of honey bees on apiaries of Tyumen region and other regions of Russia. UKRAINIAN JOURNAL OF ECOLOGY, 9(2), 1–4.
348. Dos Santos, L. G., Alves, M. L. T. M. F., Message, D., Pinto, F. A., Silva, M. V. G. B., & Teixeira, E. W. (2014). Honey bee health in apiaries in the Vale do Paraíba, São Paulo state, Southeastern Brazil. Sociobiology, 61(3), 307–312. https://doi.org/10.13102/sociobiology.v61i3.307-312
349. Doublet, V., Poeschl, Y., Gogol-Döring, A., Alaux, C., Annoscia, D., Aurori, C., Barribeau, S. M., Bedoya-Reina, O. C., Brown, M. J. F., Bull, J. C., Flenniken, M. L., Galbraith, D. A., Genersch, E., Gisder, S., Grosse, I., Holt, H. L., Hultmark, D., Lattorff, H. M. G., Le Conte, Y., … Grozinger, C. M. (2017). Unity in defence: Honeybee workers exhibit conserved molecular responses to diverse pathogens. BMC Genomics, 18(1). https://doi.org/10.1186/s12864-017-3597-6
350. Du, Y., Fan, X., Jiang, H., Wang, J., Feng, R., Zhang, W., Yu, K., Long, Q., Cai, Z., Xiong, C., Zheng, Y., Chen, D., Fu, Z., Xu, G., & Guo, R. (2021). MicroRNA-mediated cross-Kingdom regulation of apis mellifera ligustica worker to nosema ceranae. Scientia Agricultura Sinica, 54(8), 1805–1820. https://doi.org/10.3864/j.issn.0578-1752.2021.08.019
351. Du, Y., Zhou, D., Chen, H., Xiong, C., Zheng, Y., Chen, D., & Guo, R. (2019). MicroRNA dataset of normal and Nosema ceranae-infected midguts of Apis cerana cerana workers. Data in Brief, 26. https://doi.org/10.1016/j.dib.2019.104518

387. Fu, Z. M., Chen, H. Z., Liu, S. Y., Zhu, Z. W., Fan, X. X., Fan, Y. C., Wan, J. Q., Zhang, L., Xiong, C. L., Xu, G. J., Chen, D. F., & Guo, R. (2019). Immune responses of apis mellifera ligustia to nosema ceranae stress. Scientia Agricultura Sinica, 52(17), 3069–3082. https://doi.org/10.3864/j.issn.0578-1752.2019.17.014
388. Fürst, M. A., McMahon, D. P., Osborne, J. L., Paxton, R. J., & Brown, M. J. F. (2014). Disease associations between honeybees and bumblebees as a threat to wild pollinators. Nature, 506(7488), 364–366. https://doi.org/10.1038/nature12977
389. Gajger, I. T. (2011). Nozevit aerosol application for Nosema ceranae disease treatment. American Bee Journal, 151(11), 1087–1090.
390. Gajger, I. T., Ribaric, J., Matak, M., Svecnjak, L., Kozaric, Z., Nejedli, S., & Skerl, I. M. S. (2015). Zeolite clinoptilolite as a dietary supplement and remedy for honeybee (Apis mellifera L.) colonies. Veterinarni Medicina, 60(12), 696–705. https://doi.org/10.17221/8584-VETMED
391. Gajger, I. T., Tomljanović, Z., & Stanisavljević, L. J. (2013). An environmentally friendly appro ach to the control of varroa destructor mite and nosema ceranae disease in carniolan honeybee (Apis mellifera carnica) colonies. Archives of Biological Sciences, 65(4), 1585–1592. https://doi.org/10.2298/ABS1304585G
392. Gajger, I. T., Vugrek, O., Grilec, D., & Petrinec, Z. (2010). Prevalence and distribution of Nosema ceranae in Croatian honeybee colonies. Veterinarni Medicina, 55(9), 457–462. https://doi.org/10.17221/2983-VETMED
393. Gajger, I. T., Vugrek, O., Petrinec, Z., Grilec, D., & Tomljanović, Z. (2010). Detection of Nosema ceranae in honey bees from Croatia. Journal of Apicultural Research, 49(4), 340–341. https://doi.org/10.3896/IBRA.1.49.4.08
394. Gajger, I. T., Vugrek, O., Pinter, L., & Petrinec, Z. (2009). 'Nozevit patties' treatment of honey bees (Apis mellifera) for the control of Nosema ceranae disease. American Bee Journal, 149(11), 1053–1056.
395. Galajda, R., Valenčáková, A., Sučik, M., & Kandráčová, P. (2021). Nosema disease of european honey bees. Journal of Fungi, 7(9). https://doi.org/10.3390/jof7090714
396. Gamboa, V., Ravoet, J., Brunain, M., Smagghe, G., Meeus, I., Figueroa, J., Riaño, D., & De Graaf, D. C. (2015). Bee pathogens found in Bombus atratus from Colombia: A case study. Journal of Invertebrate Pathology, 129, 36–39. https://doi.org/10.1016/j.jip.2015.05.013
397. Gancarz, M., Hurd, P. J., Latoch, P., Polaszek, A., Michalska-Madej, J., Strapagiel, D., Gnat, S., Załuski, D., Rusinek, R., Starosta, A. L., Krutmuang, P., Hernández, R. M., Pascual, M. H., & Ptaszyńska, A. A. (2021). Dataset of the next-generation sequencing of variable 16S rRNA from bacteria and ITS2 regions from fungi and plants derived from honeybees kept under anthropogenic landscapes. Data in Brief, 36. https://doi.org/10.1016/j.dib.2021.107019
398. García-Vicente, E. J., Martín, M., Rey-Casero, I., Pérez, A., Martínez, R., Bravo, M., Alonso, J. M., & Risco, D. (2023). Effect of feed supplementation with probiotics and postbiotics on strength and health status of honey bee (Apis mellifera) hives during late spring. Research in Veterinary Science, 159, 237–243. https://doi.org/10.1016/j.rvsc.2023.05.001
399. Geng, S., Shi, C., Fan, X., Wang, J., Zhu, Z., Jiang, H., Fan, Y., Chen, H., Du, Y., Wang, X., Xiong, C., Zheng, Y., Fu, Z., Chen, D., & Guo, R. (2020). The

mechanism underlying MicroRNAs-mediated Nosema ceranae infection to Apis mellifera ligustica worker. Scientia Agricultura Sinica, 53(15), 3187–3204. https://doi.org/10.3864/j.issn.0578-1752.2020.15.018
400. Georgi, I., Asoutis Didaras, N., Nikolaidis, M., Dimitriou, T. G., Charistos, L., Hatjina, F., Amoutzias, G. D., & Mossialos, D. (2022). The Impact of Vairimorpha (Nosema) ceranae Natural Infection on Honey Bee (Apis mellifera) and Bee Bread Microbiota. Applied Sciences (Switzerland), 12(22). https://doi.org/10.3390/app122211476
401. Giacobino, A., Pacini, A., Molineri, A., Bulacio-Cagnolo, N., Merke, J., Orellano, E., Gaggiotti, M., & Signorini, M. (2022). Impact of nutritional and sanitary management on Apis mellifera colony dynamics and pathogen loads. Spanish Journal of Agricultural Research, 20(4). https://doi.org/10.5424/sjar/2022204-19634
402. Giacobino, A., Rivero, R., Molineri, A. I., Cagnolo, N. B., Merke, J., Orellano, E., Salto, C., & Signorini, M. (2016). Fumagillin control of Nosema ceranae (Microsporidia: Nosematidae) infection in honey bee (Hymenoptera: Apidae) colonies in Argentina. Veterinaria Italiana, 52(2), 145–151. https://doi.org/10.12834/VetIt.120.337.6
403. Giersch, T., Berg, T., Galea, F., & Hornitzky, M. (2009). Nosema ceranae infects honey bees (Apis mellifera) and contaminates honey in Australia. Apidologie, 40(2), 117–123. https://doi.org/10.1051/apido/2008065
404. Gisder, S., & Genersch, E. (2013). Molecular differentiation of Nosema apis and Nosema ceranae based on species–specific sequence differences in a protein coding gene. Journal of Invertebrate Pathology, 113(1), 1–6. https://doi.org/10.1016/j.jip.2013.01.004
405. Gisder, S., & Genersch, E. (2015). Identification of candidate agents active against N. ceranae infection in honey bees: Establishment of a medium throughput screening assay based on N. ceranae infected cultured cells. PLoS ONE, 10(2). https://doi.org/10.1371/journal.pone.0117200
406. Gisder, S., Hedtke, K., Möckel, N., Frielitz, M.-C., Linde, A., & Genersch, E. (2010). Five-year cohort study of nosema spp. In Germany: Does climate shape virulence and assertiveness of nosema ceranae? Applied and Environmental Microbiology, 76(9), 3032–3038. https://doi.org/10.1128/AEM.03097-09
407. Gisder, S., Horchler, L., Pieper, F., Schüler, V., Šima, P., & Genersch, E. (2020). Rapid gastrointestinal passage may protect Bombus terrestris from becoming a true host for Nosema ceranae. Applied and Environmental Microbiology, 86(12). https://doi.org/10.1128/AEM.00629-20
408. Gisder, S., Mockel, N., Linde, A., & Genersch, E. (2011). A cell culture model for Nosema ceranae and Nosema apis allows new insights into the life cycle of these important honey bee-pathogenic microsporidia. Environmental Microbiology, 13(2), 404–413. https://doi.org/10.1111/j.1462-2920.2010.02346.x
409. Gisder, S., Schüler, V., Horchler, L. L., Groth, D., & Genersch, E. (2017). Long-term temporal trends of Nosema spp. Infection prevalence in Northeast Germany: Continuous spread of Nosema ceranae, an emerging pathogen of honey bees (Apis mellifera), but no general replacement of Nosema apis. Frontiers in Cellular and Infection Microbiology, 7(JUL). https://doi.org/10.3389/fcimb.2017.00301

506. Khan, S. U., Anjum, S. I., Ansari, M. J., Khan, M. H. U., Kamal, S., Rahman, K., Shoaib, M., Man, S., Khan, A. J., Khan, S. U., & Khan, D. (2019). Antimicrobial potentials of medicinal plant’s extract and their derived silver nanoparticles: A focus

on honey bee pathogen. Saudi Journal of Biological Sciences, 26(7), 1815–1834. https://doi.org/10.1016/j.sjbs.2018.02.010
507. Khezri, M., Moharrami, M., Modirrousta, H., Torkaman, M., Salehi, S., Rokhzad, B., & Khanbabai, H. (2018). Molecular detection of nosema ceranae in the apiaries of Kurdistan province, Iran. Veterinary Research Forum, 9(3), 273–278. https://doi.org/10.30466/vrf.2018.32086
508. Kielmanowicz, M. G., Inberg, A., Lerner, I. M., Golani, Y., Brown, N., Turner, C. L., Hayes, G. J. R., & Ballam, J. M. (2015). Prospective Large-Scale Field Study Generates Predictive Model Identifying Major Contributors to Colony Losses. PLoS Pathogens, 11(4). https://doi.org/10.1371/journal.ppat.1004816
509. Kim, D. Y., & Lee, J. K. (2022). Development of monoclonal antibodies against spores of Nosema ceranae for the diagnosis of nosemosis. Journal of Apicultural Research. https://doi.org/10.1080/00218839.2022.2053033
510. Kim, D. Y., Maeng, S., Cho, S.-J., Park, H. J., Kim, K., Lee, J. K., & Srinivasan, S. (2023). The Ascosphaera apis Infection (Chalkbrood Disease) Alters the Gut Bacteriome Composition of the Honeybee. Pathogens, 12(5). https://doi.org/10.3390/pathogens12050734
511. Kim, D.-J., Yun, H.-G., Kim, I.-H., Gwak, W.-S., & Woo, S.-D. (2017). Efficient method for the rapid purification of Nosema ceranae spores. Mycobiology, 45(3), 204–208. https://doi.org/10.5941/MYCO.2017.45.3.204
512. Kipkoech, A., Okwaro, L. A., Muli, E., & Lattorff, H. M. G. (2023). Occurrence and distribution of Nosema ceranae in honey bee colonies in the Comoros Islands. Journal of Apicultural Research, 62(5), 1197–1206. https://doi.org/10.1080/00218839.2023.2221563
513. Klassen, S. S., Vanblyderveen, W., Eccles, L., Kelly, P. G., Borges, D., Goodwin, P. H., Petukhova, T., Wang, Q., & Guzman-Novoa, E. (2021). Nosema ceranae infections in honey bees (Apis mellifera) treated with pre/probiotics and impacts on colonies in the field. Veterinary Sciences, 8(6). https://doi.org/10.3390/vetsci8060107
514. Klee, J., Besana, A. M., Genersch, E., Gisder, S., Nanetti, A., Tam, D. Q., Chinh, T. X., Puerta, F., Ruz, J. M., Kryger, P., Message, D., Hatjina, F., Korpela, S., Fries, I., & Paxton, R. J. (2007). Widespread dispersal of the microsporidian Nosema ceranae, an emergent pathogen of the western honey bee, Apis mellifera. Journal of Invertebrate Pathology, 96(1), 1–10. https://doi.org/10.1016/j.jip.2007.02.014
515. Klee, J., Tek Tay, W., & Paxton, R. J. (2006). Specific and sensitive detection of Nosema bombi (Microsporidia: Nosematidae) in bumble bees (Bombus spp.; Hymenoptera: Apidae) by PCR of partial rRNA gene sequences. Journal of Invertebrate Pathology, 91(2), 98–104. https://doi.org/10.1016/j.jip.2005.10.012
516. Kunat-Budzyńska, M., Budzyński, M., Schulz, M., Strachecka, A., Gancarz, M., Rusinek, R., & Ptaszyńska, A. A. (2022). Natural Substances, Probiotics, and Synthetic Agents in the Treatment and Prevention of Honeybee Nosemosis. Pathogens, 11(11). https://doi.org/10.3390/pathogens11111269
517. Kurze, C., Routtu, J., & Moritz, R. F. A. (2016). Parasite resistance and tolerance in honeybees at the individual and social level. Zoology, 119(4), 290–297. https://doi.org/10.1016/j.zool.2016.03.007
518. Kyle, B., Lee, K., & Pernal, S. F. (2021). Epidemiology and Biosecurity for Veterinarians Working with Honey bees (Apis mellifera). Veterinary Clinics of North

America - Food Animal Practice, 37(3), 479–490. https://doi.org/10.1016/j.cvfa.2021.06.004
519. Lage, V. M. G. B., Santana, C. D., Patrocínio, E., Noronha, R. P., de Melo, R. L., Barbosa, C. J., & Lima, S. T. C. (2022). Prevalence of Nosema ceranae in apiculture regions of Bahia State, Brazil. Ciencia Rural, 52(9). https://doi.org/10.1590/0103-8478cr20210473
520. Lannutti, L., Gonzales, F. N., Dus Santos, M. J., Florin-Christensen, M., & Schnittger, L. (2022). Molecular Detection and Differentiation of Arthropod, Fungal, Protozoan, Bacterial and Viral Pathogens of Honeybees. Veterinary Sciences, 9(5). https://doi.org/10.3390/vetsci9050221
521. Lannutti, L., Mira, A., Basualdo, M., Rodriguez, G., Erler, S., Silva, V., Gisder, S., Genersch, E., Florin-Christensen, M., & Schnittger, L. (2020). Development of a loop-mediated isothermal amplification (LAMP) and a direct LAMP for the specific detection of Nosema ceranae, a parasite of honey bees. Parasitology Research, 119(12), 3947–3956. https://doi.org/10.1007/s00436-020-06915-w
522. Lecocq, A., Jensen, A. B., Kryger, P., & Nieh, J. C. (2016). Parasite infection accelerates age polyethism in young honey bees. Scientific Reports, 6. https://doi.org/10.1038/srep22042
523. Lee, D.-W. (2013). Detection of a microsporidium, nosema ceranae, from field population of the bumblebee, bombus terrestris, via quantitative real-time PCR. Korean Journal of Microbiology, 49(3), 270–274. https://doi.org/10.7845/kjm.2013.3052
524. Li, J. L., Chen, W. F., Wu, J., Peng, W. J., An, J. D., Schmid-Hempel, P., & Schmid-Hempel, R. (2012). Diversity of Nosema associated with bumblebees (Bombus spp.) from China. INTERNATIONAL JOURNAL FOR PARASITOLOGY, 42(1), 49–61. https://doi.org/10.1016/j.ijpara.2011.10.005
525. Li, J., Qin, H., Wu, J., Sadd, B. M., Wang, X., Evans, J. D., Peng, W., & Chen, Y. (2012). The Prevalence of Parasites and Pathogens in Asian Honeybees Apis cerana in China. PLoS ONE, 7(11). https://doi.org/10.1371/journal.pone.0047955
526. Li, W., Evans, J. D., Li, J., Su, S., Hamilton, M., & Chen, Y. (2017). Spore load and immune response of honey bees naturally infected by Nosema ceranae. Parasitology Research, 116(12), 3265–3274. https://doi.org/10.1007/s00436-017-5630-8
527. Li, Z., Hao, Y., Wang, L., Xiang, H., & Zhou, Z. (2014). Genome-wide identification and comprehensive analyses of the kinomes in four pathogenic microsporidia species. PLoS ONE, 9(12). https://doi.org/10.1371/journal.pone.0115890
528. Lim, H. C., Lambrecht, D., Forkner, R. E., & Roulston, T. (2023). Minimal sharing of nosematid and trypanosomatid parasites between honey bees and other bees, but extensive sharing of Crithidia between bumble and mason bees. Journal of Invertebrate Pathology, 198. https://doi.org/10.1016/j.jip.2023.107933
529. Liu, S., Wang, L., Guo, J., Li, J., & Xu, L. (2017). The variation of pathogens, parasites and symbionts in migratory honeybees (Apis mellifera ligustica). Scientia Agricultura Sinica, 50(5), 951–958. https://doi.org/10.3864/j.issn.0578-1752.2017.05.018
530. Lodesani, M., Costa, C., Besana, A., Dall'Olio, R., Franceschetti, S., Tesoriero, D., & Vaccari, G. (2014). Impact of control strategies for Varroa destructor on colony

survival and health in northern and central regions of Italy. Journal of Apicultural Research, 53(1), 155–164. https://doi.org/10.3896/IBRA.1.53.1.17
531. Lopes, A. R., Martín-Hernández, R., Higes, M., Segura, S. K., Henriques, D., & Pinto, M. A. (2022). Colonisation Patterns of Nosema ceranae in the Azores Archipelago. Veterinary Sciences, 9(7). https://doi.org/10.3390/vetsci9070320
532. Lopes, A. R., Martín-Hernández, R., Higes, M., Segura, S. K., Henriques, D., & Pinto, M. A. (2023). First detection of Nosema ceranae in honey bees (Apis mellifera L.) of the Macaronesian archipelago of Madeira. Journal of Apicultural Research, 62(3), 514–517. https://doi.org/10.1080/00218839.2023.2172835
533. Łopieńska-Biernat, E., Sokół, R., Michalczyk, M., Żółtowska, K., & Stryiński, R. (2017). Biochemical status of feral honey bees (Apis mellifera) infested with various pathogens. Journal of Apicultural Research, 56(5), 606–615. https://doi.org/10.1080/00218839.2017.1343020
534. Łoś, A., Skórka, P., Strachecka, A., Winiarczyk, S., Winiarczyk, M., & Wolski, D. (2020). The associations among the breeding performance of Osmia bicornis L. (Hymenoptera: Megachilidae), burden of pathogens and nest parasites along urbanisation gradient. Science of the Total Environment, 710. https://doi.org/10.1016/j.scitotenv.2019.135520
535. Lourenço, A. P., Guidugli-Lazzarini, K. R., de Freitas, N. H. A., Message, D., Bitondi, M. M. G., Simões, Z. L. P., & Teixeira, É. W. (2021). Immunity and physiological changes in adult honey bees (Apis mellifera) infected with Nosema ceranae: The natural colony environment. Journal of Insect Physiology, 131. https://doi.org/10.1016/j.jinsphys.2021.104237
536. Lu, M.-C. (2018). Beekeeping on Taiwan Island. In Asian Beekeeping in the 21st Century (pp. 159–173). https://doi.org/10.1007/978-981-10-8222-1_7
537. Luis, A. R., García, C. A. Y., Invernizzi, C., Branchiccela, B., Piñeiro, A. M. P., Morfi, A. P., Zunino, P., & Antúnez, K. (2020). Nosema ceranae and RNA viruses in honey bee populations of Cuba. Journal of Apicultural Research, 59(4), 468–471. https://doi.org/10.1080/00218839.2020.1749451
538. Ma, Z., Li, C., Pan, G., Li, Z., Han, B., Xu, J., Lan, X., Chen, J., Yang, D., Chen, Q., Sang, Q., Ji, X., Li, T., Long, M., & Zhou, Z. (2013). 1DUMMY Genome-wide transcriptional response of silkworm (Bombyx mori) to infection by the microsporidian Nosema bombycis. PLoS ONE, 8(12). https://doi.org/10.1371/journal.pone.0084137
539. Ma, Z., Wang, Y., Huang, Z., Cheng, S., Xu, J., & Zhou, Z. (2021). Isolation of protein-free chitin spore coats of Nosema ceranae and its application to screen the interactive spore wall proteins. Archives of Microbiology, 203(5), 2727–2733. https://doi.org/10.1007/s00203-021-02214-9
540. Macías-Macías, J. O., Tapia-Rivera, J. C., De la Mora, A., Tapia-González, J. M., Contreras-Escareño, F., Petukhova, T., Morfin, N., & Guzman-Novoa, E. (2020). Nosema ceranae causes cellular immunosuppression and interacts with thiamethoxam to increase mortality in the stingless bee Melipona colimana. Scientific Reports, 10(1). https://doi.org/10.1038/s41598-020-74209-3
541. Maksong, S., Yemor, T., & Yanmanee, S. (2019). Detection of nosemosis in European honeybees (Apis mellifera) on honeybees farm at Kanchanaburi, Thailand. IOP Conference Series: Materials Science and Engineering, 639(1). https://doi.org/10.1088/1757-899X/639/1/012048

552. Martín-Hernández, R., Botías, C., Bailón, E. G., Martínez-Salvador, A., Prieto, L., Meana, A., & Higes, M. (2012). Microsporidia infecting Apis mellifera: Coexistence or competition. Is Nosema ceranae replacing Nosema apis? Environmental Microbiology, 14(8), 2127–2138. https://doi.org/10.1111/j.1462-2920.2011.02645.x
553. Martín-Hernández, R., Meana, A., Prieto, L., Salvador, A. M., Garrido-Bailón, E., & Higes, M. (2007). Outcome of colonization of Apis mellifera by Nosema ceranae. Applied and Environmental Microbiology, 73(20), 6331–6338. https://doi.org/10.1128/AEM.00270-07
554. Maside, X., Gómez-Moracho, T., Jara, L., Martín-Hernández, R., De La Rúa, P., Higes, M., & Bartolomé, C. (2015). Population genetics of Nosema apis and Nosema ceranae: One host (apis mellifera) and two different histories. PLoS ONE, 10(12). https://doi.org/10.1371/journal.pone.0145609
555. Matović, K., Vidanović, D., Manić, M., Stojiljković, M., Radojičić, S., Debeljak, Z., Šekler, M., & Ćirić, J. (2020). Twenty-five-year study of Nosema spp. In honey bees (Apis mellifera) in Serbia. Saudi Journal of Biological Sciences, 27(1), 518–523. https://doi.org/10.1016/j.sjbs.2019.11.012
556. Matthijs, S., de Waele, V., Vandenberge, V., Verhoeven, B., Evers, J., Brunain, M., Saegerman, C., de Winter, P. J. J., Roels, S., de Graaf, D. C., & de Regge, N. (2020). Nationwide screening for bee viruses and parasites in belgian honey bees. Viruses, 12(8). https://doi.org/10.3390/v12080890
557. Mayack, C., Broadrup, R. L., Schick, S. J., Eppley, E. J., Khan, Z., & MacHerone, A. (2021). Increased alarm pheromone component is associated with Nosema ceranae infected honeybee colonies. Royal Society Open Science, 8(4). https://doi.org/10.1098/rsos.210194
558. Mayack, C., Cook, S. E., Niño, B. D., Rivera, L., Niño, E. L., & Seshadri, A. (2023). Poor Air Quality Is Linked to Stress in Honeybees and Can Be Compounded by the Presence of Disease. Insects, 14(8). https://doi.org/10.3390/insects14080689
559. Mayack, C., & Hakanoğlu, H. (2022). Honey Bee Pathogen Prevalence and Interactions within the Marmara Region of Turkey. Veterinary Sciences, 9(10). https://doi.org/10.3390/vetsci9100573
560. Mccallum, R., Olmstead, S., Shaw, J., & Glasgow, K. (2021). Evaluating Efficacy of Fumagilin-B®Against Nosemosis and Tracking Seasonal Trends of Nosema spp. In Nova Scotia Honey Bee Colonies. Journal of Apicultural Science, 64(2), 277–286. https://doi.org/10.2478/jas-2020-0025
561. McInnis, J. L., Williams, T., Chuang, Y.-C., & Gregg, D. A. (2020). Replication of Invertebrate Iridescent Virus 6 (IIV-6) in European Honey Bees—Potential Involvement in Colony Collapse Disorder. Southwestern Entomologist, 45(2), 335–340. https://doi.org/10.3958/059.045.0201
562. McNamara-Bordewick, N. K., McKinstry, M., & Snow, J. W. (2019). Robust transcriptional response to heat shock impacting diverse cellular processes despite lack of heat shock factor in microsporidia. mSphere, 4(3). https://doi.org/10.1128/mSphere.00219-19
563. Meana, A., Llorens-Picher, M., Euba, A., Bernal, J. L., Bernal, J., García-Chao, M., Dagnac, T., Castro-Hermida, J. A., González-Porto, A. V., Higes, M., & Martín-Hernández, R. (2017). Risk factors associated with honey bee colony loss in apiaries

in Galicia, NW Spain. Spanish Journal of Agricultural Research, 15(1). https://doi.org/10.5424/sjar/2017151-9652
564. Meana, A., Martín-Hernández, R., & Higes, M. (2010). The reliability of spore counts to diagnose Nosema ceranae infections in honey bees. Journal of Apicultural Research, 49(2), 212–214. https://doi.org/10.3896/IBRA.1.49.2.12
565. Mederle, N., Lobo, M. L., Morariu, S., Morariu, F., Darabus, G., Mederle, O., & Matos, O. (2018a). Microscopic and molecular detection of nosema ceranae in honeybee apis mellifera L. from Romania. Revista de Chimie, 69(12), 3761–3772. https://doi.org/10.37358/rc.18.12.6837
566. Mederle, N., Lobo, M. L., Morariu, S., Morariu, F., Darabus, G., Mederle, O., & Matos, O. (2018b). Microscopic and Molecular Detection of Nosema ceranae in Honeybee Apis mellifera L. from Romania Status on pathogen worldwide distribution. REVISTA DE CHIMIE, 69(12), 3761–3772.
567. Medici, S. K., Sarlo, E. G., Porrini, M. P., Braunstein, M., & Eguaras, M. J. (2012). Genetic variation and widespread dispersal of Nosema ceranae in Apis mellifera apiaries from Argentina. Parasitology Research, 110(2), 859–864. https://doi.org/10.1007/s00436-011-2566-2
568. Meena, K. R., & Kanwar, S. S. (2015). Lipopeptides as the antifungal and antibacterial agents: Applications in food safety and therapeutics. BioMed Research International, 2015. https://doi.org/10.1155/2015/473050
569. Meftahi, B., Yaghfouri, S., Mosazadeh, S., Sheibani Tezerji, R., Fakhrabadipour, M., Javdan, E., & Razmi, G. R. (2023). Parasitological and molecular study of nosemosis in migratory apiaries in Hormozgan Province, southern Iran. Journal of Entomological Society of Iran, 43(4), 383–392. https://doi.org/10.61186/jesi.43.4.6
570. Meixner, M. D., Kryger, P., & Costa, C. (2015). Effects of genotype, environment, and their interactions on honey bee health in Europe. Current Opinion in Insect Science, 10, 177–184. https://doi.org/10.1016/j.cois.2015.05.010
571. Meixner, M. D., & Uzunov, A. (2019). Genotype-environment-interactions and the occurrence of honey bee diseases affect the survival of honey bee colonies – summary from a pan-European experiment. Berliner Und Munchener Tierarztliche Wochenschrift, 132(1–2), 16–25. https://doi.org/10.2376/0005-9366-18015
572. Menail, A. H., Piot, N., Meeus, I., Smagghe, G., & Loucif-Ayad, W. (2016). Large pathogen screening reveals first report of Megaselia scalaris (Diptera: Phoridae) parasitizing Apis mellifera intermissa (Hymenoptera: Apidae). Journal of Invertebrate Pathology, 137, 33–37. https://doi.org/10.1016/j.jip.2016.04.007
573. Mendoza, Y., Antúnez, K., Branchiccela, B., Anido, M., Santos, E., & Invernizzi, C. (2014). Nosema ceranae and RNA viruses in European and Africanized honeybee colonies (Apis mellifera) in Uruguay. Apidologie, 45(2), 224–234. https://doi.org/10.1007/s13592-013-0241-6
574. Mendoza, Y., Diaz-Cetti, S., Ramallo, G., Santos, E., Porrini, M., & Invernizzi, C. (2017). Nosema ceranae Winter Control: Study of the Effectiveness of Different Fumagillin Treatments and Consequences on the Strength of Honey Bee (Hymenoptera: Apidae) Colonies. Journal of Economic Entomology, 110(1), 1–5. https://doi.org/10.1093/jee/tow228
575. Mendoza, Y., Santos, E., Antunez, K., & Invernizzi, C. (2014). Bidirectional selection of Apis mellifera (Hymenoptera: Apidae) for increased resistance and

susceptibility to nosemosis. REVISTA DE LA SOCIEDAD ENTOMOLOGICA ARGENTINA, 73(1–2), 65–69.
576. Michalczyk, M., Bancerz-Kisiel, A., & Sokół, R. (2020). Lotmaria passim as third parasite gastrointestinal tract of honey bees living in tree trunk. Journal of Apicultural Science, 64(1), 143–151. https://doi.org/10.2478/jas-2020-0012
577. Michalczyk, M., & Sokół, R. (2014). Nosemosis in honey bees. Polish Journal of Natural Sciences, 29(1), 91–99.
578. Michalczyk, M., & Sokół, R. (2018). Estimation of the influence of selected products on co-infection with N. apis/N. ceranae in Apis mellifera using real-time PCR. Invertebrate Reproduction and Development, 62(2), 92–97. https://doi.org/10.1080/07924259.2018.1433726
579. Michalczyk, M., Sokół, R., & Bancerz-Kisiel, A. (2022). Coexistence between selected pathogens in honey bee workers. Journal of Apicultural Research, 61(3), 346–351. https://doi.org/10.1080/00218839.2021.1994261
580. Michalczyk, M., SokóŁ, R., & Koziatek, S. (2016). Evaluation of the effectiveness of selected treatments of Nosema spp. Infection by the hemocytometric method and duplex PCR. Acta Veterinaria, 66(1), 115–124. https://doi.org/10.1515/acve-2016-0009
581. Michalczyk, M., Sokół, R., & Szczerba-Turek, A. (2013). Phylogenetic analysis of Nosema apis and Nosema ceranae small subunit 16S rRNA in honey bees (Apis mellifera) from north-eastern Poland. Medycyna Weterynaryjna, 69(12), 733–735.
582. Michalczyk, M., Sokół, R., Szczerba-Turek, A., & Bancerz-Kisiel, A. (2011). A comparison of the effectiveness of the microscopic method and the multiplex PCR method in identifying and discriminating the species of Nosema spp. Spores in worker bees (Apis mellifera) from winter hive debris. Polish Journal of Veterinary Sciences, 14(3), 385–391. https://doi.org/10.2478/v10181-011-0058-z
583. Modirrousta, H., Moharrami, M., & Mansouri, M. A. (2014). Retrospective study of the Nosema ceranae infection of honey bee colonies in Iran (2004-2013). Archives of Razi Institute, 69(2), 197–200.
584. Moeini, S., Malekifard, F., & Tavassoli, M. (2022). Identification of the Nosema spp., a microsporidian parasite isolated from the honey bees (Apis mellifera) and its association with honey bee colony losses in apiaries of Iran. Journal of the Hellenic Veterinary Medical Society, 73, 3667–3672. https://doi.org/10.12681/jhvms.25393
585. Mohammadian, B., Bokaie, S., Moharrami, M., Nabian, S., & Forsi, M. (2018). Distribution of nosema spp. In climatic regions of Iran. Veterinary Research Forum, 9(3), 259–263. https://doi.org/10.30466/vrf.2018.32082
586. Mohammadian, B., Bokaie, S., Moharrami, M., Nabian, S., & Forsi, M. (2019). Prevalence of honeybee colony collapse disorder and its relation to Nosema spp. And climate in apiaries of Iran. Journal of Veterinary Research, 74(1), 11–18. https://doi.org/10.22059/jvr.2017.235690.2649
587. Mouret, C., Lambert, O., Piroux, M., Beaudeau, F., Provost, B., Benet, P., Colin, M.-E., & L'Hostis, M. (2013). Prevalence of 12 infectious agents in field colonies of 18 apiaries in western France. Revue de Medecine Veterinaire, 164(12), 578–582.

601. Odemer, R. (2013). Nosema ceranae, a new threat to honey bees (Apis mellifera L.)? Tierarztliche Umschau, 68(4), 126–131.
602. Odnosum, H. V. (2017). Distribution of the Nosema ceranae (Microspora, Nosematidae) in the Apiaries in Ukraine. Vestnik Zoologii, 51(2), 161–166. https://doi.org/10.1515/vzoo-2017-0022
603. Oguz, B., Karapinar, Z., Dinçer, E., & Deĝer, M. S. (2017). Molecular detection of Nosema spp. And black queen-cell virus in honeybees in Van Province, Turkey. Turkish Journal of Veterinary and Animal Sciences, 41(2), 221–227. https://doi.org/10.3906/vet-1604-92
604. Oliver, R. (2008). The 'Nosema twins'—Part IV - Treatment. AMERICAN BEE JOURNAL, 148(3), 249–252.
605. Oliver, R. (2009). Nosema ceranae: Kiss of Death or Much Ado About Nothing? AMERICAN BEE JOURNAL, 149(8), 759–764.
606. Oliver, R. (2012). Sick bees—Part 17B - Nosema- the smoldering epidemic. American Bee Journal, 152(4), 375–380.
607. Oliver, R. (2015). The seasonality of nosema ceranae. American Bee Journal, 155(4), 447–449.
608. Oliver, R., & Dunbar, B. (2012). Nosema ceranae and honey production in healthy colonies. American Bee Journal, 152(11), 1071–1072.
609. Ongus, J. R., Fombong, A. T., Irungu, J., Masiga, D., & Raina, S. (2018). Prevalence of common honey bee pathogens at selected apiaries in Kenya, 2013/2014. International Journal of Tropical Insect Science, 38(1), 58–70. https://doi.org/10.1017/S1742758417000212
610. Ostroverkhova, N. V. (2020). Prevalence Of Nosema Ceranae (Microsporidia) In The Apis Mellifera Mellifera Bee Colonies From Long Time Isolated Apiaries Of Siberia. Far Eastern Entomologist, 407, 8–20. https://doi.org/10.25221/fee.407.2
611. Ostroverkhova, N. V. (2021). Association between the microsatellite Ap243, AC117 and SV185 polymorphisms and Nosema disease in the dark forest bee Apis mellifera mellifera. Veterinary Sciences, 8(1), 1–15. https://doi.org/10.3390/VETSCI8010002
612. Ostroverkhova, N. V., Konusova, O. L., Kucher, A. N., Kireeva, T. N., & Rosseykina, S. A. (2020). Prevalence of the microsporidian nosema spp. In honey bee populations (apis mellifera) in some ecological regions of North Asia. Veterinary Sciences, 7(3). https://doi.org/10.32890/JICT2018.17.2.8252
613. Ostroverkhova, N. V., Kucher, A. N., Golubeva, E. P., Rosseykina, S. A., & Konusova, O. L. (2019). Study of Nosema spp. In the Tomsk region, Siberia: Co-infection is widespread in honeybee colonies. Far Eastern Entomologist, 378, 12–22. https://doi.org/10.25221/FEE.378.3
614. Ozgor, E., Celebier, I., Ulusoy, M., & Keskin, N. (2017). First detection of nosema ceranae and nosema apis in greater wax moth galleria mellonella. Journal of Apicultural Science, 61(2), 185–192. https://doi.org/10.1515/JAS-2017-0015
615. Özkırım, A., Schiesser, A., & Keskin, N. (2019). Dynamics of nosema apis and nosema ceranae co-infection seasonally in honey bee (Apis mellifera L.) colonies. Journal of Apicultural Science, 63(1), 41–48. https://doi.org/10.2478/jas-2019-0001
616. Pacini, A., Giacobino, A., Molineri, A., Bulacio Cagnolo, N., Aignasse, A., Zago, L., Mira, A., Izaguirre, M., Schnittger, L., Merke, J., Orellano, E., Bertozzi, E., Pietronave, H., & Signorini, M. (2016). Risk factors associated with the abundance of

Nosema spp. In apiaries located in temperate and subtropical conditions after honey harvest. Journal of Apicultural Research, 55(4), 342–350. https://doi.org/10.1080/00218839.2016.1245396
617. Pacini, A., Mira, A., Molineri, A., Giacobino, A., Bulacio Cagnolo, N., Aignasse, A., Zago, L., Izaguirre, M., Merke, J., Orellano, E., Bertozzi, E., Pietronave, H., Russo, R., Scannapieco, A., Lanzavecchia, S., Schnittger, L., & Signorini, M. (2016). Distribution and prevalence of Nosema apis and N. ceranae in temperate and subtropical eco-regions of Argentina. Journal of Invertebrate Pathology, 141, 34–37. https://doi.org/10.1016/j.jip.2016.11.002
618. Pacini, A., Molineri, A., Antúnez, K., Cagnolo, N. B., Merke, J., Orellano, E., Bertozzi, E., Zago, L., Aignasse, A., Pietronave, H., Rodríguez, G., Palacio, M. A., Signorini, M., & Giacobino, A. (2021). Environmental conditions and beekeeping practices associated with Nosema ceranae presence in Argentina. Apidologie, 52(2), 400–417. https://doi.org/10.1007/s13592-020-00831-9
619. Pajuelo, A. G., Torres, C., & Bermejo, F. J. O. (2008). Colony losses: A double blind trial on the influence of supplementary protein nutrition and preventative treatment with fumagillin against Nosema ceranae. Journal of Apicultural Research, 47(1), 84–86. https://doi.org/10.1080/00218839.2008.11101429
620. Pan, G., Xu, J., Li, T., Xia, Q., Liu, S.-L., Zhang, G., Li, S., Li, C., Liu, H., Yang, L., Liu, T., Zhang, X., Wu, Z., Fan, W., Dang, X., Xiang, H., Tao, M., Li, Y., Hu, J., … Zhou, Z. (2013). Comparative genomics of parasitic silkworm microsporidia reveal an association between genome expansion and host adaptation. BMC Genomics, 14(1). https://doi.org/10.1186/1471-2164-14-186
621. Papežíková, I., Palíková, M., Syrová, E., Zachová, A., Somerlíková, K., Kováčová, V., & Pecková, L. (2020). Effect of feeding honey bee (Apis mellifera Hymenoptera: Apidae) colonies with honey, sugar solution, inverted sugar, and wheat starch syrup on nosematosis prevalence and intensity. Journal of Economic Entomology, 113(1), 26–33. https://doi.org/10.1093/jee/toz251
622. Papini, R., Mancianti, F., Canovai, R., Cosci, F., Rocchigiani, G., Benelli, G., & Canale, A. (2017). Prevalence of the microsporidian Nosema ceranae in honeybee (Apis mellifera) apiaries in Central Italy. Saudi Journal of Biological Sciences, 24(5), 979–982. https://doi.org/10.1016/j.sjbs.2017.01.010
623. Parvanov, P., & Rusenova, N. (2014). Etiology and clinicoepidemiological profile of apiaries with colony collapse disorder-like symptoms in Bulgaria. Bulgarian Journal of Veterinary Medicine, 17(3), 199–206.
624. Paxton, R. J. (2010). Does infection by Nosema ceranae cause 'colony Collapse Disorder' in honey bees (Apis mellifera)? Journal of Apicultural Research, 49(1), 80–84. https://doi.org/10.3896/IBRA.1.49.1.11
625. Pelin, A., Selman, M., Aris-Brosou, S., Farinelli, L., & Corradi, N. (2015). Genome analyses suggest the presence of polyploidy and recent human-driven expansions in eight global populations of the honeybee pathogen Nosema ceranae. Environmental Microbiology, 17(11), 4443–4458. https://doi.org/10.1111/1462-2920.12883
626. Pereira, K. S., Meeus, I., & Smagghe, G. (2019). Honey bee-collected pollen is a potential source of Ascosphaera apis infection in managed bumble bees. Scientific Reports, 9(1). https://doi.org/10.1038/s41598-019-40804-2

639. Ponkit, R., Naree, S., Pichayangkura, R., Beaurepaire, A., Paxton, R. J., Mayack, C. L., & Suwannapong, G. (2023). Chito-Oligosaccharide and Propolis Extract of Stingless Bees Reduce the Infection Load of Nosema ceranae in Apis dorsata (Hymenoptera: Apidae). Journal of Fungi, 9(1). https://doi.org/10.3390/jof9010020
640. Porrini, C., Mutinelli, F., Bortolotti, L., Granato, A., Laurenson, L., Roberts, K., Gallina, A., Silvester, N., Medrzycki, P., Renzi, T., Sgolastra, F., & Lodesani, M. (2016). The status of honey bee health in Italy: Results from the nationwide bee monitoring network. PLoS ONE, 11(5). https://doi.org/10.1371/journal.pone.0155411
641. Porrini, L. P., Porrini, M. P., Garrido, P. M., Principal, J., Barrios Suarez, C. J., Bianchi, B., Fernandez Iriarte, P. J., & Eguaras, M. J. (2017). First identification of nosema ceranae (Microsporidia) infecting apis mellifera in Venezuela. Journal of Apicultural Science, 61(1), 149–152. https://doi.org/10.1515/JAS-2017-0010
642. Porrini, M. P., Porrini, L. P., Garrido, P. M., de Melo e Silva Neto, C., Porrini, D. P., Muller, F., Nuñez, L. A., Alvarez, L., Iriarte, P. F., & Eguaras, M. J. (2017). Nosema ceranae in South American Native Stingless Bees and Social Wasp. Microbial Ecology, 74(4), 761–764. https://doi.org/10.1007/s00248-017-0975-1
643. Prados, E. A., Hernández, R. M., & Pascual, M. H. (2018). Results of a monitoring program of pesticide residues in Beebread in Spain. Using Toxic unit approach to identify scenarios of risk for management programs. In P. A. Oomen & J. Pistorius (Eds.), HAZARDS OF PESTICIDES TO BEES (Vol. 462, Issue 13th International Symposium of the ICP-PR-Bee-Protection-Group on Hazards of Pesticides to Bees, pp. 194–194). https://doi.org/10.5073/jka.2018.462.061
644. Ptaszyńska, A. A., Borsuk, G., Anusiewicz, M., & Mułenko, W. (2012). Location of Nosema spp. Spores within the body of the honey bee. Medycyna Weterynaryjna, 68(10), 618–621.
645. Ptaszyńska, A. A., Borsuk, G., Mulenko, W., & Demetraki-Paleolog, J. (2014). Differentiation of Nosema apis and Nosema ceranae spores under Scanning Electron Microscopy (SEM). Journal of Apicultural Research, 53(5), 537–544. https://doi.org/10.3896/IBRA.1.53.5.02
646. Ptaszyńska, A. A., Borsuk, G., Mułenko, W., & Olszewski, K. (2012). Monitoring of nosemosis in the lublin region and preliminary morphometric studies of nosema spp. Spores. Medycyna Weterynaryjna, 68(10), 622–625.
647. Ptaszyńska, A. A., Borsuk, G., Woźniakowski, G., Gnat, S., & Małek, W. (2014). Loop-mediated isothermal amplification (LAMP) assays for rapid detection and differentiation of Nosema apis and N. ceranae in honeybees. FEMS Microbiology Letters, 357(1), 40–48. https://doi.org/10.1111/1574-6968.12521
648. Ptaszyńska, A. A., Gancarz, M., Hurd, P. J., Borsuk, G., Wiącek, D., Nawrocka, A., Strachecka, A., Załuski, D., & Paleolog, J. (2018). Changes in the bioelement content of summer and winter western honeybees (Apis mellifera) induced by Nosema ceranae infection. PLoS ONE, 13(7). https://doi.org/10.1371/journal.pone.0200410
649. Ptaszyńska, A. A., Latoch, P., Hurd, P. J., Polaszek, A., Michalska-Madej, J., Grochowalski, Ł., Strapagiel, D., Gnat, S., Załuski, D., Gancarz, M., Rusinek, R., Krutmuang, P., Martín Hernández, R., Higes Pascual, M., & Starosta, A. L. (2021). Amplicon sequencing of variable 16s rrna from bacteria and its2 regions from fungi and plants, reveals honeybee susceptibility to diseases results from their forage

availability under anthropogenic landscapes. Pathogens, 10(3). https://doi.org/10.3390/pathogens10030381
650. Ptaszyńska, A. A., & Mulenko, W. (2013). Selected aspects of the structure, development, taxonomy and biology of microsporidian parasites belonging to the genus Nosema. Medycyna Weterynaryjna, 69(12), 716–725.
651. Punko, R. N., Currie, R. W., Nasr, M. E., & Hoover, S. E. (2021). Epidemiology of Nosema spp. And the effect of indoor and outdoor wintering on honey bee colony population and survival in the Canadian Prairies. PLoS ONE, 16(10 October). https://doi.org/10.1371/journal.pone.0258801
652. Punko, R. N., Currie, R. W., Nasr, M. E., & Hoover, S. E. (2023). Effect of Fumagilin-B treatment timing on nosema (Vairimorpha spp.; Microspora: Nosematidae) abundance and honey bee (Hymenoptera: Apidae) colonies under winter management in the Canadian Prairies. Journal of Economic Entomology, 116(3), 651–661. https://doi.org/10.1093/jee/toad066
653. Purkiss, T., & Lach, L. (2019). Pathogen spillover from Apis mellifera to a stingless bee. Proceedings of the Royal Society B: Biological Sciences, 286(1908). https://doi.org/10.1098/rspb.2019.1071
654. Ramos-Cuellar, A. K., De la Mora, A., Contreras-Escareño, F., Morfin, N., Tapia-González, J. M., Macías-Macías, J. O., Petukhova, T., Correa-Benítez, A., & Guzman-Novoa, E. (2022). Genotype, but Not Climate, Affects the Resistance of Honey Bees (Apis mellifera) to Viral Infections and to the Mite Varroa destructor. Veterinary Sciences, 9(7). https://doi.org/10.3390/vetsci9070358
655. Ran, M., Shi, Y., Li, B., Xiang, H., Tao, M., Meng, X., Li, T., Li, C., Bao, J., Pan, G., & Zhou, Z. (2023). Genome-Wide Characterization and Comparative Genomic Analysis of the Serpin Gene Family in Microsporidian Nosema bombycis. International Journal of Molecular Sciences, 24(1). https://doi.org/10.3390/ijms24010550
656. Rangel, J., Baum, K., Rubink, W. L., Coulson, R. N., Johnston, J. S., & Traver, B. E. (2016). Prevalence of Nosema species in a feral honey bee population: A 20-year survey. Apidologie, 47(4), 561–571. https://doi.org/10.1007/s13592-015-0401-y
657. Rangel, J., Gonzalez, A., Stoner, M., Hatter, A., & Traver, B. E. (2018). Genetic diversity and prevalence of Varroa destructor, Nosema apis, and N. ceranae in managed honey bee (Apis mellifera) colonies in the Caribbean island of Dominica, West Indies. Journal of Apicultural Research, 57(4), 541–550. https://doi.org/10.1080/00218839.2018.1494892
658. Rangel, J., Traver, B. E., Stevens, G., Howe, M., & Fell, R. D. (2013). Survey for nosema spp. In belize apiaries. Journal of Apicultural Research, 52(2), 62–66. https://doi.org/10.3896/IBRA.1.52.2.12
659. Rangel, J., Traver, B., Stoner, M., Hatter, A., Trevelline, B., Garza, C., Shepherd, T., Seeley, T. D., & Wenzel, J. (2020). Genetic diversity of wild and managed honey bees (Apis mellifera) in Southwestern Pennsylvania, and prevalence of the microsporidian gut pathogens Nosema ceranae and N. apis. Apidologie, 51(5), 802–814. https://doi.org/10.1007/s13592-020-00762-5
660. Ravoet, J., Maharramov, J., Meeus, I., De Smet, L., Wenseleers, T., Smagghe, G., & de Graaf, D. C. (2013). Comprehensive Bee Pathogen Screening in Belgium Reveals Crithidia mellificae as a New Contributory Factor to Winter Mortality. PLoS ONE, 8(8). https://doi.org/10.1371/journal.pone.0072443

685. Sagastume, S., Martín-Hernández, R., Higes, M., & Henriques-Gil, N. (2016). Genotype diversity in the honey bee parasite Nosema ceranae: Multi-strain isolates, cryptic sex or both? BMC Evolutionary Biology, 16(1), 1–11. https://doi.org/10.1186/s12862-016-0797-7
686. Salkova, D., Shumkova, R., Balkanska, R., Palova, N., Neov, B., Radoslavov, G., & Hristov, P. (2022). Molecular Detection of Nosema spp. In Honey in Bulgaria. Veterinary Sciences, 9(1). https://doi.org/10.3390/vetsci9010010
687. Salvarrey, S., Antúnez, K., Arredondo, D., Plischuk, S., Revainera, P., Maggi, M., & Invernizzi, C. (2021). Parasites and RNA viruses in wild and laboratory reared bumble bees Bombus pauloensis (Hymenoptera: Apidae) from Uruguay. PLoS ONE, 16(4 April). https://doi.org/10.1371/journal.pone.0249842
688. Sánchez Collado, J. G., Higes, M., Barrio, L., & Martín-Hernández, R. (2014). Flow cytometry analysis of Nosema species to assess spore viability and longevity. Parasitology Research, 113(5), 1695–1701. https://doi.org/10.1007/s00436-014-3814-z
689. Santrac, V., Granato, A., & Mutinelli, F. (2010). Detection of Nosema ceranae in Apis mellifera from Bosnia and Herzegovina. Journal of Apicultural Research, 49(1), 100–101. https://doi.org/10.3896/IBRA.1.49.1.16
690. Sarlo, E. G., Medici, S. K., Porrini, M. P., Melisa Garrido, P., Floris, I., & Eguaras, M. J. (2011). Comparison between different Fumagillin dosage and evaluation method in the apiary control of Nosemosis type C. Redia, 94, 39–44.
691. Schäfer, M. O., Ritter, W., Pettis, J. S., & Neumann, P. (2010). Winter losses of honeybee colonies (Hymenoptera: Pidae): The role of infestations with Aethina tumida (Coleoptera: Nitidulidae) and Varroa destructor (Parasitiformes: Varroidae). Journal of Economic Entomology, 103(1), 10–16. https://doi.org/10.1603/EC09233
692. Schick, S. J., Broadrup, R. L., Mayack, C., White, H. K., & MacHerone, A. (2018). Integrating GC/TOF exposome profiling and genetic disease screening to provide a holistic perspective on honey bee health. American Laboratory, 50(3), 21–23.
693. Schüler, V., Liu, Y.-C., Gisder, S., Horchler, L., Groth, D., & Genersch, E. (2023). Significant, but not biologically relevant: Nosema ceranae infections and winter losses of honey bee colonies. Communications Biology, 6(1). https://doi.org/10.1038/s42003-023-04587-7
694. Senderskiy, I., Ignatieva, A., & Dolgikh, V. (2020). Vairimorpha (Nosema) ceranae (Opisthosporidia: Microsporidia) in vitro Infection of Sf9 Insect Cell Line as an Experimental Model of Parasite—Host Interrelations. In Y. Tokarev & V. Glupov (Eds.), IV ALL-RUSSIAN PLANT PROTECTION CONGRESS WITH INTERNATIONAL PARTICIPATION: PHYTOSANITARY TECHNOLOGIES IN ENSURING INDEPENDENCE AND COMPETITIVENESS OF THE AGRICULTURAL SECTOR OF RUSSIA (Vol. 18, Issue 4th All-Russian Plant Protection Congress with international participation-Phytosanitary Technologies in Ensuring Independence and Competitiveness of the Agricultural Sector of Russia). https://doi.org/10.1051/bioconf/20201800026
695. Senderskiy, I. V., Ignatieva, A. N., Kireeva, D. S., & Dolgikh, V. V. (2021). Production of polyclonal anti-β-tubulin antibodies and immunodetection of Vairimorpha (Nosema) ceranae (Opisthosporidia: Microsporidia) proliferative stages

in the midguts of Apis mellifera and in the Sf9 cell culture. Protistology, 15(1), 3–9. https://doi.org/10.21685/1680-0826-2021-15-1-1
696. Shafer, A. B. A., Williams, G. R., Shutler, D., Rogers, R. E. L., & Stewart, D. T. (2009). Cophylogeny of nosema (Microsporidia: Nosematidae) and bees (Hymenoptera: Apidae) suggests both cospeciation and a host-switch. Journal of Parasitology, 95(1), 198–203. https://doi.org/10.1645/GE-1724.1
697. Shang, R., Zhu, F., Li, Y., He, P., Qi, J., Chen, Y., Sun, F., Zhang, Y., Wang, Q., & Shen, Z. (2021). Identification and localization of Nup170 in the microsporidian Nosema bombycis. Parasitology Research, 120(6), 2125–2134. https://doi.org/10.1007/s00436-021-07129-4
698. Shao, S. S., Yan, W. Y., & Huang, Q. (2021). Identification of novel miRNAs from the microsporidian parasite Nosema ceranae. Infection, Genetics and Evolution, 93. https://doi.org/10.1016/j.meegid.2021.104930
699. Sharma, D., Katna, S., Sharma, R., Rana, B. S., Sharma, H. K., Bhardwaj, V., & Chauhan, A. (2019). First detection of nosema ceranae infecting apis mellifera in India. Journal of Apicultural Science, 63(1), 165–170. https://doi.org/10.2478/jas-2019-0002
700. Shaw, J., Cutler, G. C., Manning, P., McCallum, R. S., & Astatkie, T. (2022). Does sending honey bee, Apis mellifera (Hymenoptera: Apidae), colonies to lowbush blueberry, Vaccinium angustifolium (Ericaceae), for pollination increase Nosema spp. (Nosematidae) spore loads? Canadian Entomologist, 154(1). https://doi.org/10.4039/tce.2022.30
701. Shirzadi, A., & Razmi, G. (2021). A microscopy and molecular studies of nosema ceranae infection in mazandaran province of Iran. Uludag Aricilik Dergisi, 21(2), 198–205. https://doi.org/10.31467/uluaricilik.991579
702. Shumkova, R., Balkanska, R., & Hristov, P. (2021). The herbal supplements nozemat herb® and nozemat herb plus®: An alternative therapy for n. Ceranae infection and its effects on honey bee strength and production traits. Pathogens, 10(2), 1–19. https://doi.org/10.3390/pathogens10020234
703. Shumkova, R., Georgieva, A., Radoslavov, G., Sirakova, D., Dzhebir, G., Neov, B., Bouga, M., & Hristov, P. (2018). The first report of the prevalence of Nosema ceranae in Bulgaria. PeerJ, 2018(1). https://doi.org/10.7717/peerj.4252
704. Shumkova, R., Neov, B., Georgieva, A., Teofanova, D., Radoslavov, G., & Hristov, P. (2020). Resistance of native honey bees from rhodope mountains and lowland regions of Bulgaria to Nosema ceranae and viral pathogens. Bulgarian Journal of Veterinary Medicine, 23(2), 206–217. https://doi.org/10.15547/bjvm.2201
705. Shutler, D., Head, K., Burgher-MacLellan, K. L., Colwell, M. J., Levitt, A. L., Ostiguy, N., & Williams, G. R. (2014). Honey bee Apis mellifera parasites in the absence of Nosema ceranae fungi and Varroa destructor mites. PLoS ONE, 9(6). https://doi.org/10.1371/journal.pone.0098599
706. Simeunovic, P., Stevanovic, J., Cirkovic, D., Radojicic, S., Lakic, N., Stanisic, L., & Stanimirovic, Z. (2014). Nosema ceranae and queen age influence the reproduction and productivity of the honey bee colony. Journal of Apicultural Research, 53(5), 545–554. https://doi.org/10.3896/IBRA.1.53.5.09
707. Sinpoo, C., Disayathanoowat, T., Williams, P. H., & Chantawannakul, P. (2019). Prevalence of infection by the microsporidian Nosema spp. In native

bumblebees (Bombus spp.) in northern Thailand. PLoS ONE, 14(3). https://doi.org/10.1371/journal.pone.0213171
708. Sinpoo, C., Paxton, R. J., Disayathanoowat, T., Krongdang, S., & Chantawannakul, P. (2018). Impact of Nosema ceranae and Nosema apis on individual worker bees of the two host species (Apis cerana and Apis mellifera) and regulation of host immune response. Journal of Insect Physiology, 105, 1–8. https://doi.org/10.1016/j.jinsphys.2017.12.010
709. Smart, M. D., & Sheppard, W. S. (2012). Nosema ceranae in age cohorts of the western honey bee (Apis mellifera). Journal of Invertebrate Pathology, 109(1), 148–151. https://doi.org/10.1016/j.jip.2011.09.009
710. Snow, J. W. (2016). A fluorescent method for visualization of Nosema infection in whole-mount honey bee tissues. Journal of Invertebrate Pathology, 135, 10–14. https://doi.org/10.1016/j.jip.2016.01.007
711. Snow, J. W. (2022). Nosema apis and N. ceranae Infection in Honey bees: A Model for Host-Pathogen Interactions in Insects. In Experientia supplementum (2012) (Vol. 114, pp. 153–177). https://doi.org/10.1007/978-3-030-93306-7_7
712. Snow, J. W., Ceylan Koydemir, H., Karinca, D. K., Liang, K., Tseng, D., & Ozcan, A. (2019). Rapid imaging, detection, and quantification of Nosema ceranae spores in honey bees using mobile phone-based fluorescence microscopy. Lab on a Chip, 19(5), 789–797. https://doi.org/10.1039/c8lc01342j
713. Sokół, R., & Michalczyk, M. (2012). Detection of Nosema spp. In worker bees of different ages during the flow season. Journal of Apicultural Science, 56(2), 19–25. https://doi.org/10.2478/v10289-012-0020-z
714. Sokół, R., & Michalczyk, M. (2016). Detection of Nosema spp. In worker bees, pollen and bee bread during the honey flow season. Acta Veterinaria Brno, 85(3), 261–266. https://doi.org/10.2754/avb201685030261
715. Sokół, R., Michalczyk, M., & Michołap, P. (2018). Preliminary studies on the occurrence of honeybee pathogens in the national bumblebee population. Annals of Parasitology, 64(4), 385–390. https://doi.org/10.17420/ap6404.175
716. Soroker, V., Hetzroni, A., Yakobson, B., David, D., David, A., Voet, H., Slabezki, Y., Efrat, H., Levski, S., Kamer, Y., Klinberg, E., Zioni, N., Inbar, S., & Chejanovsky, N. (2011). Evaluation of colony losses in Israel in relation to the incidence of pathogens and pests. Apidologie, 42(2), 192–199. https://doi.org/10.1051/apido/2010047
717. Stanimirović, Z., Glavinić, U., Ristanić, M., Jelisić, S., Vejnović, B., Niketić, M., & Stevanović, J. (2022). Diet Supplementation Helps Honey Bee Colonies in Combat Infections by Enhancing their Hygienic Behaviour. Acta Veterinaria, 72(2), 145–166. https://doi.org/10.2478/acve-2022-0013
718. Stanković, M., Bartolić, D., Mutavdžić, D., Marković, S., Grubić, S., Jovanović, N. M., & Radotić, K. (2023). Estimation of honey bee colony infection with Nosema ceranae and Varroa destructor using fluorescence spectroscopy in combination with differential scanning calorimetry of honey samples. Journal of Apicultural Research, 62(3), 507–513. https://doi.org/10.1080/00218839.2021.1889803
719. Stevanovic, J., Schwarz, R. S., Vejnovic, B., Evans, J. D., Irwin, R. E., Glavinic, U., & Stanimirovic, Z. (2016). Species-specific diagnostics of Apis mellifera trypanosomatids: A nine-year survey (2007–2015) for trypanosomatids and

microsporidians in Serbian honey bees. Journal of Invertebrate Pathology, 139, 6–11. https://doi.org/10.1016/j.jip.2016.07.001
720. Stevanovic, J., Simeunovic, P., Gajic, B., Lakic, N., Radovic, D., Fries, I., & Stanimirovic, Z. (2013). Characteristics of Nosema ceranae infection in Serbian honey bee colonies. Apidologie, 44(5), 522–536. https://doi.org/10.1007/s13592-013-0203-z
721. Stevanovic, J., Stanimirovic, Z., Genersch, E., Kovacevic, S. R., Ljubenkovic, J., Radakovic, M., & Aleksic, N. (2011). Dominance of Nosema ceranae in honey bees in the Balkan countries in the absence of symptoms of colony collapse disorder. Apidologie, 42(1), 49–58. https://doi.org/10.1051/apido/2010034
722. Stojanov, I., Ratajac, R., Radulović, J. P., Petrović, J., Rašović, M. B., & Pušić, I. (2021). CONTROL AND VIABILITY OF BEE NOSEMOSES. Archives of Veterinary Medicine, 14(2), 119–129. https://doi.org/10.46784/eavm.v14i2.288
723. Straw, E. A., & Brown, M. J. F. (2021). No evidence of effects or interaction between the widely used herbicide, glyphosate, and a common parasite in bumble bees. PeerJ, 9. https://doi.org/10.7717/peerj.12486
724. Sun, J., Qin, F., Sun, F., He, P., Wei, E., Wang, R., Zhu, F., Wang, Q., Tang, X., Zhang, Y., & Shen, Z. (2023). Identification and subcellular colocalization of protein transport protein Sec61α and Sec61γ in Nosema bombycis. Gene, 851. https://doi.org/10.1016/j.gene.2022.146971
725. Sun, J., Zhu, F., Chen, H., Yao, M., Zhu, G., Zhang, Y., Wang, Q., & Shen, Z. (2020). Identification and subcellular localisation of hexokinase-2 in Nosema bombycis. Folia Parasitologica, 67, 1–8. https://doi.org/10.14411/FP.2020.023
726. Sun, M., Qian, J., Li, Q., Zhang, T., Zhang, J., Zhao, H., Zhang, K., Zhu, L., Chen, T., Chen, D., Fu, Z., & Guo, R. (2023). Cloning, molecular characteristics and expression pattern of transmethylase-like protein 5 gene NcMettl5 in Nosema ceranae. Mycosystema, 42(5), 1102–1113. https://doi.org/10.13346/j.mycosystema.220278
727. Suraporn, S., Natsopoulou, M. E., Doublet, V., McMahon, D. P., & Paxton, R. J. (2013). Nosema ceranae is not detected in honey bees (Apis spp.) of northeast Thailand. Journal of Apicultural Research, 52(5), 259–261. https://doi.org/10.3896/IBRA.1.52.5.13
728. Suwannapong, G., Maksong, S., Phainchajoen, M., Benbow, M. E., & Mayack, C. (2018). Survival and health improvement of Nosema infected Apis florea (Hymenoptera: Apidae) bees after treatment with propolis extract. Journal of Asia-Pacific Entomology, 21(2), 437–444. https://doi.org/10.1016/j.aspen.2018.02.006
729. Suwannapong, G., Maksong, S., Seanbualuang, P., & Benbow, M. E. (2010). Experimental infection of red dwarf honeybee, Apis florea, with Nosema ceranae. Journal of Asia-Pacific Entomology, 13(4), 361–364. https://doi.org/10.1016/j.aspen.2010.07.003
730. Suwannapong, G., Yemor, T., Boonpakdee, C., & Benbow, M. E. (2011). Nosema ceranae, a new parasite in Thai honeybees. Journal of Invertebrate Pathology, 106(2), 236–241. https://doi.org/10.1016/j.jip.2010.10.003
731. Syromyatnikov, M. Y., Savinkova, O. V., Panevina, A. V., Solodskikh, S. A., Lopatin, A. V., & Popov, V. N. (2019). Quality Control of Bee-Collected Pollen Using Bumblebee Microcolonies and Molecular Approaches Reveals No Correlation between Pollen Quality and Pathogen Presence. Journal of Economic Entomology, 112(1), 49–59. https://doi.org/10.1093/jee/toy345

781. Vavilova, V. Y., Konopatskaia, I., Luzyanin, S. L., Woyciechowski, M., & Blinov, A. G. (2017). Parasites of the genus Nosema, Crithidia and Lotmaria in the honeybee and bumblebee populations: A case study in India. Vavilovskii Zhurnal Genetiki i Selektsii, 21(8), 943–951. https://doi.org/10.18699/VJ17.317
782. Vejnovic, B., Stevanovic, J., Schwarz, R. S., Aleksic, N., Mirilovic, M., Jovanovic, N. M., & Stanimirovic, Z. (2018). Quantitative PCR assessment of Lotmaria passim in Apis mellifera colonies co-infected naturally with Nosema ceranae. Journal of Invertebrate Pathology, 151, 76–81. https://doi.org/10.1016/j.jip.2017.11.003
783. Villa, J. D., Bourgeois, A. L., & Danka, R. G. (2013). Negative evidence for effects of genetic origin of bees on Nosema ceranae, positive evidence for effects of Nosema ceranae on bees. Apidologie, 44(5), 511–518. https://doi.org/10.1007/s13592-013-0201-1
784. Wagoner, K. M., Boncristiani, H. F., & Rueppell, O. (2013). Multifaceted responses to two major parasites in the honey bee (Apis mellifera). BMC Ecology, 13. https://doi.org/10.1186/1472-6785-13-26
785. Wang, Q., Dai, P., Guzman-Novoa, E., Wu, Y., Hou, C., & Diao, Q. (2019). Nosema ceranae, the most common microsporidium infecting Apis mellifera in the main beekeeping regions of China since at least 2005. Journal of Apicultural Research, 58(4), 562–566. https://doi.org/10.1080/00218839.2019.1632148
786. Wang, Y., Geng, H., Dang, X., Xiang, H., Li, T., Pan, G., & Zhou, Z. (2017). Comparative Analysis of the Proteins with Tandem Repeats from 8 Microsporidia and Characterization of a Novel Endospore Wall Protein Colocalizing with Polar Tube from Nosema bombycis. Journal of Eukaryotic Microbiology, 64(5), 707–715. https://doi.org/10.1111/jeu.12412
787. Wang, Z., Wang, S., Fan, X., Zhang, K., Zhang, J., Zhao, H., Gao, X., Zhang, Y., Guo, S., Zhou, D., Li, Q., Na, Z., Chen, D., & Guo, R. (2023). Systematic Characterization and Regulatory Role of lncRNAs in Asian Honey Bees Responding to Microsporidian Infestation. International Journal of Molecular Sciences, 24(6). https://doi.org/10.3390/ijms24065886
788. Wei, X., Evans, J. D., Chen, Y., & Huang, Q. (2022). Spillover and genome selection of the gut parasite Nosema ceranae between honey bee species. Frontiers in Cellular and Infection Microbiology, 12. https://doi.org/10.3389/fcimb.2022.1026154
789. Wei, X., Zheng, J., Evans, J. D., & Huang, Q. (2022). Transgenerational genomic analyses reveal allelic oscillation and purifying selection in a gut parasite Nosema ceranae. Frontiers in Microbiology, 13. https://doi.org/10.3389/fmicb.2022.927892
790. Wells, T., Wolf, S., Nicholls, E., Groll, H., Lim, K. S., Clark, S. J., Swain, J., Osborne, J. L., & Haughton, A. J. (2016). Flight performance of actively foraging honey bees is reduced by a common pathogen. Environmental Microbiology Reports, 8(5), 728–737. https://doi.org/10.1111/1758-2229.12434
791. Whitaker, J., Szalanski, A. L., & Kence, M. (2011). Molecular detection of Nosema ceranae and N. apis from Turkish honey bees. Apidologie, 42(2), 174–180. https://doi.org/10.1051/apido/2010045
792. Williams, G. R., Sampson, M. A., Shutler, D., & Rogers, R. E. L. (2008). Does fumagillin control the recently detected invasive parasite Nosema ceranae in western

honey bees (Apis mellifera)? Journal of Invertebrate Pathology, 99(3), 342–344. https://doi.org/10.1016/j.jip.2008.04.005
793. Williams, G. R., Shafer, A. B. A., Rogers, R. E. L., Shutler, D., & Stewart, D. T. (2008). First detection of Nosema ceranae, a microsporidian parasite of European honey bees (Apis mellifera), in Canada and central USA. Journal of Invertebrate Pathology, 97(2), 189–192. https://doi.org/10.1016/j.jip.2007.08.005
794. Williams, G. R., Shutler, D., Little, C. M., Burgher-Maclellan, K. L., & Rogers, R. E. L. (2011). The microsporidian Nosema ceranae, the antibiotic Fumagilin-B®, and western honey bee (Apis mellifera) colony strength. Apidologie, 42(1), 15–22. https://doi.org/10.1051/apido/2010030
795. Williams, G. R., Shutler, D., & Rogers, R. E. L. (2010). Effects at Nearctic north-temperate latitudes of indoor versus outdoor overwintering on the microsporidium Nosema ceranae and western honey bees (Apis mellifera). Journal of Invertebrate Pathology, 104(1), 4–7. https://doi.org/10.1016/j.jip.2010.01.009
796. Williams, M.-K. F., Cleary, D. A., Tripodi, A. D., & Szalanski, A. L. (2021). Co-occurrence of Lotmaria passim and Nosema ceranae in honey bees (Apis mellifera L.) from six states in the United States. Journal of Apicultural Research. https://doi.org/10.1080/00218839.2021.1960745
797. Wojcik, A., & Chorbinski, P. (2014). Nosema ceranae, a widespread pathogen of honey bees (Apis mellifera). MEDYCYNA WETERYNARYJNA-VETERINARY MEDICINE-SCIENCE AND PRACTICE, 70(12), 735–739.
798. Wolf, S., McMahon, D. P., Lim, K. S., Pull, C. D., Clark, S. J., Paxton, R. J., & Osborne, J. L. (2014). So near and yet so far: Harmonic radar reveals reduced homing ability of nosema infected honeybees. PLoS ONE, 9(8). https://doi.org/10.1371/journal.pone.0103989
799. Wu, J. Y., Smart, M. D., Anelli, C. M., & Sheppard, W. S. (2012). Honey bees (Apis mellifera) reared in brood combs containing high levels of pesticide residues exhibit increased susceptibility to Nosema (Microsporidia) infection. Journal of Invertebrate Pathology, 109(3), 326–329. https://doi.org/10.1016/j.jip.2012.01.005
800. Wu, Y., Ye, Y., Zhang, J., Qian, J., Zhang, W., Yu, K., Ji, T., Lin, Z., Zhao, H., Chen, D., & Guo, R. (2022). Expression profiles of nce-miR-12220 and its target genes during the Nosema ceranae infection process of Apis mellifera ligustica workers. Mycosystema, 41(10), 1546–1557. https://doi.org/10.13346/j.mycosystema.220026
801. Wu, Y., Zheng, Y., Chen, Y., Chen, G., Zheng, H., & Hu, F. (2020). Apis cerana gut microbiota contribute to host health though stimulating host immune system and strengthening host resistance to Nosema ceranae. Royal Society Open Science, 7(5). https://doi.org/10.1098/rsos.192100
802. Xing, W., Zhou, D., Long, Q., Sun, M., Guo, R., & Wang, L. (2021). Immune response of eastern honeybee worker to nosema ceranae infection revealed by transcriptomic investigation. Insects, 12(8). https://doi.org/10.3390/insects12080728
803. Xiong, X., Geden, C. J., Bergstralh, D. T., White, R. L., Werren, J. H., & Wang, X. (2023). New insights into the genome and transmission of the microsporidian pathogen Nosema muscidifuracis. Frontiers in Microbiology, 14. https://doi.org/10.3389/fmicb.2023.1152586
804. Xu, J., He, Q., Ma, Z., Li, T., Zhang, X., Debrunner-Vossbrinck, B. A., Zhou, Z., & Vossbrinck, C. R. (2016). The genome of nosema sp. Isolate YNPr: A

comparative analysis of genome evolution within the nosema/vairimorpha clade. PLoS ONE, 11(9). https://doi.org/10.1371/journal.pone.0162336
805. Yamandú, M., Jorge, H., Juan, C., Helena, K., Gustavo, R., Sebastián, D. C., & Ciro, I. (2013). Control of Nosema ceranae in Honey Bees (A pis mellifera) Colonies in Eucalyptus grandis Plantations. AGROCIENCIA-URUGUAY, 17(1), 108–113.
806. Yang, B., Peng, G., Li, T., & Kadowaki, T. (2013). Molecular and phylogenetic characterization of honey bee viruses, Nosema microsporidia, protozoan parasites, and parasitic mites in China. Ecology and Evolution, 3(2), 298–311. https://doi.org/10.1002/ece3.464
807. Yang, D., Dang, X., Peng, P., Long, M., Ma, C., Qin, J. J. G., Wu, H., Liu, T., Zhou, X., Pan, G., & Zhou, Z. (2014). NbHSWP11, a microsporidia nosema bombycis protein, localizing in the spore wall and membranes, reduces spore Adherence to Host Cell BME. Journal of Parasitology, 100(5), 623–632. https://doi.org/10.1645/13-286.1
808. Yang, D., Xu, X., Zhao, H., Yang, S., Wang, X., Zhao, D., Diao, Q., & Hou, C. (2018). Diverse factors affecting efficiency of RNAi in honey bee viruses. Frontiers in Genetics, 9(SEP). https://doi.org/10.3389/fgene.2018.00384
809. Yemor, T., Phiancharoen, M., Eric Benbow, M., & Suwannapong, G. (2015). Effects of stingless bee propolis on Nosema ceranae infected Asian honey bees, Apis cerana. Journal of Apicultural Research, 54(5), 468–473. https://doi.org/10.1080/00218839.2016.1162447
810. Yi, M., Lü, Q., Liu, K., Wang, L., Wu, Y., Zhou, Z., & Long, M. (2019). Expression, Purification and Localization Analysis of Polar Tube Protein 2 (NbPTP2) from Nosema bombycis. Scientia Agricultura Sinica, 52(10), 1830–1838. https://doi.org/10.3864/j.issn.0578-1752.2019.10.015
811. Yoshiyama, M., & Kimura, K. (2011). Distribution of Nosema ceranae in the European honeybee, Apis mellifera in Japan. Journal of Invertebrate Pathology, 106(2), 263–267. https://doi.org/10.1016/j.jip.2010.10.010
812. Zanet, S., Battisti, E., Alciati, R., Trisciuoglio, A., Cauda, C., & Ferroglio, E. (2019). Nosema ceranae contamination in bee keeping material: The use of ozone as disinfection method. Journal of Apicultural Research, 58(1), 62–66. https://doi.org/10.1080/00218839.2018.1517989
813. Zbrozek, M., Fearon, M. L., Weise, C., & Tibbetts, E. A. (2023). Honeybee visitation to shared flowers increases Vairimorpha ceranae prevalence in bumblebees. Ecology and Evolution, 13(9). https://doi.org/10.1002/ece3.10528
814. Zerek, A., Yaman, M., & Dik, B. (2022). Prevalence of nosemosis in honey bees (Apis mellifera L., 1758) of the Hatay province in Turkey. Journal of Apicultural Research, 61(3), 368–374. https://doi.org/10.1080/00218839.2021.2008706
815. Zhou, D. D., Shi, X. Y., Wang, J., Fan, Y. C., Zhu, Z. W., Jiang, H. B., Fan, X. X., Xiong, C. L., Zheng, Y. Z., Fu, Z. M., Xu, G. J., Chen, D. F., & Guo, R. (2020). Investigation of competing endogenous RNA regulatory network and putative function of long non-coding RNAs in nosema ceranae spore. Scientia Agricultura Sinica, 53(10), 2122–2136. https://doi.org/10.3864/j.issn.0578-1752.2020.10.018
816. Zhu, F., Shen, Z., Xu, L., & Guo, X. (2013). Molecular characteristics of the alpha- and beta-tubulin genes of Nosema philosamiae. Folia Parasitologica, 60(5), 411–415. https://doi.org/10.14411/fp.2013.043
817. Zhu, X., Zhou, S., & Huang, Z. Y. (2014). Transportation and pollination service increase abundance and prevalence of Nosema ceranae in honey bees (Apis

mellifera). Journal of Apicultural Research, 53(4), 469–471. https://doi.org/10.3896/IBRA.1.53.4.06
818. Zhu, Z., Wang, J., Fan, X., Long, Q., Chen, H., Ye, Y., Zhang, K., Ren, Z., Zhang, Y., Niu, Q., Chen, D., & Guo, R. (2022). CircRNA-regulated immune responses of asian honey bee workers to microsporidian infection. Frontiers in Genetics, 13. https://doi.org/10.3389/fgene.2022.1013239
819. Zinatullina, Z. Y., Dolnikova, T. Y., Domatskaya, T. F., & Domatsky, A. N. (2018). Monitoring diseases of honey bees (Apis mellifera) in Russia. UKRAINIAN JOURNAL OF ECOLOGY, 8(3), 106–112.
